# Carbohydrate-active enzyme annotation in microbiomes using dbCAN

**DOI:** 10.1101/2024.01.10.575125

**Authors:** Jinfang Zheng, Le Huang, Haidong Yi, Yuchen Yan, Xinpeng Zhang, Jerry Akresi, Yanbin Yin

**Author notes:** To whom correspondence Yanbin Yin.

## Abstract

CAZymes or carbohydrate-active enzymes are critically important for human gut health, lignocellulose degradation, global carbon recycling, soil health, and plant disease. We developed dbCAN as a web server in 2012 and actively maintain it for automated CAZyme annotation. Considering data privacy and scalability, we provide run_dbcan as a standalone software package since 2018 to allow users perform more secure and scalable CAZyme annotation on their local servers. Here, we offer a comprehensive computational protocol on automated CAZyme annotation of microbiome sequencing data, covering everything from short read pre-processing to data visualization of CAZyme and glycan substrate occurrence and abundance in multiple samples. Using a real-world metagenomic sequencing dataset, this protocol describes commands for dataset and software preparation, metagenome assembly, gene prediction, CAZyme prediction, CAZyme gene cluster (CGC) prediction, glycan substrate prediction, and data visualization. The expected results include publication-quality plots for the abundance of CAZymes, CGCs, and substrates from multiple CAZyme annotation routes (individual sample assembly, co-assembly, and assembly-free). For the individual sample assembly route, this protocol takes ∼33h on a Linux computer with 40 CPUs, while other routes will be faster. This protocol does not require programming experience from users, but it does assume a familiarity with the Linux command-line interface and the ability to run Python scripts in the terminal. The target audience includes the tens of thousands of microbiome researchers who routinely use our web server. This protocol will encourage them to perform more secure, rapid, and scalable CAZyme annotation on their local computer servers.

## Introduction

CAZymes are short for carbohydrate-active enzymes, including six classes of enzymes and functional modules: glycosyltransferases [GTs], glycoside hydrolases [GHs], polysaccharide lyases [PLs], carbohydrate esterases [CEs], enzymes of auxiliary activities [AAs], and carbohydrate binding modules (CBMs)^1^. According to the expert curated CAZy database^1-3^ (www.cazy.org), CAZymes are found in all kinds of cellular organisms (bacteria, archaea, fungi, plants, algae, animals, protists) and viruses, but are most abundant in plant cell wall degrading microorganisms, e.g., 11.7% of genes in the human gut *Bacteroides cellulosilyticus* encode CAZymes. CAZymes have demonstrated significance in human gut health^4^, lignocellulose degradation^5^, soil health^6^, global carbon recycling^7^, and plant diseases^8^.

### dbCAN web server

Given the importance of automated CAZyme annotation in microbial genomes and metagenomes, we developed dbCAN as a web server in 2012^9^ (**Fig. 1**). The core component of this web server is a database of profile hidden Markov models (HMMs) that were built for CAZyme family-specific functional domains and modules (**Table 1**). This HMM database is searched with user submitted protein sequences as query and HMMER as the search tool^10^. The dbCAN web server has been updated twice in the last decade, dbCAN2 in 2018^11^ and dbCAN3 in 2023^12^ (**Fig. 1**), and now become the most widely used automated CAZyme annotation tool. Its popularity is reflected by the high citation number (>3,200) of our web server papers^9,11,12^ by the microbiome, gastroenterology, biotechnology, marine science, environment, agriculture, nutrition, and bioenergy research communities^7,13-23^. The web server processes over 40,000 user submitted jobs in the year 2023 from over 150 countries (https://clustrmaps.com/site/19yt1). The underlying HMM database of dbCAN is also integrated into over a dozen of other bioinformatics software systems, such as the App Catalog of the Department of Energy’s Systems Biology Knowledgebase (KBase)^24^, the microbiome and virome metabolism annotation tools DRAM^25^ and METABOLIC^26^, and the CAZyme functional specificity analysis tool SACCHARIS^27^.

**Table 1.** Features and functions in the three versions of dbCAN.

| Features | Functions | dbCAN | dbCAN2 | dbCAN3 |
| --- | --- | --- | --- | --- |
| HMMER vs CAZyme domain HMMdb | CAZyme family annotation | Yes | Yes | Yes |
| HMMER vs subfamily HMMdb | CAZyme subfamily annotation |  |  | Yes |
| DIAMOND vs CAZy | CAZyme family annotation |  | Yes | Yes |
| Hotpep vs PPR | CAZyme family annotation |  | Yes | Yes |
| CGC-finder | CAZyme gene cluster prediction |  | Yes | Yes |
| CAZyme subfamilies mapped to EC and glycans (via dbCAN-sub) | Substrate prediction for CAZymes |  |  | Yes |
| Majority voting on CAZymes (substrates from dbCAN-sub) in CGCs | Substrate prediction for CGCs |  |  | Yes |
| BLAST vs PULs in dbCAN-PUL | Substrate prediction for CGCs |  |  | Yes |

**Fig. 1:**
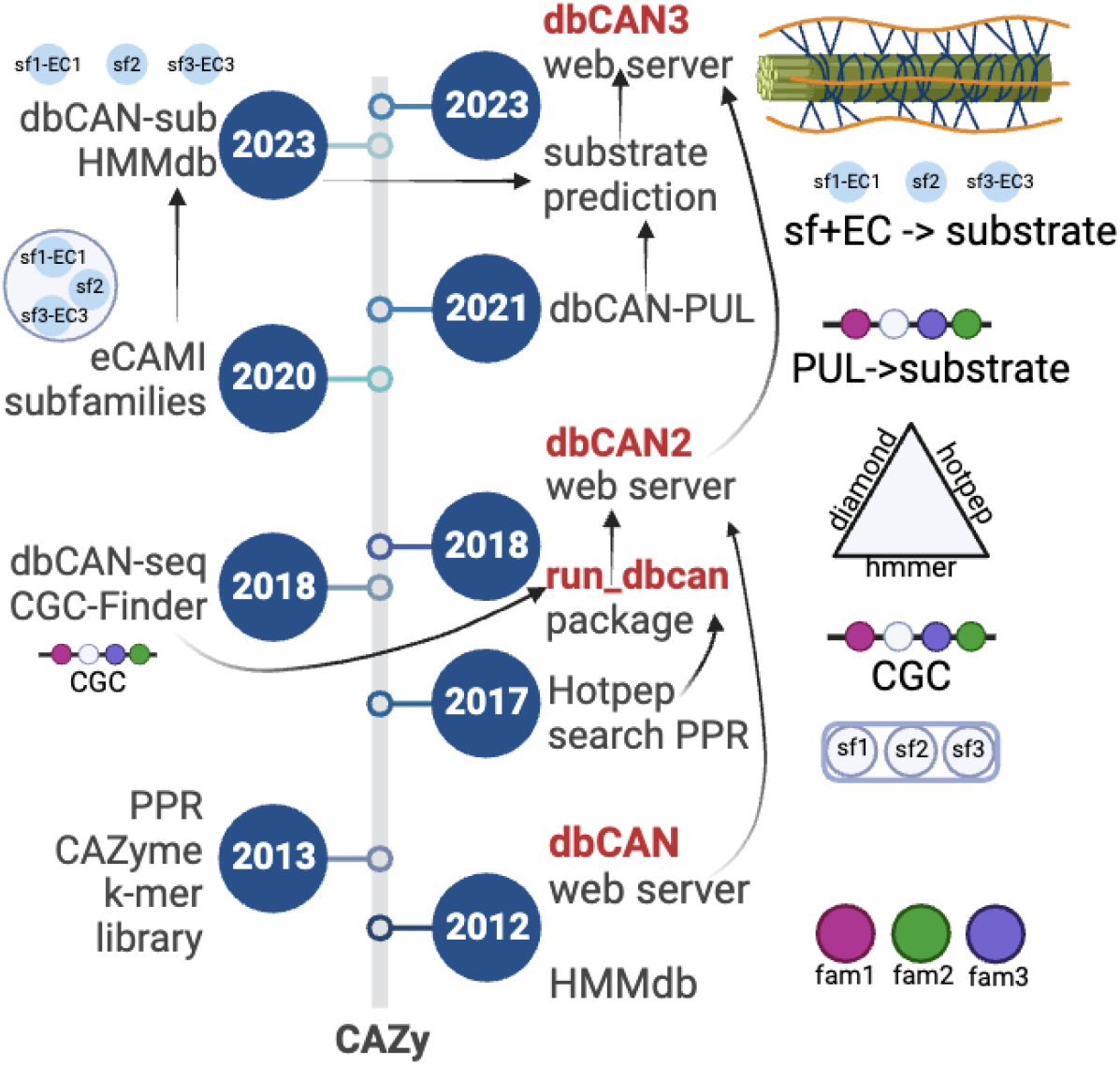
Brief history of dbCAN development. dbCAN was developed in 2012 based on the expert curated CAZy database^1-3^. CAZy was established by Henrissat B in 1990s as the CAZyme classification database that defines over 470 CAZyme families (33 families are further classified into over 300 subfamilies) according to biochemically characterized proteins. dbCAN was a HMMdb for all CAZy families/subfamilies and a web server that enabled automated CAZyme annotation at the genome scale. It had two major updates in 2018 and 2023 by incorporating new functions (**Table 1**) developed by us and others (detailed in the main text). A standalone software package run_dbcan was released in 2018 on GitHub that allows users download, install, and run CAZyme annotation on their local computers. The graphics illustrate the main functions/milestones developed in each tool. “fam” are families and “sf” are subfamilies. CGCs are CAZyme gene clusters.

### Improved dbCAN functions

In addition to the HMM database, new functions were added into dbCAN2 and dbCAN3 (**Table 1**). In 2018, we developed CGC-finder^28^ and integrated it into dbCAN2^11^ for predicting CAZyme gene clusters (CGCs) (**Fig. 1**). CGCs are defined as physically linked gene clusters in bacterial genomes that contain four different classes of signature genes: (i) CAZymes, (ii) signaling transduction proteins (STPs), (iii) transporters (TCs), and (iv) transcription factors (TFs)^28^. Protein products of these genes are often co-expressed and work synergistically to fully degrade various complex carbohydrates^29^. In the literature biochemically characterized CGCs are termed polysaccharide utilization loci (PULs)^30^ indicating the specific glycan substrates, e.g., starch utilization loci, xylan utilization loci, or mucin utilization loci^31^. PULDB was published in 2018 by the CAZy team to collect experimentally verified PULs and their homologous gene clusters in genomes of the *Bacteroidetes* phylum^32^. However, PULs in PULDB must have the tandem *susC* and *susD* gene pair, which are absent in Gram-positive PULs^30,33^. A standalone tool PULpy^34^ is available to predict these *susCD-*containing CGCs in genomes of Gram-negative bacteria, implementing an algorithm used by PULDB^35^.

We published dbCAN-PUL in 2021 to cover experimentally verified PULs from various bacterial phyla with characterized glycan substrates including Gram-positive PULs^36^. The co-presence of different CAZyme genes in the same PUL/CGC is a highly useful approach to assist the discovery of new CAZyme families that do not share sequence similarity with established CAZyme families in the CAZy database. For example, Ndeh D *et al*. characterized six new pectin degrading GH families in a pectin utilization loci of *Bacteroides thetaiotaomicron* genome, where the pectin PUL also contains other pectin degrading CAZymes^37^. Aside from CGC-finder, dbCAN2 also integrated DIAMOND^38^ search against CAZy annotated protein sequences and an amino acid k-mer-based search tool Hotpep^39^ against the CAZyme signature k-mer peptide library PPR^40^ into the annotation pipeline (**Fig. 1**). Hotpep and PPR were later replaced by eCAMI^41^ in 2022.

In 2023, dbCAN2 was further updated to dbCAN3^12^ (**Fig. 1**), featuring glycan substrate prediction for both CAZymes and CGCs (**Table 1**). This substrate prediction function is critical to microbiome researchers who are interested in knowing not only what CAZyme families exist in their genomes or metagenomes, but also what glycan substrates can be metabolized in their samples (e.g., by bacterial isolates or microbiota)^42-44^. For example, a paper published in 2018 compared human gut microbiomes of two groups of patients with type 2 diabetes^45^. Using dbCAN, they mined the metagenomic sequencing data of the microbiomes for CAZyme families, and further manually curated the CAZyme families to find those targeting various dietary fibers including starch, inulin, pectin, and mucin. Their conclusion was that patients given dietary fibers benefited from their microbiome utilization of these fibers leading to alleviated disease symptoms. Another paper in 2021 used dbCAN to mine metagenomic sequencing data for CAZymes from Native American palaeofaeces samples (1,000-2,000 years ago)^46^, as well as gut microbiome samples of industrialized and non-industrialized modern humans. They manually annotated and grouped the CAZyme families into different glycan substrate groups (e.g., cellulose, chitin, hemicellulose, pectin, starch/glycogen, mucin). Evidence was shown that the human palaeofaeces samples are enriched with CAZyme families targeting starch/glycogen and chitin, and share more similar CAZyme enrichment patterns with non-industrialized samples than industrialized samples.

To develop a glycan substrate prediction function in dbCAN, we first classified 426 CAZy defined families into over 26,000 subfamilies using eCAMI^41^ and created HMMs for subfamilies (named dbCAN-sub, **Fig. 1**). We further manually curated the CAZy webpages and literature to map CAZyme subfamilies to EC (enzyme commission) numbers and further to glycan substrates (**Table 1**). Therefore, users can now perform glycan substrate prediction for CAZymes through dbCAN-sub search and substrate mapping. To enable substrate prediction for CGCs, we included two approaches in dbCAN3. The first one is BLAST sequence similarity search of CGCs against dbCAN-PUL^36^, and the second is a majority voting on substrates predicted for CAZymes (via dbCAN-sub) within CGCs. Our benchmark experiments showed that the dbCAN-PUL search approach has a better performance than the majority voting approach^12^.

### Standalone software package run_dbcan

Along with dbCAN2 published in 2018, we also released a standalone Python software package named run_dbcan (**Fig. 1**), which can be installed and run locally by users for automated CAZyme annotation. The standalone package was important for two reasons. The first is the data privacy: users may not want to upload their unpublished data to a public web server, and prefer to perform CAZyme annotation on their local private servers. The second is the data scalability: with lower cost DNA sequencing, researchers can now routinely perform large scale and multi-sample microbiome sequencing and generate gigabase (GB) or terabase (TB) amount of data, which is not possible to use our web server for CAZyme annotation. The run_dbcan package is freely available on GitHub (https://github.com/linnabrown/run_dbcan) and can be installed using Bioconda (https://anaconda.org/bioconda/run-dbcan) or Docker (https://hub.docker.com/r/haidyi/run_dbcan). This has allowed users perform more secure and scalable CAZyme annotation of large microbiome sequencing data on their local servers (single server or computing cluster). According to the record on Anaconda, the run_dbcan bioconda package has been downloaded over 6,000 times in the last three years. The run_dbcan package is updated following the major updates of the dbCAN web server. Therefore, the current run_dbcan release also incorporates the substrate prediction function for CAZymes and CGCs.

In addition to dbCAN, there are other tools that can also perform automated CAZyme annotation locally or on the web. However, they are either offline (e.g., CAZymes Analysis Toolkit, CAT^46^), no longer maintained (e.g., PPR^40^ and Hotpep^39^), or have limited functions (**Table 1**) compared to dbCAN (e.g., CUPP^47^). Specifically, no other software packages like run_dbcan can perform not only CAZyme annotation but also CGC identification and glycan substrate prediction. Even for the single CAZyme annotation function, an independent group showed that dbCAN/run_dbcan has the best accuracy^48^ compared to Hotpep, eCAMI, and CUPP.

### Motivation for this protocol

We provide detailed documentation on GitHub for run_dbcan users to learn how to install and run the program. However, given the varying levels of users’ bioinformatics and microbiome data analysis skills, we frequently receive emails and messages on GitHub regarding various questions and issues ranging from software installation, data pre-processing, job running, parameter setting, and result interpretation. This motivates us to develop this protocol. The targeted audience includes tens of thousands of dbCAN users and microbiome researchers, especially early career scientists (e.g., graduate students and postdocs), who would like to use our standalone run_dbcan software package to process microbiome sequencing reads of large sizes (e.g., over 100GB) on their local servers.

Additionally, dbCAN and run_dbcan require assembled contigs as input. However, in practice microbiome researchers usually start with raw sequencing (metagenomic or metatranscriptomic) reads from multiple samples, which must be pre-processed and assembled before CAZyme annotation. Users often also want CAZyme abundance comparison and data visualization across multiple microbiome samples. Hence, this protocol paper will offer a comprehensive tutorial on CAZyme annotation, covering everything from the input sequencing reads to data visualization of CAZyme occurrence and abundance in multiple samples. It will include instructions on software preparation, read pre-processing, metagenome assembly, gene prediction, CAZyme prediction, CGC prediction, glycan substrate prediction, and data visualization (**Fig. 1**).

### Overview of the protocol

In this protocol, we will use a real-world microbiome dataset as an example to demonstrate the computational workflow for automated CAZyme and glycan substrate annotation (occurrence and abundance) in microbiomes. A typical workflow is illustrated in **Fig. 2**. Firstly, raw sequencing reads are pre-processed including the removal of contamination, adapters, and low-quality read trimming to obtain the clean reads by running trim_galore^49^ and Kraken2^50^ (steps1-2). Next, the clean reads of each sample are assembled into contigs by MEGAHIT^51^ (step3). The contigs are passed to Prokka^52^ to annotate the gene models (step4). In step5, the contigs are annotated for CAZymes and CGCs by run_dbcan using the protein sequence file (faa) and gene annotation file (gff) produced by Prokka. Step6 produces a list of annotated CAZymes and CGCs and their location information in contigs. The substrate prediction function in run_dbcan infers the glycan substrates for CAZyme and CGC in step7. To calculate the abundance of the CAZyme, substrate and CGC (steps8-14), the clean reads from step2 are mapped to the nucleotide coding sequences (CDS) of proteins from step4. To visualize the occurrence and abundance results (steps15-20), Python scripts are provided to make publication-quality plots, which can be saved in PDF format.

**Fig. 2:**
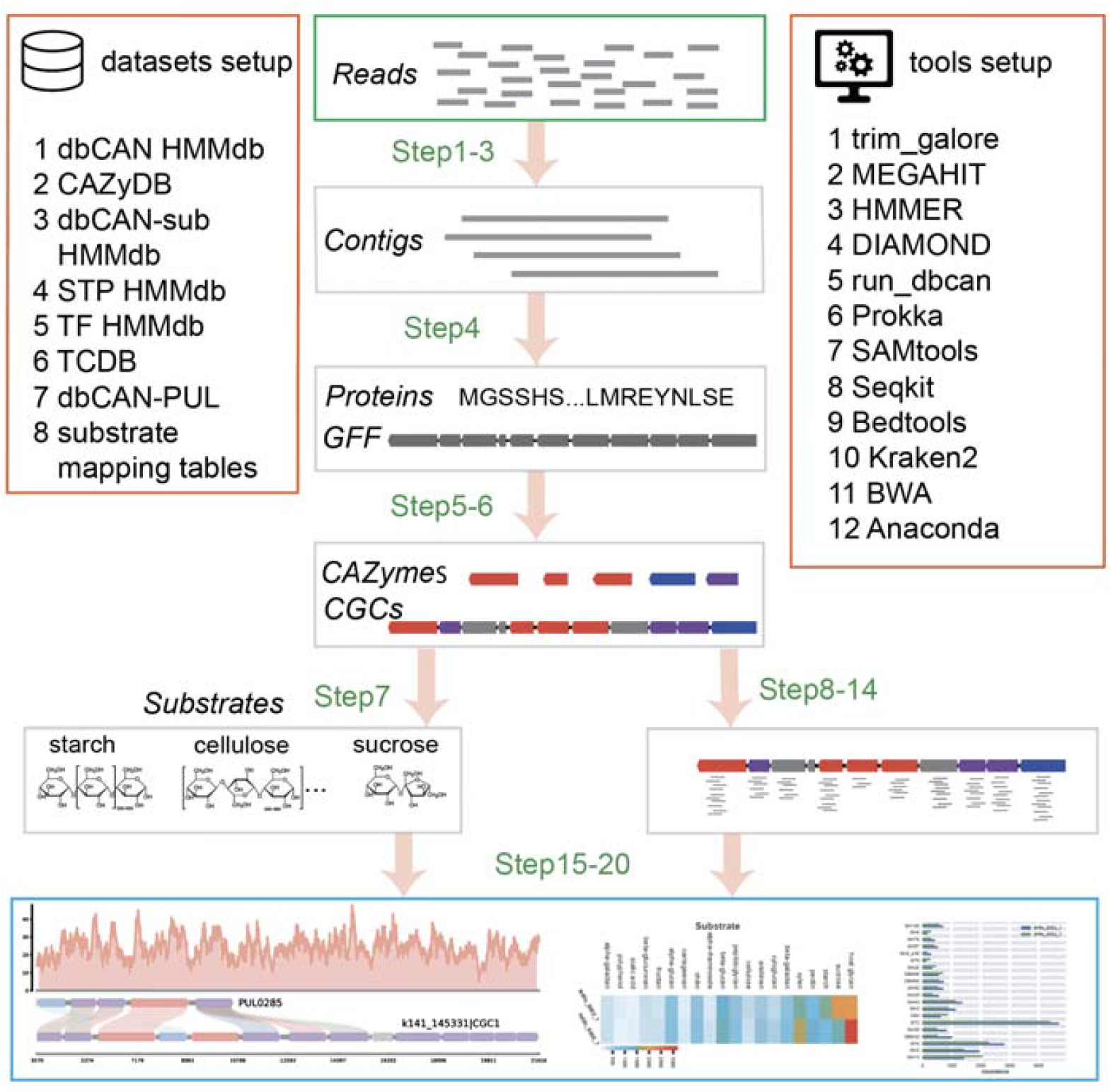
Overview of the protocol. Raw reads undergo essential pre-processing steps, encompassing de-contamination, low-quality read trimming, and assembly (steps 1-3). Subsequently, proteins are predicted from contigs using Prokka (step 4). CAZymes, CGCs, and glycan substrates are predicted by run_dbcan (steps 5-7). Abundances of CAZyme, CGC, and substrates in the samples are calculated from read mapping results (step8-14). Finally, results will be subject to data visualization, encompassing bar plots, heatmaps, and read count plots (step15-20).

This protocol does not require programming experience from the users, but it does assume a familiarity with the Linux command-line interface and the ability to run Python scripts in the terminal. Users should also be able to edit Linux commands and plain-text scripts in the command-line terminal.

### Limitations

While dbCAN and run_dbcan can annotate CAZymes in both prokaryotes and eukaryotes, the prediction of CGCs only works for prokaryotes. This is because the clustering of CAZymes and other associated genes (STPs, TCs, and TFs) to form CGC/PUL has only been verified in prokaryotic genomes. It is unknown whether the CGC/PUL concept also applies to eukaryotes. Even it does, due to the presence of introns and large intergenic regions, the CGC definition in eukaryotes will have to be carefully revised via experimental and computational evaluations. Therefore, the current protocol is limited to CAZyme annotation in microbiome sequencing data of prokaryotes. However, users can still perform CAZyme annotation and substrate prediction for CAZymes in eukaryotes given protein sequences as the input.

### Experimental design

#### Dataset description

We selected a real-world microbiome dataset for this protocol. This dataset (named Carter2023) was published in 2023 from an ultra-deep sequencing of 351 fecal microbiome samples of Hadza hunter-gatherers^16^. The goal of the study was to identify bacterial species in the human gut that were lost during modern industrialization. The microbiomes of hunter-gatherers were found to be strongly affected by their seasonal dietary cycling: more meat and grain consumption in dry seasons, and more berry and honey consumption in wet seasons. The CAZyme annotation of microbiome sequencing data confirmed their earlier paper published in 2017^53^ that the CAZyme abundance and diversity were higher in fecal microbiome samples of dry seasons than wet seasons. For this protocol, we will select two samples (paired-end 2x140bp reads) of a same human individual from a bush camp of Tanzania (late wet season vs late dry season of 2014) for CAZyme annotation (**Table 2**). All the data are publicly available in the SRA database and the original paper^16^ has already addressed the data privacy and permission issues.

**Table 2.**
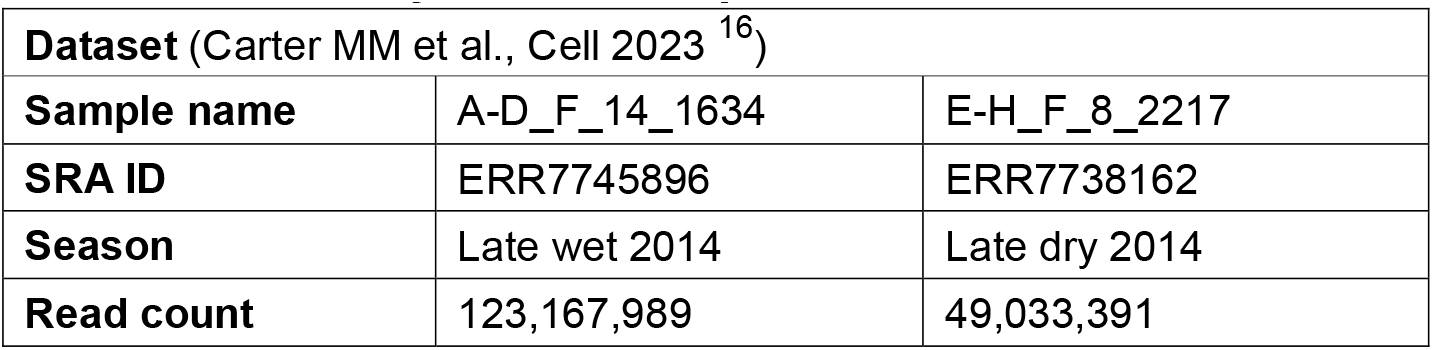
Summary of the example microbiome dataset.

| **Dataset** (Carter MM et al., Cell 2023 <sup>16</sup>) |  |  |
| --- | --- | --- |
| **Sample name** | A-D_F_14_1634 | E-H_F_8_2217 |
| **SRA ID** | ERR7745896 | ERR7738162 |
| **Season** | Late wet 2014 | Late dry 2014 |
| **Read count** | 123,167,989 | 49,033,391 |

Although the example microbiome dataset is from human gut, the protocol will also work for metagenomic and metatranscriptomic data in other ecological microbiomes (e.g., soil, sediment, marine, fresh water, wastewater, plant, forest, food). This protocol can also be easily modified for CAZyme annotation in microbiome data with more than two samples. We will perform the end-to-end analysis of the Carter2023 microbiome data, covering everything from sequencing read pre-processing to data visualization of the occurrence and abundance of CAZymes, CGCs, and substrates in the two stool microbiome samples (**Fig. 2**).

#### Read assembly strategies

For all the sequencing reads, quality trimming and cleaning are the first step to obtain high-quality clean reads. Kraken2^50^ is run to determine any possible sources of contamination reads, e.g., from human hosts or other eukaryotic sources. This step will help to decide what reference genomes or databases can be used to filter out the contamination reads. The final clean reads are then assembled into contigs using MEGAHIT^51^, a metagenome assembler. However, different strategies exist for multi-sample read assembly, a key step for the experimental design of CAZyme annotation that will directly impact the downstream data processing workflow (**Fig. 3**).

**Fig. 3:**
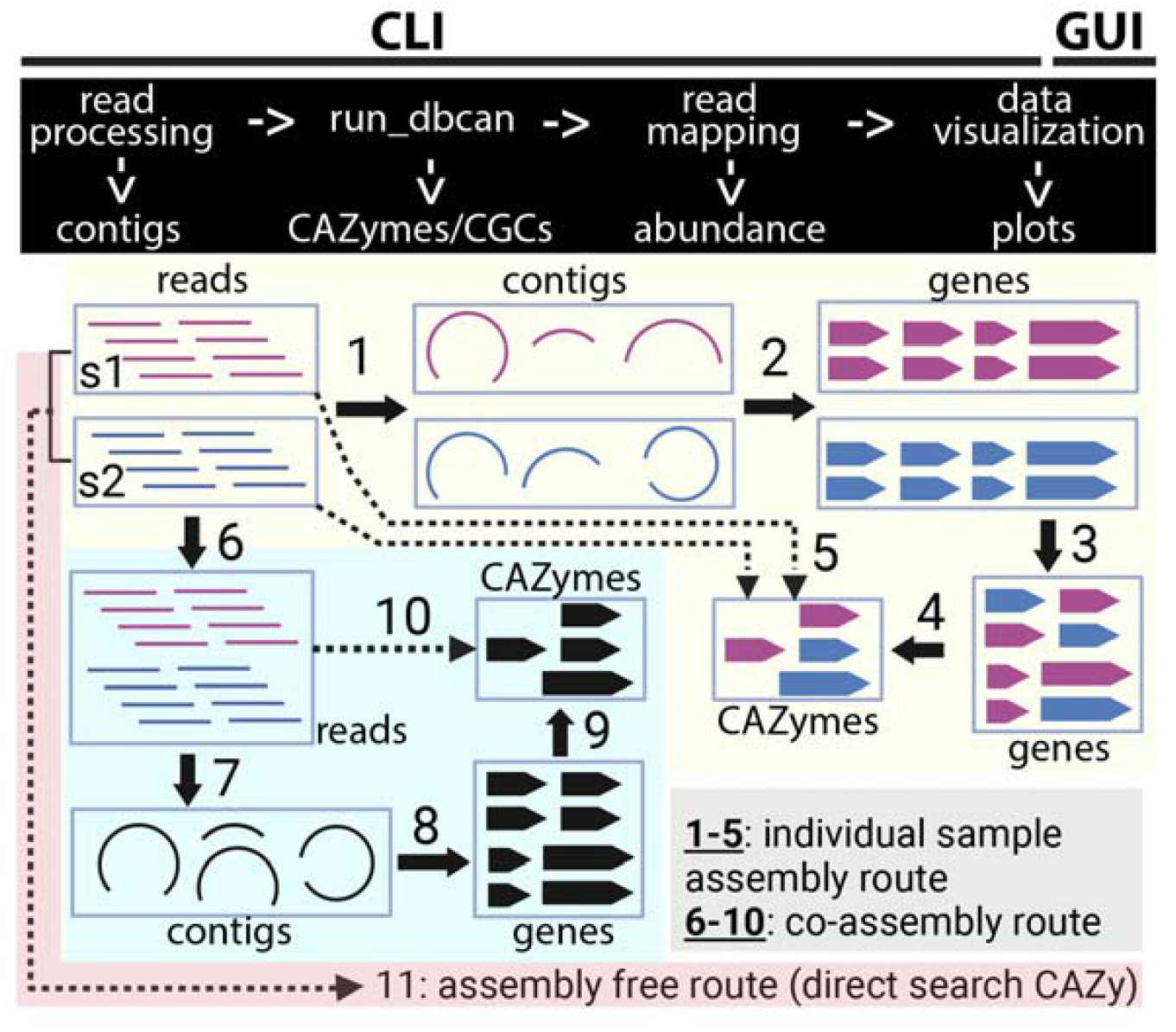
Experimental design of CAZyme annotation in microbiomes. The workflow consists of four modules shown on the top. All modules are installed and run in a command line interface (CLI) on a Linux computer. Once the data visualization module is finished, users should transfer the data plots (in PDF) to a computer with a graphical user interface (GUI) for visualization. Reads of two example microbiome samples can be processed with three routes for CAZyme annotation. One route involves read assembly of each individual sample, while the second requires pooling the reads first and co-assemble the pool reads. Numbers are the analysis steps. Once CAZymes and CGCs are identified by run_dbcan, reads are mapped back (dashed lines, 5 and 10) to calculate the abundance in different samples. The third route is assembly free, and does not need run_dbcan but directly search reads against the CAZy pre-annotated CAZyme protein sequences (or CAZyDB). The individual sample assembly route can be slightly modified by sub-sampling a subset of reads instead of using all reads for assembly, which could be a 4^th^ route.

The first strategy is an individual sample assembly route, where reads of each sample are individually assembled (step1 of **Fig. 3**). The resulting contigs are then subject to gene prediction (step2). The predicted proteins of all samples can be optionally combined to generate a non-redundant set of proteins (step3, e.g., by mmseqs2^54^ clustering with sequence identity > 0.95 and coverage > 0.95) as the total catalog of the encoded protein content in all the samples. The contigs and the non-redundant protein catalog are used as input to run_dbcan for the prediction of CAZymes, CGCs, and substrates (step4). The sequencing reads of each sample are mapped back to CAZymes and CGCs (step5) to calculate the abundances at the CAZyme family, subfamily, CGC, and substrate levels (**Fig. 4**).

**Fig. 4:**
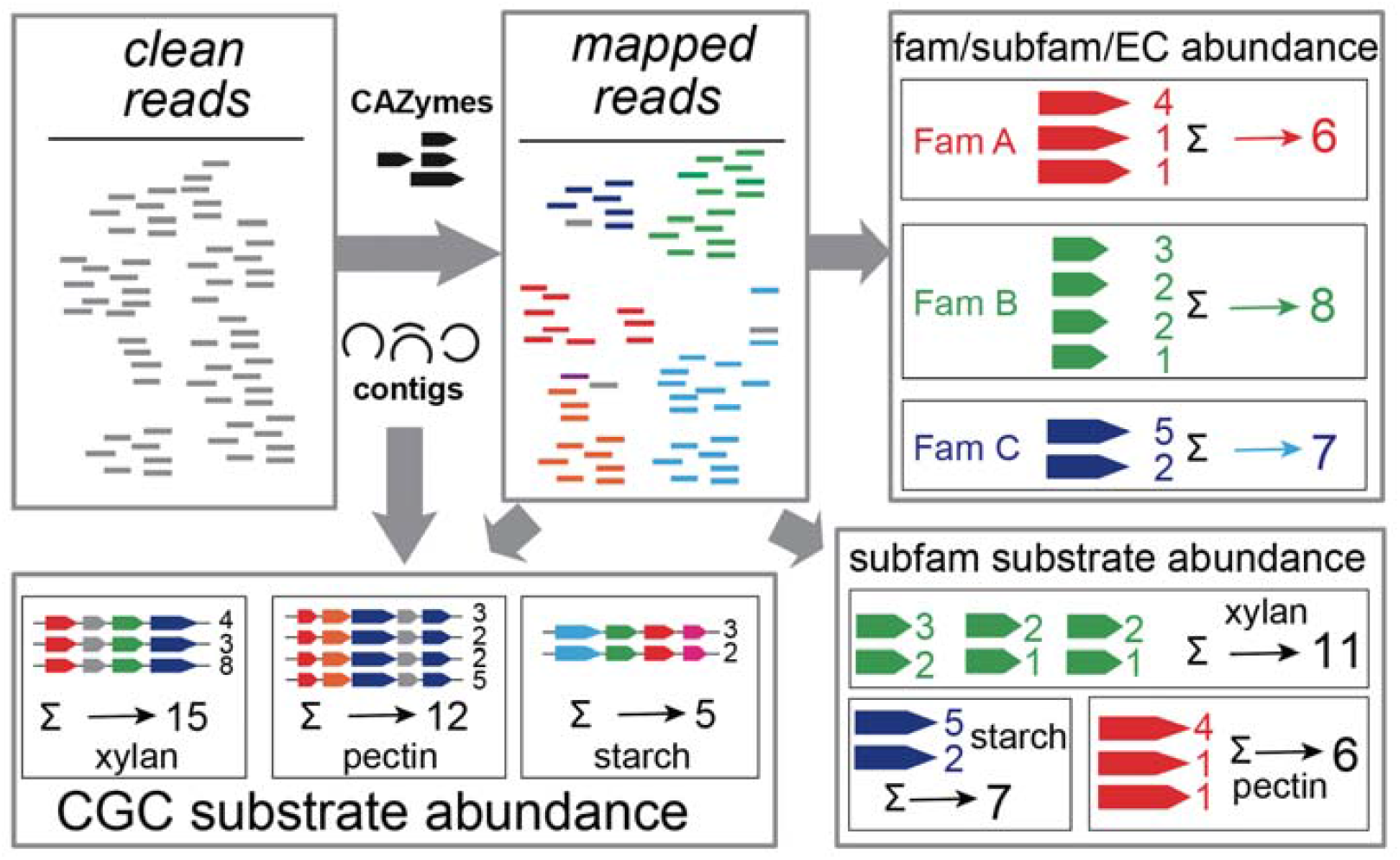
Abundance calculation for CAZyme families, subfamilies, and substrates. CAZyme coding sequences and contig nucleotide sequences are used as references for read mapping. CAZyme family and subfamily abundances are calculated as the sum of normalized read counts (TPM) of each CAZyme gene. CGC abundance is calculated as the mean of TPM of all genes in the gene cluster. Based on CAZyme subfamily abundance and CGC abundance, substrate abundances are calculated as the sum of TPM of all subfamilies or CGCs of the substrate group (predicted by run_dbcan). Numbers are the normalized read counts as examples.

The second strategy is a co-assembly route, meaning reads of the multiple samples are pooled together (step6) for assembly into contigs (step7). The gene prediction (step8), CAZyme annotation (step9), and read mapping back (step10) are conducted in the same way as the individual sample assembly route. The two strategies were recently compared, and evidence was shown that co-assembly resulted in a higher number of complete genes and a lower number of fragmented genes^55^. More importantly, co-assembly captured contigs and genes with low abundance in the samples. However, co-assembly requires substantially more computing resources and much longer running time, which is not realistic for pooling a larger number of samples (e.g., > 5). Also, different samples likely contain closely related but not identical strains. This strain heterogeneity may cause significant problems for the co-assembly process (e.g., confuse the de Bruijn graph) leading to more fragmented and shorter contigs. Therefore, the individual sample assembly is a much more practical and widely used strategy^56^.

In addition to assembly-based approaches, the assembly-free strategy is also used, e.g., in the popular general metagenomic functional annotation tool HUMAnN3^57^ and the inferred fiber degradation profile tool (IFDP)^43^. The assembly-free approach does not need run_dbcan but rather directly searches sequencing reads against the CAZy annotated protein sequence database (i.e., CAZyDB). A recent metatranscriptomic workflow showed that the assembly-free approach had much higher false positive rate and lower precision than the assembly-based approach^58^. However, as it skips the time-consuming read assembly step, the assembly-free route (step11, **Fig. 3**) is expected to have the advantage in running speed. Therefore, we also provide commands in this protocol paper to directly map sequencing reads to the CAZyDB for assembly-free CAZyme annotation. Note this approach can only perform the annotation at the CAZyme family level and is not able to predict CGCs, which requires genomic context information in the assembled contigs.

#### CAZyme annotation at family, subfamily, CGC, and substrate levels

Prokka is employed for gene finding and annotation^52^ (step2 and step8 in **Fig. 3**). Prokka calls Prodigal^59^ to predict protein sequences, also integrates other tools to predict RNA genes, and provides functional descriptions for predicted proteins based on homology searches. The essential output files comprise of gene structure file (gff) and protein sequence file (faa). For CAZyme annotation (step4 and step9), run_dbcan integrates results from three approaches to achieve more precise CAZyme annotation (**Table 1, Fig. 1**). It further identifies three additional classes of proteins (TCs, STPs, and TFs), and with the Prokka gff file predicts CGCs. It goes on to predict glycan substrates for CAZymes and CGCs using a manually curated CAZyme subfamily → EC → substrate mapping table, homology search against dbCAN-PUL^36^, and majority voting (**Table 1**, details in the dbCAN3 paper ^12^).

#### Reads mapping and abundance calculation

To estimate the abundance of CAZymes, CGCs, and substrates in microbiome samples, clean reads are mapped using BWA^60^ to the CAZyme coding sequences (CDS) identified in step4 and step9 (**Fig. 3**) and the entire contigs assembled in step1 and step7 (**Fig. 3**). Mapping to CAZyme CDS will be used to calculate the abundance of CAZymes and CGCs (**Fig. 4**), which are further used for calculating the abundance of families, subfamilies, ECs, and substrates.

The abundance is calculated as TPM (Transcripts Per Million), commonly used in RNA sequencing data analysis^61^. Other measures like RPM (Read Per Million) or RPKM/FPKM (Read/Fragment Per Kilobase per Million reads) are also calculated in this protocol. We first calculate three types of abundances (**Fig. 4**): (i) CAZyme family abundance, (ii) CAZyme subfamily abundance, (iii) CGC abundance. Family and subfamily abundances are calculated as the sum of the TPM of all CAZymes assigned to the respective family and subfamily. CGC abundance is calculated as the average TPM of all the genes in the CGC.

With the three abundances, we further calculate three abundances for glycan substrates (**Fig. 4**): (i) CAZyme subfamily-based glycan substrate abundance (sum of TPM of all CAZymes annotated as having the same substrate according to dbCAN-sub search); (ii) CGC substrate abundance based on homology search against dbCAN-PUL (sum of TPM of all CGCs annotated as having the same substrate according to dbCAN-PUL search); (iii) CGC substrate abundance based on majority voting (sum of TPM of all CGCs annotated as having the same substrate according to dbCAN-sub search, **Table 1**, details in the dbCAN3 paper^12^).

#### Data visualization

To facilitate visualization of the occurrence and abundance results, Python scripts are developed to generate bar plots, heatmap plots, and read mapping coverage plots.

## Materials

### Equipment

#### Operating system

All the modules of this protocol (**Fig. 3**) are designed to run on a command line (CLI) environment with a Linux OS (e.g., Ubuntu). We recommend users install these modules and execute all commands on a high-performance Linux cluster or workstation with >32 CPUs and 128GB RAM instead of a laptop, as the assembly of raw reads has a high demand of CPU and RAM.

Once users finish the data visualization module (**Fig. 3**), the resulting image files (PDF format) can be copied to a desktop or laptop with GUI for data visualization. In practice, users can choose not to use our read processing module and read mapping module. They may instead use their preferred tools for preparing input data for run_dbcan module and for calculating abundance for CAZymes and substrates. In that case, they can skip the installation of our read processing module and read mapping module in this protocol.

#### Data files

The example dataset (Carter2023) is described above and detailed in **Table 2**. The raw read data, intermediate data from each analysis step, and final result data and visualization files are organized in nested folders available on our website https://bcb.unl.edu/dbCAN_tutorial/dataset1-Carter2023/, **Fig. 5**) and https://dbcan.readthedocs.io. These websites also include data files and protocols for two additional example datasets (Wastyk2021^22^ and Priest2023^62^), which are available in **Supplementary Protocols**. We will use the independent sample assembly route for Carter2023 in the main text to demonstrate all the commands. Commands for the other routes are provided in **Supplementary Protocols**.

**Fig. 5:**
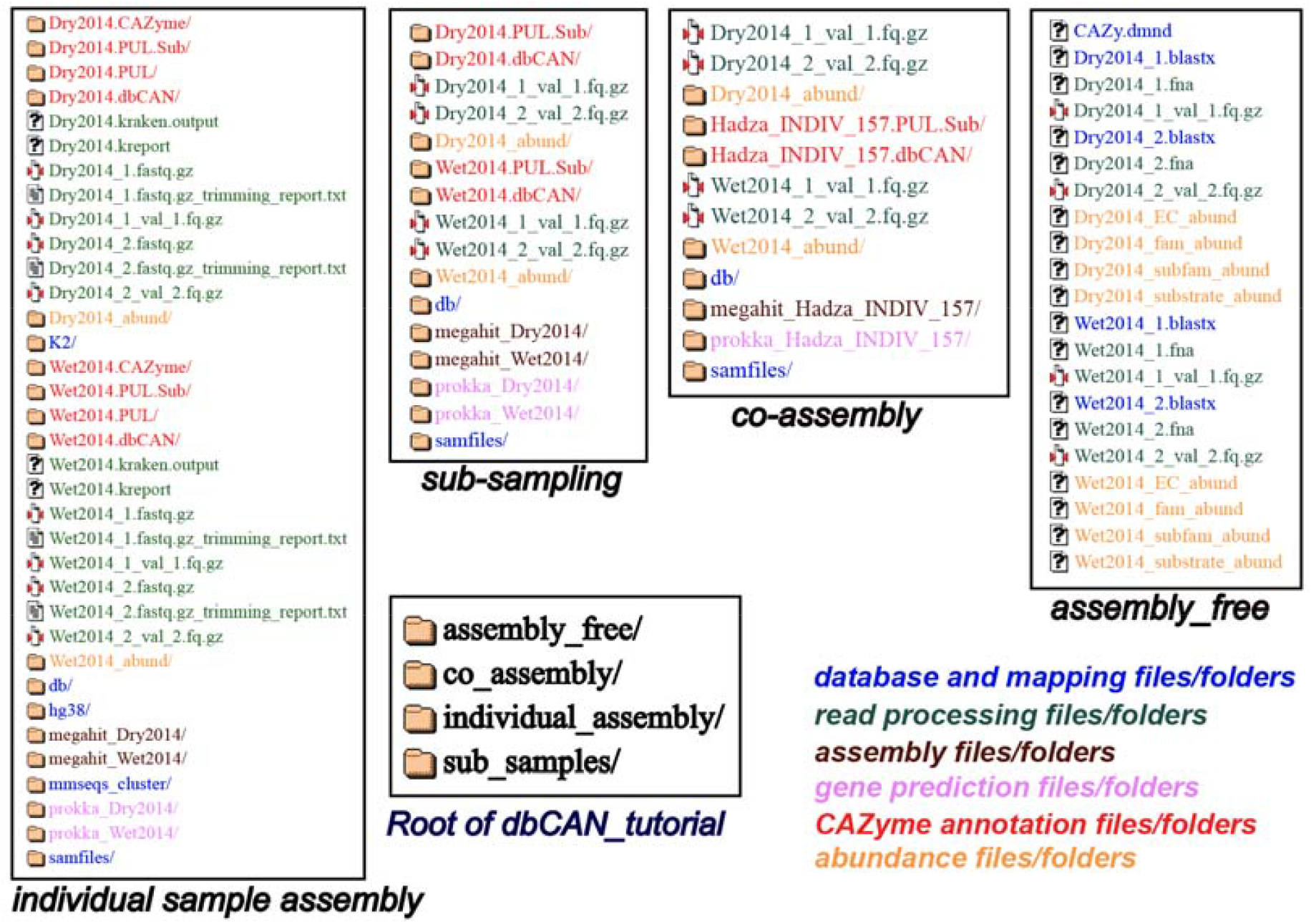
Screenshots of dbCAN tutorial website to show the file organization. The root folder (https://bcb.unl.edu/dbCAN_tutorial/dataset1-Carter2023/) of the website has four folders corresponding to four routes for CAZyme annotation of the Carter2023 dataset. The sub-sampling route is essentially the same as the individual sample assembly route except that only a subset of reads is used. The content of each folder is color coded according to the data analysis types. Commands for the individual sample assembly route are in the main text, while commands for other routes are available in the **Supplementary Protocols**.

#### Software and versions

- Anaconda (https://www.anaconda.com, version 23.7.3)
- MEGAHIT^51^ (https://github.com/voutcn/megahit, version 1.2.9)
- BWA^60^ (https://github.com/lh3/bwa, version 0.7.17-r1188)
- HMMER^63^ (http://hmmer.org/, version 3.3)
- DIAMOND^64^ (https://github.com/bbuchfink/diamond, version 2.1.8)
- BLAST^65^ (https://ftp.ncbi.nih.gov/blast/, version 2.14)
- TrimGalore^49^ (https://github.com/FelixKrueger/TrimGalore, version 0.6.0)
- Prokka^52^ (https://github.com/tseemann/prokka, version 1.4)
- Samtools^66^ (https://github.com/samtools/samtools, version 1.7)
- Seqkit^67^ (https://bioinf.shenwei.me/seqkit/, version 2.5.1)
- Bedtools^68^ (https://bedtools.readthedocs.io/en/latest/, version 2.27.1)
- Kraken2^50^ (https://ccb.jhu.edu/software/kraken2/, version 2.1.1)
- run_dbcan (https://github.com/linnabrown/run_dbcan, version 4.0.0)
- BBTools (https://jgi.doe.gov/data-and-tools/software-tools/bbtools/, version 37.62)
- Seqkt (https://github.com/lh3/seqtk, version 1.2-r94)
- Minimap2 (https://github.com/lh3/minimap2, version 2.26-r1175)
- Flye (https://github.com/fenderglass/Flye, version 2.9.3-b1797)
- Mmseqs2 (https://github.com/soedinglab/MMseqs2, release 15-6f452)

### Equipment setup

The setup has four steps (S1-S4).

#### CRITICAL STEP

Anaconda will be used as the software package management system for this protocol. Anaconda uses the conda command to create a virtual environment to facilitate the easy installation of software packages and running command line jobs. With the conda environment, users do not need worry about the potential issues of package dependencies and version conflicts. Like in all bioinformatics data analysis tasks, we recommend users organize their data files by creating a dedicated folder for each data analysis step. In this protocol paper, all computer code is written in `Courier New` font and starts with a ‘$’ sign as they are shown in a terminal console.

##### S1 Download Carter2023 (Table 2) raw reads (∼10min)

~~~
*$ wget
https://bcb.unl.edu/dbCAN_tutorial/dataset1-Carter2023/individual_assembly/Dry2014_1.fastq.gz
$ wget
https://bcb.unl.edu/dbCAN_tutorial/dataset1-Carter2023/individual_assembly/Dry2014_2.fastq.gz
$ wget
https://bcb.unl.edu/dbCAN_tutorial/dataset1-Carter2023/individual_assembly/Wet2014_1.fastq.gz
$ wget
https://bcb.unl.edu/dbCAN_tutorial/dataset1-Carter2023/individual_assembly/Wet2014_2.fastq.gz*
~~~

These raw data were originally downloaded from https://www.ncbi.nlm.nih.gov/sra/?term=ERR7745896 and https://www.ncbi.nlm.nih.gov/sra/?term=ERR7738162 and renamed to indicate their collected seasons (**Table 2**).

##### S2 Install Anaconda (∼3min)

Download and install the latest version of Anaconda for Linux from https://www.anaconda.com/download#downloads. Once Anaconda is successfully installed, proceed to create a dedicated conda environment named “CAZyme_annotation” and activate it. Subsequently, all the required tools can be seamlessly installed within this environment.

~~~
*$ conda create -n CAZyme_annotation python=3.9
$ conda activate CAZyme_annotation*
~~~

##### S3 Install all bioinformatics tools (∼10min)

~~~
*$ conda install -c conda-forge -c bioconda -c defaults prokka -y
$ conda install -c bioconda megahit trim-galore -y
$ conda install -c bioconda blast bwa diamond -y
$ conda install -c bioconda hmmer -y
$ conda install -c bioconda samtools bedtools seqkit -y
$ conda install -c bioconda kraken2 -y
$ conda install -c agbiome bbtools
$ conda install -c bioconda seqtk flye minimap2
$ conda install -c conda-forge -c bioconda mmseqs2
$ conda install dbcan -c conda-forge -c bioconda*
~~~

#### CRITICAL STEP

Alternatively, users can run a single configuration file dbcan.yml (replace S2 and S3) that streamlines the above configuration of all the essential software required for this protocol.

~~~
*$ git clone https://github.com/linnabrown/run_dbcan.git
$ cd run_dbcan
$ conda env create -f dbcan.yml
$ conda activate CAZyme_annotation*
~~~

##### S4 Configure databases required by run_dbcan (∼2h)

Various data files are needed for run_dbcan. Run the following commands to download and configure the required data files:

~~~
*$ mkdir db
$ cd db
$ wget http://bcb.unl.edu/dbCAN2/download/Databases/V12/CAZyDB.07262023.fa
$ mv CAZyDB.07262023.fa CAZyDB.fa && diamond makedb --in CAZyDB.fa -d CAZy
$ wget https://bcb.unl.edu/dbCAN2/download/Databases/V12/dbCAN-HMMdb-V12.txt
$ mv dbCAN-HMMdb-V12.txt dbCAN.txt && hmmpress dbCAN.txt
$ wget http://bcb.unl.edu/dbCAN2/download/Databases/dbCAN_sub.hmm
$ hmmpress dbCAN_sub.hmm
$ wget https://bcb.unl.edu/dbCAN2/download/Databases/V12/tcdb.fa
$ diamond makedb --in tcdb.fa -d tcdb
$ wget http://bcb.unl.edu/dbCAN2/download/Databases/V12/tf-1.hmm && hmmpress tf-1.hmm
$ wget http://bcb.unl.edu/dbCAN2/download/Databases/V12/tf-2.hmm && hmmpress tf-2.hmm
$ wget https://bcb.unl.edu/dbCAN2/download/Databases/V12/stp.hmm && hmmpress stp.hmm
$ wget http://bcb.unl.edu/dbCAN2/download/Databases/fam-substrate-mapping-08012023.tsv
$ mv fam-substrate-mapping-08012023.tsv fam-substrate-mapping.tsv
$ wget http://bcb.unl.edu/dbCAN2/download/Databases/CAZyID_subfam_mapping.tsv
$ wget http://bcb.unl.edu/dbCAN2/download/Databases/subfam_EC_mapping.tsv
$ wget http://bcb.unl.edu/dbCAN2/download/Databases/PUL.faa
$ makeblastdb -in PUL.faa -dbtype prot
$ wget http://bcb.unl.edu/dbCAN2/download/Databases/dbCAN-PUL_12-12-2023.xlsx
$ mv dbCAN-PUL_12-12-2023.xlsx dbCAN-PUL.xlsx
$ wget http://bcb.unl.edu/dbCAN2/download/Databases/dbCAN-PUL.tar.gz
$ tar xvf dbCAN-PUL.tar.gz && rm dbCAN-PUL.tar.gz
$ cd* ..
~~~

Download database required by Kraken2 (very slow; can be skipped if users do not intend to run Kraken2):

~~~
*$ kraken2-build --standard --db K2*
~~~

#### CRITICAL STEP

The downloaded files must be all in the right location (the db folder). The `CAZyDB.07262023.fa` file is needed for DIAMOND search (**Table 1**). The `dbCAN-HMMdb-V12.txt` and `dbCAN_sub.hmm` files are for HMMER search. The `tcdb.fa, tf-1.hmm, tf-2.hmm`, and `stp.hmm` files are for CGC prediction. The `PUL.faa` file consists of protein sequences from experimentally validated PULs for BLAST search to predict substrates for CGCs. The `dbCAN-PUL_12-12-2023.txt` and `dbCAN-PUL_12-12-2023.xlsx` files contain PUL-substrate mapping curated from literature. Lastly, the `fam-substrate-mapping-08012023.tsv` file is the family-EC-substrate mapping table for the prediction of CAZyme substrates.

**! CAUTION**

Users should use a clean version of Anaconda. If the above steps failed, we suggest users reinstall their Anaconda. The Anaconda installation and configuration step may experience prolonged time while resolving environment dependencies. Users should be patient during this process. Alternatively, users may consider “mamba”, another Python package manager that offers similar functionality to Anaconda. Information and access to mamba software can be found at https://github.com/mamba-org/mamba.

If all these failed, users may use our docker image (https://dbcan.readthedocs.io/en/latest/user_guide/run_from_raw_reads.html) with the following steps:

1. Install Docker on your system (e.g., Linux, MacOS);
2. Pull the image haidyi/cazyme_annotation from Docker Hub;
3. Run the cazyme_annotation tool via Docker (all above tools have been built into the Docker image).

### Procedure

We use the Carter2023 dataset (**Table 2**) and the individual sample assembly route of **Fig. 3** for this procedure. **Supplementary Protocols** contains the procedures for the co-assembly route and assembly-free route for this dataset.

The procedure has 4 modules (**Fig. 3**) and 16 steps (P1-P16).

All the data including the pre-computed intermediate files and final files can be found at https://bcb.unl.edu/dbCAN_tutorial/dataset1-Carter2023/individual_assembly/ (**Fig. 5**).

#### Module 1: Reads processing (Fig. 3) to obtain contigs

##### P1 Check contamination (TIMING ∼10min)

~~~
*$ kraken2 --threads 32 --quick --paired --db K2 --report Wet2014.kreport --output Wet2014*.
*kraken.output Wet2014_1.fastq.gz Wet2014_2.fastq.gz
$ kraken2 --threads 32 --quick --paired --db K2 --report Dry2014.kreport --output Dry2014*.
*kraken.output Dry2014_1.fastq.gz Dry2014_2.fastq.gz*
~~~

Kraken2 found very little contamination in the Carter2023 data. Consequently, there was no need for the contamination removal step.

If contamination is identified, users can align the reads to the reference genomes of potential contamination source organisms to remove the aligned reads (**Box 1**). The most common source in human microbiome studies is from human hosts.

###### Box 1

**Example to remove contamination reads from human**

Kraken2 will produce the following output files.

~~~
*-rw-rw-r-- 1 jinfang jinfang 2.0G Dec 12 10:24 Dry2014.kraken.output
-rw-rw-r-- 1 jinfang jinfang 1.2M Dec 12 10:25 Dry2014.kreport
-rw-rw-r-- 1 jinfang jinfang 5.1G Dec 12 09:47 Wet2014.kraken.output
-rw-rw-r-- 1 jinfang jinfang 1.1M Dec 12 09:48 Wet2014.kreport*
~~~

Suppose from these files, we have identified humans as the contamination source, we can use the following commands to remove the contamination reads by aligning reads to the human reference genome.

~~~
*$ wget
https://ftp.ensembl.org/pub/release-110/fasta/homo_sapiens/dna/Homo_sapiens.GRCh38.dna.primary_assembly.fa.gz
$ bwa index -p hg38 Homo_sapiens.GRCh38.dna.primary_assembly.fa.gz
$ bwa mem hg38 Wet2014_1.fastq.gz Wet2014_2.fastq.gz -t 32 -o Wet2014.hg38.sam
$ bwa mem hg38 Dry2014_1.fastq.gz Dry2014_2.fastq.gz -t 32 -o Dry2014.hg38.sam
$ samtools view -f 12 Wet2014.hg38.sam > Wet2014.hg38.unmap.bam
$ samtools view -f 12 Dry2014.hg38.sam > Dry2014.hg38.unmap.bam
$ samtools fastq -1 Wet2014_1.clean.fq.gz -2 Wet2014_2.clean.fq.gz Wet2014.hg38.unmap.bam
$ samtools fastq -1 Dry2014_1.clean.fq.gz -2 Dry2014_2.clean.fq.gz Dry2014.hg38.unmap.bam*
~~~

##### P2 Trim adapter and low-quality reads (TIMING ∼20min)

~~~
*$ trim_galore --paired Wet2014_1.fastq.gz Wet2014_2.fastq.gz --illumina -j 36
$ trim_galore --paired Dry2014_1.fastq.gz Dry2014_2.fastq.gz --illumina -j 36*
~~~

We specified `--illumina` to indicate that the reads were generated using the Illumina sequencing platform. Nonetheless, trim_galore can automatically detect adapters, providing flexibility for users who may know the specific sequencing platform. Details of trimming are available in the trimming report file (**Box 2**).

###### Box 2

**Example output of trim_galore**

In addition to the trimmed read files, Trim_galore also generates a trimming report file. The trimming report contains details on read trimming, such as the number of trimmed reads.

~~~
*-rw-rw-r-- 1 jinfang jinfang 4.2K Dec 13 01:48 Dry2014_1.fastq.gz_trimming_report.txt
-rw-rw-r-- 1 jinfang jinfang 2.0G Dec 13 01:55 Dry2014_1_val_1.fq.gz
-rw-rw-r-- 1 jinfang jinfang 4.4K Dec 13 01:55 Dry2014_2.fastq.gz_trimming_report.txt
-rw-rw-r-- 1 jinfang jinfang 2.4G Dec 13 01:55 Dry2014_2_val_2.fq.gz
-rw-rw-r-- 1 jinfang jinfang 4.4K Dec 13 01:30 Wet2014_1.fastq.gz_trimming_report.txt
-rw-rw-r-- 1 jinfang jinfang 3.4G Dec 13 01:46 Wet2014_1_val_1.fq.gz
-rw-rw-r-- 1 jinfang jinfang 4.6K Dec 13 01:46 Wet2014_2.fastq.gz_trimming_report.txt
-rw-rw-r-- 1 jinfang jinfang 3.7G Dec 13 01:46 Wet2014_2_val_2.fq.gz*
~~~

**! CAUTION**

During the trimming process, certain reads may be entirely removed due to low quality in its entirety. Using the `--retain_unpaired` parameter in trim_galore allows for the preservation of single-end reads. In this protocol, this option was not selected, so that both reads of a forward-revise pair were removed.

##### P3 Assemble reads into contigs (TIMING ∼4h20min)

~~~
*$ megahit -m 0.5 -t 32 -o megahit_ Wet2014 -1 Wet2014_1_val_1.fq.gz -2 Wet2014_2_val_2.fq.gz
--out-prefix Wet2014 --min-contig-len 1000
$ megahit -m 0.5 -t 32 -o megahit_ Dry2014 -1 Dry2014_1_val_1.fq.gz -2 Dry2014_2_val_2.fq.gz
--out-prefix Dry2014 --min-contig-len 1000*
~~~

MEGAHIT generates two output folders: `megahit_Wet2014` and `megahit_Dry2014`. Each contains five files and one sub-folder (**Box 3**). `Wet2014.contigs.fa` is the final contig sequence file. We set `--min-contig-len 1000`, a common practice to retain all contigs longer than 1,000 base pairs.

###### Box 3

**Example output of MEGAHIT**

~~~
*-rw-rw-r-- 1 jinfang jinfang 262 Dec 13 04:19 checkpoints.txt
-rw-rw-r-- 1 jinfang jinfang 0 Dec 13 04:19 done
drwxrwxr-x 2 jinfang jinfang 4.0K Dec 13 04:19 intermediate_contigs
-rw-rw-r-- 1 jinfang jinfang 1.1K Dec 13 02:22 options.json
-rw-rw-r-- 1 jinfang jinfang 258M Dec 13 04:19 Wet2014.contigs.fa
-rw-rw-r-- 1 jinfang jinfang 208K Dec 13 04:19 Wet2014.log*
~~~

**! CAUTION**

A common practice in metagenomics after assembly is to further bin contigs into metagenome-assembled genomes (MAGs)^24,69^. However, in this protocol, we chose not to generate MAGs because not all contigs can be binned into MAGs, and those un-binned contigs can also encode CAZymes. Users may refer to a recent paper^24^ for the protocol to generate MAGs. All the remaining steps will directly work for MAGs as well.

##### P4 Predict genes by Prokka (TIMING ∼21h)

~~~
*$ prokka --kingdom Bacteria --cpus 32 --outdir prokka_ Wet2014 --prefix Wet2014 --addgenes --addmrna
--locustag Wet2014 megahit_ Wet2014/Wet2014.contigs.fa
$ prokka --kingdom Bacteria --cpus 32 --outdir prokka_ Dry2014 --prefix Dry2014 --addgenes --addmrna
--locustag Dry2014 megahit_ Dry2014/Dry2014.contigs.fa*
~~~

The parameter `--kingdom Bacteria` is required for bacterial gene prediction. To optimize performance, `--CPU 32` instructs the utilization of 32 CPUs. Reduce this number if you do not have this many CPUs on your computer. The output files comprise of both protein and CDS sequences in Fasta format (e.g., `Wet2014.faa` and `Wet2014.ffn` in **Box 4**).

###### Box 4

**Example output of Prokka**

~~~
*-rw-rw-r-- 1 jinfang jinfang 8.4M Dec 14 00:51 Wet2014.err
-rw-rw-r-- 1 jinfang jinfang 75M Dec 13 21:38 Wet2014.faa
-rw-rw-r-- 1 jinfang jinfang 204M Dec 13 21:38 Wet2014.ffn
-rw-rw-r-- 1 jinfang jinfang 259M Dec 13 20:47 Wet2014.fna
-rw-rw-r-- 1 jinfang jinfang 264M Dec 13 21:38 Wet2014.fsa
-rw-rw-r-- 1 jinfang jinfang 599M Dec 14 00:52 Wet2014.gbk
-rw-rw-r-- 1 jinfang jinfang 372M Dec 13 21:38 Wet2014.gff
-rw-rw-r-- 1 jinfang jinfang 2.2M Dec 14 00:52 Wet2014.log
-rw-rw-r-- 1 jinfang jinfang 1.2G Dec 14 00:52 Wet2014.sqn
-rw-rw-r-- 1 jinfang jinfang 68M Dec 13 21:38 Wet2014.tbl
-rw-rw-r-- 1 jinfang jinfang 30M Dec 13 21:38 Wet2014.tsv
-rw-rw-r-- 1 jinfang jinfang 152 Dec 13 21:38 Wet2014.txt*
~~~

#### Module 2: run_dbcan annotation (Fig. 3) to obtain CAZymes, CGCs, and substrates

##### CRITICAL STEP

Users can skip P5 and P6, and directly run P7 (much slower though), if they want to predict not only CAZymes and CGCs, but also substrates.

###### P5 CAZyme annotation at the CAZyme family level (TIMING ∼10min)

~~~
*$ run_dbcan prokka_Wet2014/Wet2014.faa protein --hmm_cpu 32 --out_dir Wet2014.CAZyme --tools hmmer
--db_dir db
$ run_dbcan prokka_Dry2014/Dry2014.faa protein --hmm_cpu 32 --out_dir Dry2014.CAZyme --tools hmmer
--db_dir db*
~~~

Two arguments are required for run_dbcan: the input sequence file (`faa` files) and the sequence type (`protein`). By default, run_dbcan will use three methods (HMMER vs dbCAN HMMdb, DIAMOND vs CAZy, HMMER vs dbCAN-sub HMMdb) for CAZyme annotation (**Table 1, Fig. 2**). This default setting is equivalent to the use `--tools all` parameter (**Box 5**). Here we only invoke the HMMER vs dbCAN HMMdb for CAZyme annotation at the family level.

###### Box 5

**CAZyme annotation with default setting**

If the `--tools` parameter is not set, it is the default setting, which is the same as *--tools all*. This will take much longer time to finish (∼5h) due to the large size of dbCAN-sub HMMdb (used for substrate prediction for CAZymes, see **Table 1**).

~~~
*$ run_dbcan prokka_Wet2014/Wet2014.faa protein --out_dir Wet2014.CAZyme --dia_cpu 32 --hmm_cpu 32
--dbcan_thread 32 --tools all
$ run_dbcan prokka_Dry2014/Dry2014.faa protein --out_dir Dry2014.CAZyme --dia_cpu 32 --hmm_cpu 32
--dbcan_thread 32 --tools all*
~~~

The sequence type can be `protein, prok, meta.` If the input sequence file contains metagenomic contig sequences (`fna` file), the sequence type has to be `meta`, and Prodigal will be called to predict genes.

~~~
*$ run_dbcan prokka_Wet2014/Wet2014.fna meta --out_dir Wet2014.CAZyme --dia_cpu 32 --hmm_cpu 32
--dbcan_thread 32
$ run_dbcan prokka_Dry2014/Dry2014.fna meta --out_dir Dry2014.CAZyme --dia_cpu 32 --hmm_cpu 32
--dbcan_thread 32*
~~~

###### P6 CGC prediction (TIMING ∼15 min)

The following commands will re-run run_dbcan to not only predict CAZymes but also CGCs with protein `faa` and gene location `gff` files.

~~~
*$ run_dbcan prokka_Wet2014/Wet2014.faa protein --tools hmmer --tf_cpu 32 --stp_cpu 32 -c
prokka_Wet2014/Wet2014.gff --out_dir Wet2014.PUL --dia_cpu 32 --hmm_cpu 32
$ run_dbcan prokka_Dry2014/Dry2014.faa protein --tools hmmer --tf_cpu 32 --stp_cpu 32 -c
prokka_ Dry2014/Dry2014.gff --out_dir Dry2014.PUL --dia_cpu 32 --hmm_cpu 32*
~~~

As mentioned above (**Table 1, Fig. 2**), CGC prediction is a featured function added into dbCAN2 in 2018. To identify CGCs with the `protein` sequence type, a gene location file (`gff`) must be provided together. If the input sequence type is `prok` or `meta`, meaning users only have contig `fna` files, the CGC prediction can be activated by setting `-c cluster`.

**! CAUTION**

If the users would like to create their own `gff` file (instead of using Prokka or Prodigal), it is important to make sure the value of ID attribute in the `gff` file matches the protein ID in the protein `faa` file.

#### ?Troubleshooting

If no result is found in CGC output file, it is most likely because the sequence IDs in `gff` file and `faa` file do not match. Another less likely reason is that the contigs are too short and fragmented and not suitable for CGC prediction.

##### P7 Substrate prediction for CAZymes and CGCs (TIMING ∼5h)

The following commands will re-run run_dbcan to predict CAZymes, CGCs, and their substrates with the `--cgc_substrate` parameter.

~~~
*$ run_dbcan prokka_Wet2014/Wet2014.faa protein --dbcan_thread 32 --tf_cpu 32 --stp_cpu 32 -c
prokka_Wet2014/Wet2014.gff --cgc_substrate --hmm_cpu 32 --out_dir Wet2014.dbCAN --dia_cpu 32
$ run_dbcan prokka_Dry2014/Dry2014.faa protein --dbcan_thread 32 --tf_cpu 32 --stp_cpu 32 -c
prokka_Dry2014/Dry2014.gff --cgc_substrate --hmm_cpu 32 --out_dir Dry2014.dbCAN --dia_cpu 32*
~~~

Files in the output folder is explained in **Box 6**.

**! CAUTION**

The above commands do not set the `--tools` parameter, which means all three methods for CAZyme annotation will be activated (**Box 5**). Because dbCAN-sub HMMdb (for CAZyme substrate prediction) is 200 times larger than dbCAN HMMdb, the runtime will be much longer. Users can specify `--tools hmmer`, so that the HMMER search against dbCAN-sub will be disabled. However, this will turn off the substrate prediction for CAZymes and CGCs based on CAZyme substrate majority voting. Consequently, the substrate prediction will be solely based on homology search against PULs in dbCAN-PUL (**Fig. 1, Table 1**).

~~~
*$ run_dbcan prokka_Wet2014/Wet2014.faa protein --tools hmmer --stp_cpu 32 -c
prokka_Wet2014/Wet2014.gff --cgc_substrate --out_dir Wet2014.PUL.Sub --dia_cpu 32 --hmm_cpu 32
--tf_cpu 32
$ run_dbcan prokka_Dry2014/Dry2014.faa protein --tools hmmer --stp_cpu 32 -c
prokka_Dry2014/Dry2014.gff --cgc_substrate --out_dir Dry2014.PUL.Sub --dia_cpu 32 --hmm_cpu 32
--tf_cpu 32*
~~~

###### Box 6

**Example output folder content of run_dbcan substrate prediction**

In the output directory (https://bcb.unl.edu/dbCAN_tutorial/dataset1-Carter2023/individual_assembly/Wet2014.dbCAN/), a total of 17 files and 1 folder are generated:

~~~
*-rw-rw-r-- 1 jinfang jinfang 33M Dec 17 09:36 PUL_blast.out
-rw-rw-r-- 1 jinfang jinfang 3.3M Dec 17 09:35 CGC.faa
-rw-rw-r-- 1 jinfang jinfang 18M Dec 17 09:35 cgc.gff
-rw-rw-r-- 1 jinfang jinfang 836K Dec 17 09:35 cgc.out
-rw-rw-r-- 1 jinfang jinfang 374K Dec 17 09:35 cgc_standard.out
-rw-rw-r-- 1 jinfang jinfang 1.8M Dec 17 09:35 cgc_standard.out.json
-rw-rw-r-- 1 jinfang jinfang 785K Dec 17 09:31 dbcan-sub.hmm.out
-rw-rw-r-- 1 jinfang jinfang 511K Dec 17 09:31 diamond.out
-rw-rw-r-- 1 jinfang jinfang 638K Dec 17 09:31 dtemp.out
-rw-rw-r-- 1 jinfang jinfang 414K Dec 17 09:31 hmmer.out
-rw-rw-r-- 1 jinfang jinfang 386K Dec 17 09:35 overview.txt
-rw-rw-r-- 1 jinfang jinfang 2.8M Dec 17 09:35 stp.out
-rw-rw-r-- 1 jinfang jinfang 63K Dec 17 09:36 substrate.out
drwxrwxr-x 2 jinfang jinfang 36K Dec 17 09:39 synteny.pdf
-rw-rw-r-- 1 jinfang jinfang 799K Dec 17 09:32 tf-1.out
-rw-rw-r-- 1 jinfang jinfang 645K Dec 17 09:34 tf-2.out
-rw-rw-r-- 1 jinfang jinfang 2.3M Dec 17 09:35 tp.out
-rw-rw-r-- 1 jinfang jinfang 75M Dec 17 02:07 uniInput*
~~~

**PUL_blast**.**out**: BLAST results between CGCs and PULs.

**CGC**.**faa**: Protein Fasta sequences encoded in all CGCs.

**cgc**.**gff**: reformatted from the user input gff file by marking CAZymes, TFs, TCs, and STPs.

**cgc**.**out**: raw output of CGC predictions.

**cgc_standard**.**out**: simplified version of cgc.out for easy parsing in TSV format. An example row has the following columns:

1. CGC_id: *CGC1*
2. type: *CAZyme*
3. contig_id: *k141_272079*
4. gene_id: *Wet2014_00308*
5. start: *5827 6*.
6. end: *7257*
7. strand: *-*
8. annotation: *GH1*

*Explanation: the gene Wet2014_00308 encodes a GH1 CAZyme in the CGC1 of the contig k141_272079. CGC1 also has other genes, which are provided in other rows. Wet2014_00308 is on the negative strand of k141_272079 from 5827 to 7257. The type can be one of the four signature gene types (CAZymes, TCs, TFs, STPs) or the null type (not annotated as one of the four signature genes)*.

**cgc_standard**.**out**.**json**: JSON format of cgc_standard.out.

**dbcan-sub**.**hmm**.**out**: HMMER search result against dbCAN-sub HMMdb, including a column with CAZyme substrates extracted from `fam-substrate-mapping-08012023.tsv`.

**diamond**.**out**: DIAMOND search result against the CAZy annotated protein sequences (`CAZyDB.07262023.fa`).

**dtemp**.**out**: temporary file.

**hmmer**.**out**: HMMER search result against dbCAN HMMdb.

**overview**.**txt**: summary of CAZyme annotation from three methods in TSV format. An example row has the following columns:

1. Gene_ID: *Wet2014_00076 2*.
2. EC#: *3.2.1.99:3*
3. dbCAN: *GH43_4(42-453)*
4. dbCAN_sub: *GH43_e149*
5. DIAMOND: *GH43_4*
6. #ofTools: *3*

*Explanation: the protein Wet2014_000076 is annotated by 3 tools to be a CAZyme: (1) GH43_4 (CAZy defined subfamily 4 of GH43) by HMMER vs dbCAN HMMdb with a domain range from aa position 42 to 453, (2) GH43_e149 (eCAMI defined subfamily e149; e indicates it is from eCAMI not CAZy) by HMMER vs dbCAN-sub HMMdb (derived from eCAMI subfamilies), and (3) GH43_4 by DIAMOND vs CAZy annotated protein sequences. The second column 3.2.1.99:3 is extracted from eCAMI, meaning that the eCAMI subfamily GH43_e149 contains 3 member proteins which have an EC 3.2.1.99 according to CAZy. In most cases, the 3 tools will have the same CAZyme family assignment. When they give different assignment. We recommend a preference order: dbCAN > eCAMI/dbCAN-sub > DIAMOND. See our dbCAN2 paper*^*11*^, *dbCAN3 paper*^*12*^, *and eCAMI*^*41*^ *for more details*.

*Note: If users invoked the --use_signalP parameter when running run_dbcan, there will be an additional column called signalP in the overview.txt*.

**stp**.**out**: HMMER search result against the MiST^70^ compiled signal transduction protein HMMs from Pfam.

**tf-1**.**out**: HMMER search result against the DBD^71^ compiled transcription factor HMMs from Pfam^72^.

**tf-2**.**out**: HMMER search result against the DBD compiled transcription factor HMMs from Superfamily^73^.

**tp**.**out**: DIAMOND search result against the TCDB ^74^ annotated protein sequences.

**substrate**.**out**: summary of substrate prediction results for CGCs in TSV format from two approaches^12^ (dbCAN-PUL blast search and dbCAN-sub majority voting). An example row has the following columns:

1. CGC_ID: *k141_227425*|*CGC1*
2. Best hit PUL_ID in dbCAN-PUL: *PUL0402*
3. Substrate of the hit PUL: *xylan*
4. Sum of bitscores for homologous gene pairs between CGC and PUL: *2134.0*
5. Types of homologous gene pairs: *TC-TC;CAZyme-CAZyme*
6. Substrate predicted by majority voting of CAZymes in CGC: *xylan*
7. Voting score: *2.0*

*Explanation: The CGC1 of contig k141_227425 has its best hit PUL0402 (from* `PUL_blast.out`*) with xylan as substrate (from* `dbCAN-PUL_12-12-2023.xlsx`*). Two signature genes are matched between k141_227425*|*CGC1 and PUL0402 (from* `PUL_blast.out`*): one is a CAZyme and the other is a TC. The sum of blast bit scores of the two homologous pairs (TC-TC and CAZyme-CAZyme) is 2134.0. Hence, the substrate of k141_227425*|*CGC1 is predicted to be xylan according to dbCAN-PUL blast search. The last two columns are based on the dbCAN-sub result (*`dbcan-sub.hmm.out`*), as the file tells that two CAZymes in k141_227425*|*CGC1 are predicted to have xylan substrate. The voting score is thus 2.0, so that according to the majority voting rule, k141_227425*|*CGC1 is predicted to have a xylan substrate*.

*Note: for many CGCs, only one of the two approaches produces substrate prediction. In some cases, the two approaches produce different substrate assignments. We recommend a preference order: dbCAN-PUL blast search > dbCAN-sub majority voting. See our dbCAN3 paper*^12^ *for more details*.

**synteny**.**pdf**: a folder with syntenic block alignment plots between all CGCs and PULs.

**uniInput**: renamed Fasta file from input protein sequence file.

#### Module 3: Read mapping (Fig. 3) to calculate abundance for CAZyme families, subfamilies, CGCs, and substrates

##### P8 Read mapping to all CDS of each sample (TIMING ∼20 min)

~~~
*$ bwa index prokka_Wet2014/Wet2014.ffn
$ bwa index prokka_Dry2014/Dry2014.ffn
$ mkdir samfiles
$ bwa mem -t 32 -o samfiles/Wet2014.CDS.sam prokka_Wet2014/Wet2014.ffn Wet2014_1_val_1.fq.gz
Wet2014 _2_val_2.fq.gz
$ bwa mem -t 32 -o samfiles/Dry2014.CDS.sam prokka_Dry2014/Dry2014.ffn Dry2014_1_val_1.fq.gz
Dry2014_2_val_2.fq.gz*
~~~

Reads are mapped to the `ffn` files from Prokka.

##### P9 Read mapping to all contigs of each sample (TIMING ∼20min)

~~~
*$ bwa index megahit_Wet2014/Wet2014.contigs.fa
$ bwa index megahit_Dry2014/Dry2014.contigs.fa
$ bwa mem -t 32 -o samfiles/Wet2014.sam megahit_Wet2014/Wet2014.contigs.fa Wet2014_1_val_1.fq.gz
Wet2014_2_val_2.fq.gz
$ bwa mem -t 32 -o samfiles/Dry2014.sam megahit_Dry2014/Dry2014.contigs.fa Dry2014_1_val_1.fq.gz
Dry2014_2_val_2.fq.gz*
~~~

Reads are mapped to the `contig` files from MEGAHIT.

##### P10 Sort SAM files by coordinates (TIMING ∼8min)

~~~
*$ cd samfiles
$ samtools sort -@ 32 -o Wet2014.CDS.bam Wet2014.CDS.sam
$ samtools sort -@ 32 -o Dry2014.CDS.bam Dry2014.CDS.sam
$ samtools sort -@ 32 -o Wet2014.bam Wet2014.sam
$ samtools sort -@ 32 -o Dry2014.bam Dry2014.sam
$ rm -rf *sam
$ cd* ..
~~~

##### P11 Read count calculation for all proteins of each sample using Bedtools (TIMING ∼2min)

~~~
*$ mkdir Wet2014_abund && cd Wet2014_abund
$ seqkit fx2tab -l -n -i* ..*/prokka_Wet2014/Wet2014.ffn* | *awk ‘{print $1”\t”$2}’ > Wet2014.length
$ seqkit fx2tab -l -n -i* ..*/prokka_Wet2014/Wet2014.ffn* | *awk ‘{print $1”\t”0”\t”$2}’ > Wet2014.bed
$ bedtools coverage -g Wet2014.length -sorted -a Wet2014.bed -counts
-b* ..*/samfiles/Wet2014.CDS.bam > Wet2014.depth.txt
$ cd* .. *&& mkdir Dry2014_abund && cd Dry2014_abund
$ seqkit fx2tab -l -n -i* ..*/prokka_Dry2014/Dry2014.ffn* | *awk ‘{print $1”\t”$2}’ > Dry2014.length
$ seqkit fx2tab -l -n -i* ..*/prokka_Dry2014/Dry2014.ffn* | *awk ‘{print $1”\t”0”\t”$2}’ > Dry2014.bed
$ bedtools coverage -g Dry2014.length -sorted -a Dry2014.bed -counts
-b* ..*/samfiles/Dry2014.CDS.bam > Dry2014.depth.txt
$ cd* ..
~~~

Read counts are saved in `depth.txt` files of each sample.

##### P12 Read count calculation for a given region of contigs using Samtools (TIMING ∼2min)

~~~
*$ cd Wet2014_abund
$ samtools index* ..*/samfiles/Wet2014.bam
$ samtools depth -r k141_41392:152403-165349* ..*/samfiles/Wet2014.bam > Wet2014.cgc.depth.txt
$ cd* ..
~~~

The parameter `-r k141_41392:152403-165349` specifies a region in a contig. For any CGC, its positional range can be found in the file `cgc_standard.out` produced by run_dbcan (**Box 6**). The `depth.txt` files contain the raw read counts for the specified region.

**! CAUTION**

The contig IDs are automatically generated by MEGAHIT. There is a small chance that a same contig ID appears in both samples. However, the two contigs in the two samples do not match each other even the ID is the same. For example, the contig ID `k141_4139` is most likely only found in the Wet2014 sample. Even if there is a `k141_41392` in Dry2014, the actual contigs in two samples are different.

##### P13 dbcan_utils to calculate the abundance of CAZyme families, subfamilies, CGCs, and substrates (TIMING ∼1min)

~~~
*$ dbcan_utils fam_abund -bt Wet2014.depth.txt -i* ..*/Wet2014.dbCAN -a TPM
$ dbcan_utils fam_substrate_abund -bt Wet2014.depth.txt -i* ..*/Wet2014.dbCAN -a TPM
$ dbcan_utils CGC_abund -bt Wet2014.depth.txt -i* ..*/Wet2014.dbCAN -a TPM
$ dbcan_utils CGC_substrate_abund -bt Wet2014.depth.txt -i* ..*/Wet2014.dbCAN -a TPM
$ cd* .. *&& cd Dry2014_abund
$ dbcan_utils fam_abundfam_substrate_abund -bt Dry2014.depth.txt -i* ..*/Dry2014.dbCAN -a TPM
$ dbcan_utils fam_substrate_abund -bt Dry2014.depth.txt -i* ..*/Dry2014.dbCAN -a TPM
$ dbcan_utils CGC_abund -bt Dry2014.depth.txt -i* ..*/Dry2014.dbCAN -a TPM
$ dbcan_utils CGC_substrate_abund -bt Dry2014.depth.txt -i* ..*/Dry2014.dbCAN -a TPM
cd* ..
~~~

We developed a set of Python scripts as `dbcan_utils` (included in the run_dbcan package) to take the raw read counts for all genes as input and output the normalized abundances (**Box 7**) of CAZyme families, subfamilies, CGCs, and substrates (**Fig. 4**). The parameter `-a TPM` can also be two other metrics: RPM, or RPKM^61^.

RPKM = # of mapped reads to a gene G / [(total # of mapped reads to all genes /10^6^) x (gene G length/1000)]

RPM = # of mapped reads to a gene G / (total # of mapped reads to all genes/10^6^).

TPM = [# of mapped reads to a gene G / (gene G length/1000)] / sum [# of mapped reads to each gene / (the gene length/1000)].

###### Box 7

**Example output of dbcan_utils**

As an example, the Wet2014_abund folder

(https://bcb.unl.edu/dbCAN_tutorial/dataset1-Carter2023/individual_assembly/Wet2014_abund/) has 7 TSV files:

~~~
*-rw-rw-r-- 1 jinfang jinfang 201106 Dec 31 01:58 CGC_abund.out
-rw-rw-r-- 1 jinfang jinfang 2204 Dec 31 01:58 CGC_substrate_majority_voting.out
-rw-rw-r-- 1 jinfang jinfang 16282 Dec 31 01:58 CGC_substrate_PUL_homology.out
-rw-rw-r-- 1 jinfang jinfang 2695 Dec 31 01:58 EC_abund.out
-rw-rw-r-- 1 jinfang jinfang 3949 Dec 31 01:58 fam_abund.out
-rw-rw-r-- 1 jinfang jinfang 44138 Dec 31 01:58 fam_substrate_abund.out
-rw-rw-r-- 1 jinfang jinfang 27314 Dec 31 01:58 subfam_abund.out
-rw-rw-r-- 1 jinfang jinfang 270535 Dec 31 02:43 Wet2014.cgc.depth.txt*
~~~

Explanation of columns in these TSV files is as follows:

**fam_abund**.**out**: CAZy family (from HMMER vs dbCAN HMMdb), sum of TPM, # of CAZymes in the family

**subfam_abund**.**out:** eCAMI subfamily (from HMMER vs dbCAN-sub HMMdb), sum of TPM, # of CAZymes in the subfamily

**EC_abund**.**out:** EC number (extracted from dbCAN-sub subfamily), sum of TPM, # of CAZymes with the EC

**fam_substrate_abund**.**out:** Substrate (from HMMER vs dbCAN-sub HMMdb), sum of TPM (all CAZymes in this substrate group), GeneID (all CAZyme IDs in this substrate group)

**CGC_abund**.**out:** CGC_ID (e.g., k141_338400|CGC1), mean of TPM (all genes in the CGC), Seq_IDs (IDs of all genes in the CGC), TPM (of all genes in the CGC), Families (CAZyme family or other signature gene type of all genes in the CGC)

**CGC_substrate_PUL_homology**.**out:** Substrate (from dbCAN-PUL blast search), sum of TPM, CGC_IDs (all CGCs predicted to have the substrate from dbCAN-PUL blast search), TPM (of CGCs in this substrate group)

**CGC_substrate_majority_voting**.**out:** Substrate (from dbCAN-sub majority voting), sum of TPM, CGC_IDs (all CGCs predicted to have the substrate from dbCAN-sub majority voting), TPM (of CGCs in this substrate group)

**! CAUTION**

As shown in **Fig. 3** (step3), proteins from multiple samples can be combined to generate a non-redundant set of proteins (**Box 8**). This may reduce the runtime for the run_dbcan step (step4), as only one `faa` file will be processed. However, this does not work for the CGC prediction, as contigs (`fna` files) from each sample will be needed. Therefore, this step is recommended if users only want the CAZyme annotation, and not recommended if CGCs are also to be predicted.

###### Box 8

**Combine proteins from multiple samples (optional TIMING ∼3h)**

This protein sequence clustering step will create a mapping table with sequence cluster ID and protein IDs from each sample.

~~~
*$ mkdir mmseqs_cluster && cd mmseqs_cluster
$ ln -s* ..*/db* .
*$ cat* ..*/prokka_Wet2014/Wet2014.faa* ..*/prokka_Dry2014/Dry2014.faa > Dry_Wet.faa
$ mmseqs easy-cluster --threads 32 -c 0.95 --min-seq-id 0.95 --cov-mode 2 Dry_Wet.faa Dry_Wet_cluster tmp
$ mv Dry_Wet_cluster_cluster_rep.fasta Dry_Wet.cluster.faa*
~~~

This `Dry_Wet.cluster.faa` file now contains the Fasta sequences of representative proteins of all mmseqs2 clusters, i.e., the non-redundant protein sequences from the two samples. The mapping table file `Dry_Wet_cluster_cluster.tsv` contains two columns, mmseqs2 cluster representative protein ID, and protein IDs in the cluster. Users can query this table to find the correspondence between CAZymes in Wet2014 and in Dry2014.

#### Module 4: dbcan_plot for data visualization (Fig. 3) of abundances of CAZymes, CGCs, and substrates (TIMING variable)

##### CRITICAL STEP

To visualize the CAZyme annotation result, we provide a set of Python scripts as `dbcan_plot` to make publication quality plots with the dbcan_utils results as the input. The `dbcan_plot` scripts are included in the run_dbcan package. Once the plots are made in PDF format, they can be transferred to users’ Windows or Mac computers for visualization.

Five data folders will be needed as the input for `dbcan_plot`: (i) two abundance folders `Wet2014_abund` and `Dry2014_abund`, (ii) two CAZyme annotation folders `Wet2014.dbCAN` and `Dry2014.dbCAN`, and (iii) the `dbCAN-PUL` folder (under the `db` folder, released from `dbCAN-PUL.tar.gz`).

###### P14 Heatmap for CAZyme substrate abundance across samples (Fig. 6A) (TIMING 1min)

~~~
*$ dbcan_plot heatmap_plot --samples Wet2014,Dry2014 -i Wet2014_abund/
fam_substrate_abund.out,Dry2014_abund/ fam_substrate_abund.out --show_abund --top 20*
~~~

Here we plot the top 20 substrates in the two samples (**Fig. 6A**). The input files are the two CAZyme substrate abundance files calculated based on dbCAN-sub result. The default heatmap is ranked by substrate abundances. To rank the heatmap according to abundance profile using the clustermap function of the seaborn package (https://github.com/mwaskom/seaborn), users can invoke the `--cluster_map` parameter.

**Fig. 6:**
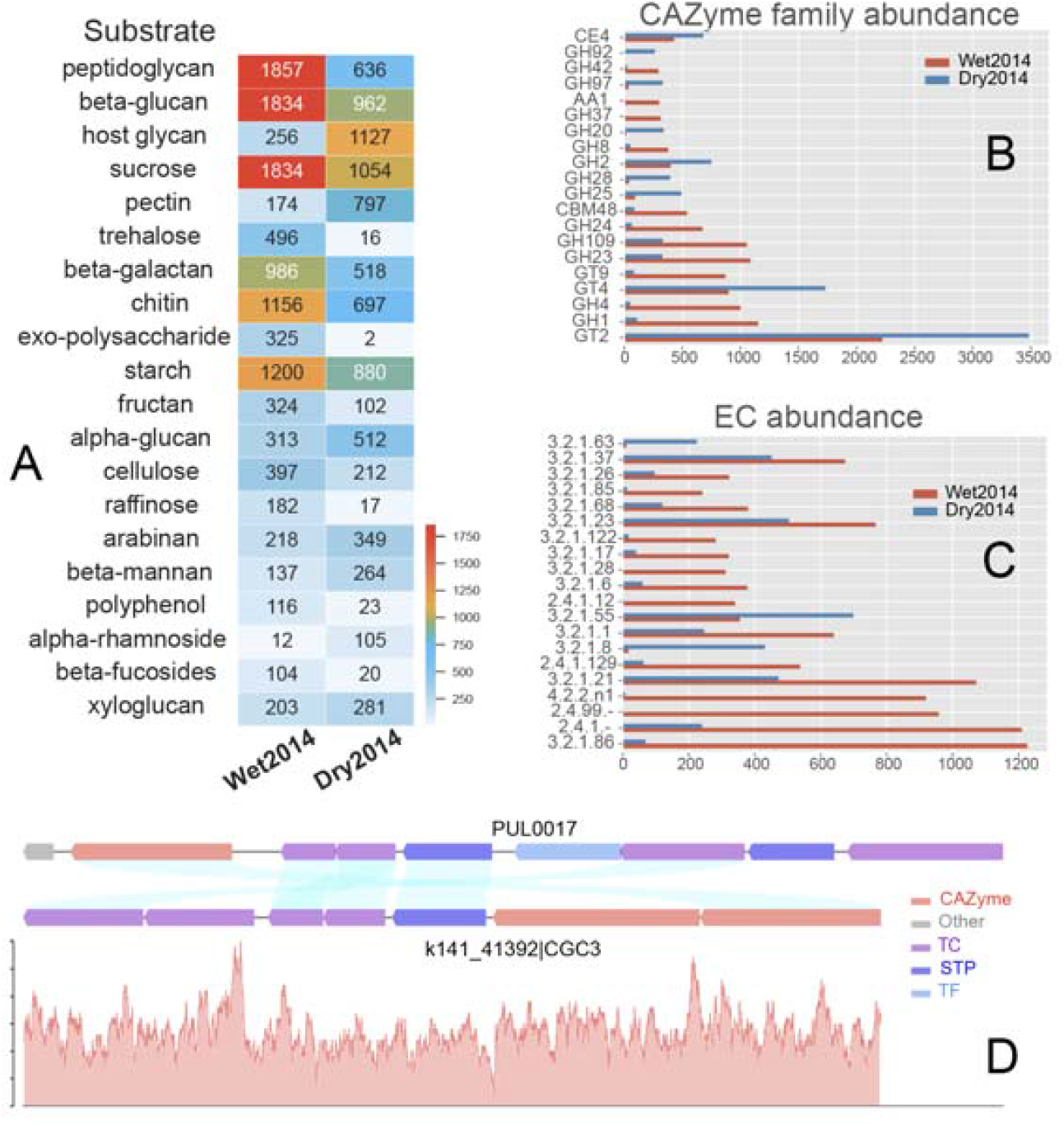
Data visualization of abundances of CAZymes, CGCs, and substrates. (**A**) Heatmap of CAZyme substrate abundance in the two samples (Carter2023 dataset: see Table 2) calculated based on dbCAN-sub search results and substrate mapping. Abundance values are TPM. (**B**) Barplot of 20 CAZyme family abundance in the two samples calculated based on dbCAN search results. (**C**) Barplot of 20 CAZyme EC abundance in the two samples calculated based on dbCAN-sub search results. (**D**) Synteny plot between an example CGC (CGC3 of contig k141_41396 from the Wet2014 sample) and its best PUL hit (PUL0017 from dbCAN-PUL, with experimentally verified substrate cellobiose ^76^) with read mapping coverage plot (y-axis is the read depth) shown in the bottom.

###### P15 Barplot for CAZyme family/subfamily/EC abundance across samples (Fig. B,C) (TIMING 1min)

~~~
*$ dbcan_plot bar_plot --samples Wet2014,Dry2014 --vertical_bar --top 20 -i
Wet2014_abund/fam_abund.out,Dry2014_abund/fam_abund.out
dbcan_plot bar_plot --samples Wet2014,Dry2014 --vertical_bar --top 20 -i
Wet2014_abund/subfam_abund.out,Dry2014_abund/subfam_abund.out*
~~~

Users can choose to generate a barplot instead of heatmap using the `bar_plot` method.

###### P16 Synteny plot between a CGC and its best PUL hit with read mapping coverage to CGC (Fig. 6D) (TIMING 1min)

~~~
*$ dbcan_plot CGC_synteny_coverage_plot -i Wet2014.dbCAN --cgcid ‘k141_41392*|*CGC3’ --readscount
Wet2014_abund/Wet2014.cgc.depth.txt*
~~~

The `Wet2014.dbCAN` folder contains the `PUL.out` file. Using this file, the `cgc_standard.out` file, and the best PUL’s `gff` file in `dbCAN-PUL.tar.gz`, the `CGC_synteny_plot` method will create the CGC-PUL synteny plot. The `–cgcid` parameter is required to specify which CGC to plot (`k141_41392|CGC3` in this example). The `Wet2014.cgc.depth.txt` file is used to plot the read mapping coverage.

If users only want to plot the CGC structure:

~~~
*$ dbcan_plot CGC_plot -i Wet2014.dbCAN --cgcid ‘k141_41392*|*CGC3’*
~~~

If users only want to plot the CGC structure plus the read mapping coverage:

~~~
*$ dbcan_plot CGC_coverage_plot -i Wet2014.dbCAN --cgcid ‘k141_41392*|*CGC3’ --readscount Wet2014_abund/Wet2014.cgc.depth.txt*
~~~

If users only want to plot the synteny between the CGC and PUL:

~~~
*$ dbcan_plot CGC_synteny_plot -i Wet2014.dbCAN --cgcid ‘k141_41392*|*CGC3’*
~~~

**! CAUTION**

The CGC IDs in different samples do not match each other. For example, specifying `-i Wet2014.dbCAN` is to plot the `‘k141_41392|CGC3’` in the Wet2014 sample. The `‘k141_41392|CGC3’` in the Dry2014 sample most likely does not exist, and even it does, the CGC has a different sequence even if the ID is the same.

## Troubleshooting

We provide **Table 3** to list possible issues and solutions. Users can also post issues on run_dbcan GitHub site.

**Table 3.**
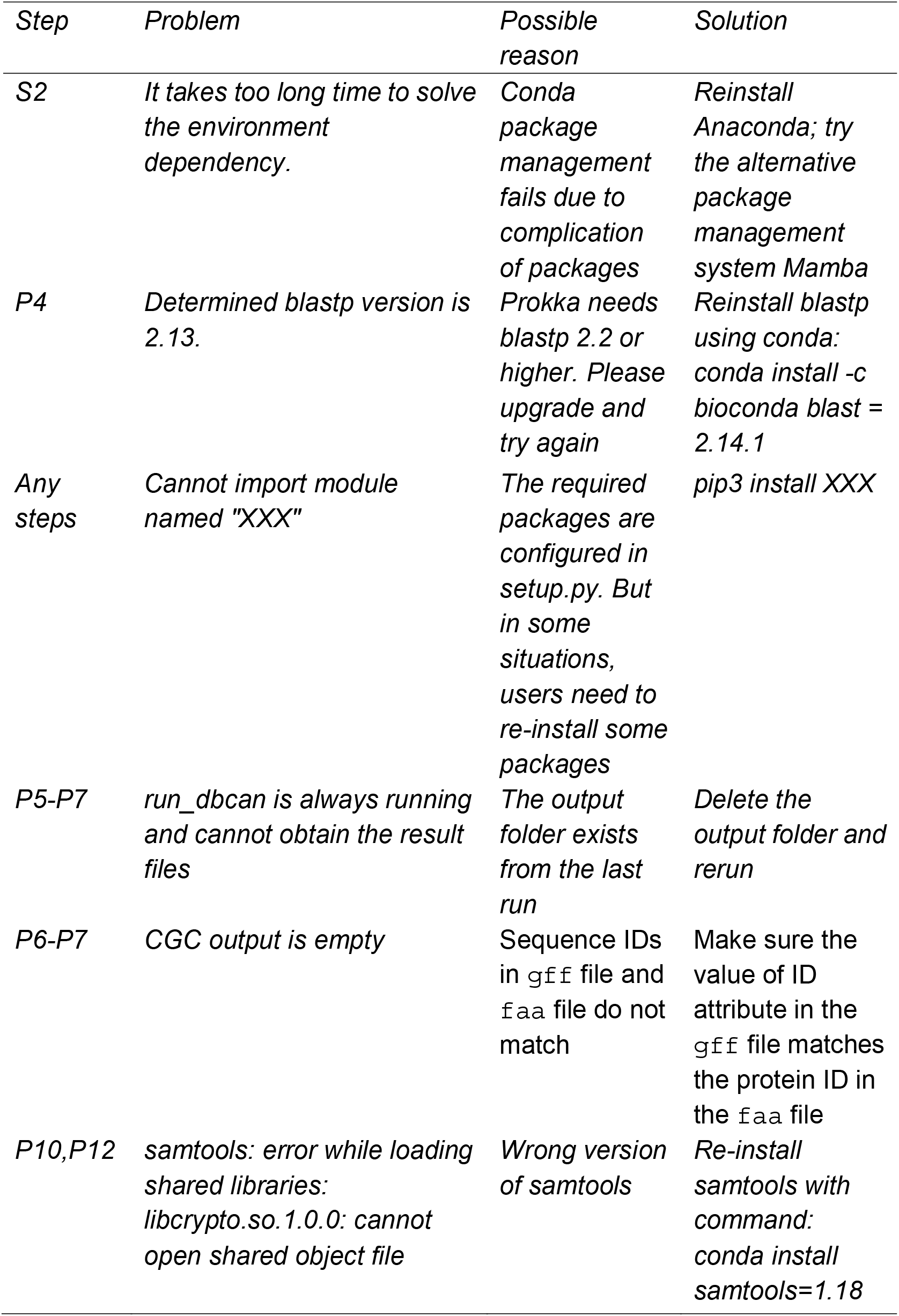
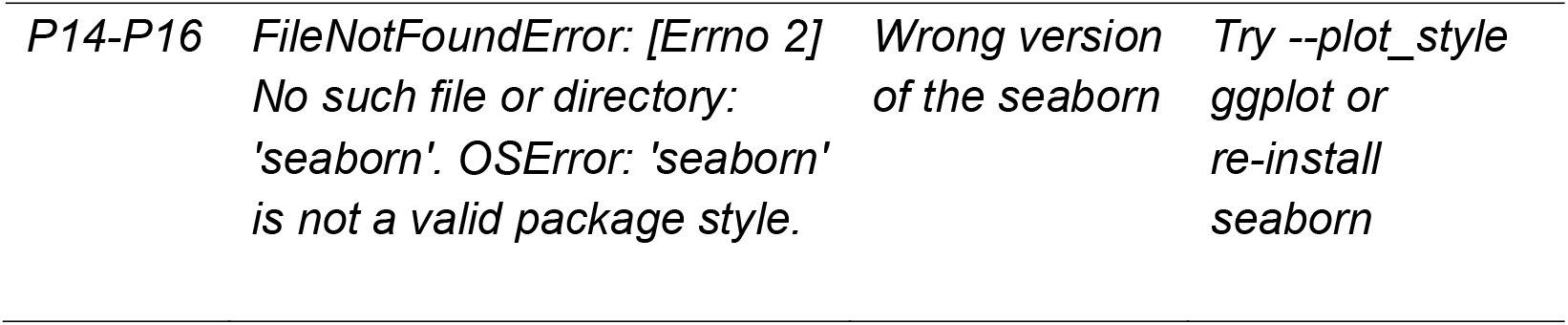
Troubleshooting table.

| *Step* | *Problem* | *Possible reason* | *Solution* |
| --- | --- | --- | --- |
| *S2* | *It takes too long time to solve the environment dependency.* | *Conda package management fails due to complication of packages* | *Reinstall Anaconda; try the alternative package management system Mamba* |
| *P4* | *Determined blastp version is 2.13.* | *Prokka needs blastp 2.2 or higher. Please upgrade and try again* | *Reinstall blastp using conda: conda install -c bioconda blast = 2.14.1* |
| *Any steps* | *Cannot import module named "XXX"* | *The required packages are configured in setup.py. But in some situations, users need to re-install some packages* | *pip3 install XXX* |
| *P5-P7* | *run_dbcan is always running and cannot obtain the result files* | *The output folder exists from the last run* | *Delete the output folder and rerun* |
| *P6-P7* | *CGC output is empty* | *Sequence IDs in gff file and faa file do not match* | *Make sure the value of ID attribute in the gff file matches the protein ID in the faa file* |
| *P10,P12* | *samtools: error while loading shared libraries: libcrypto.so.1.0.0: cannot open shared object file* | *Wrong version of samtools* | *Re-install samtools with command: conda install samtools=1.18* |
| *P14-P16* | *FileNotFoundError: [Errno 2]*<br>*No such file or directory:*<br>*'seaborn'. OSError: 'seaborn'*<br>*is not a valid package style.* | *Wrong version*<br>*of the seaborn* | *Try --plot_style*<br>*ggplot or*<br>*re-install*<br>*seaborn* |

## TIMING

Step P1. Contamination checking ∼10min

Step P2. Raw reads processing ∼20min

Step P3. Metagenome assembly ∼4h20min

Step P4. Gene models prediction ∼21h

Step P5. CAZyme annotation ∼10min

Step P6. PUL prediction ∼15min

Step P7. Substrate prediction both for CAZyme and PUL ∼5h

Step P8-P12. Reads mapping ∼52min

Step P13. Abundance estimation ∼1min

Step P14-P16. Data visualization ∼3min

Running this protocol on the Carter2023 dataset will take ∼33h on a Linux computer with 40 CPUs and 128GB of RAM. The most time-consuming step is P4 (Prokka gene prediction). Prodigal^59^ can be used to replace Prokka to only predict proteins, which will be significantly faster. The second time-consuming step is P7 (substrate prediction for CGCs and CAZymes). If users choose not to predict substates, this step will take ∼15min. RAM usage was not specifically monitored during the execution. The step with the highest RAM usage is likely P3 (read assembly).

## Anticipated results

Here we describe and compare the anticipated results for the Carter2023 dataset individual sample assembly route (protocol above in the main text), co-assembly route, and assembly-free route (protocols in **Supplementary Protocols, Fig. 3**).

### Assembled contigs and predicted proteins

The individual sample assembly route produced more contigs and proteins in the Dry2014 sample than in the Wet2014 sample (**Table 4**). This is unexpected as Dry2014 has only half amount of reads as Wet2014 (**Table 2**). Dry2014 also has a higher contig N50 and a higher percentage of unassembled reads. Carter2023 is an ultra-deep sequencing dataset of human gut microbiomes. Wet2014 was shown to have the highest *in situ* average growth rate while Dry2014 had the lowest^16^, suggesting a revival of lower abundant species (having lower growth rate) in Dry2014. An earlier study also found that Dry2014 had a higher phylogenetic diversity than Wet2014^53^. The higher species diversity might be able to explain the higher count and length of contigs, and the higher number of proteins in Dry2014 despite its lower number of reads. We also verified all these findings by sub-sampling an equal number of reads (20,000,000) from Wet2014 and Dry2014 (**Table 2**) and repeating the METAHIT assembly, Prokka gene prediction, and CAZyme occurrence and abundance calculation (**Supplementary Protocols**).

**Table 4.**
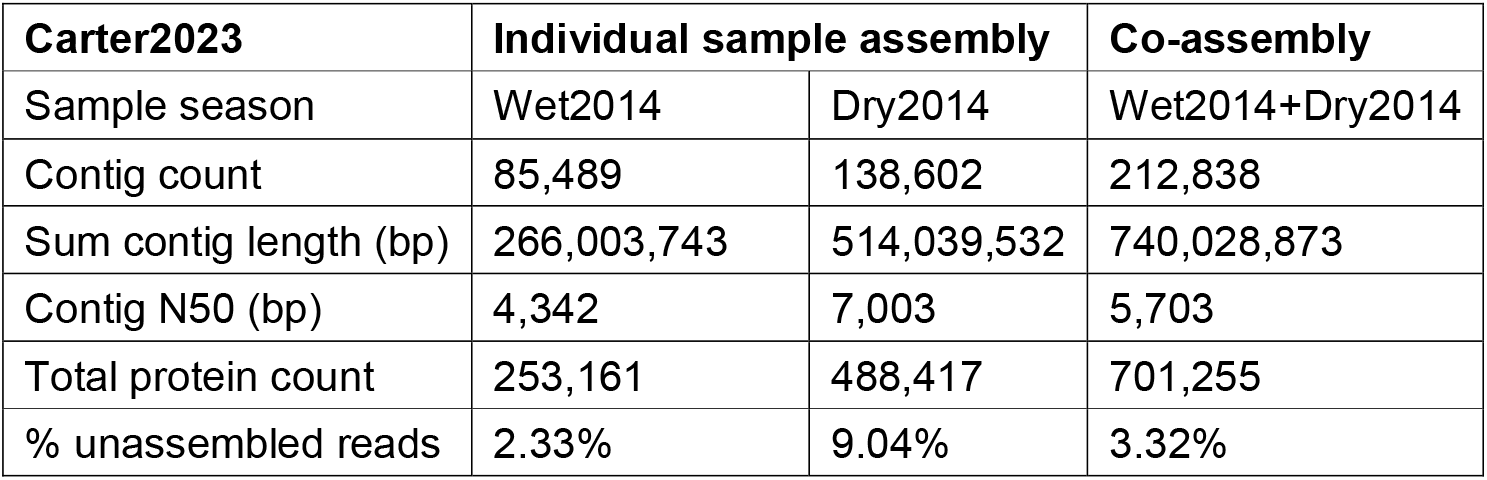
Comparison of contigs and proteins among the two routes.

As detailed in **Supplementary Protocols**, we also developed a protocol for the co-assembly route (**Fig. 3**), where reads of Wet2014 and Dry2014 were pooled together before all the analyses (all data including precomputed data and final results at: https://bcb.unl.edu/dbCAN_tutorial/dataset1-Carter2023/co_assembly/). A recent study found that co-assembly of 124 marine metagenomic samples led to a slightly higher total contig length but a lower number of proteins than individual assembly^55^. We found that for the Carter2023 dataset from human gut microbiome, the contig count, contig length, and protein count from co-assembly are close to the simple sum of the results from the two individual assemblies (**Table 4**). The percentage of unassembled reads for the co-assembly is 3.32%, closer to that of Wet2014 (2.33%) than Dry2014 (9.04%). This suggests that co-assembly has improved the assembly of contigs from the low abundant species.

As mentioned in **Box 8**, we also generated a non-redundant set of proteins for the individual sample assembly route by mmseqs2 sequence clustering (sequence identity > 95% and coverage > 95%). This produced 685,433 non-redundant proteins (from a total 741,578 proteins), lower than the co-assembly protein count 701,255. With the cluster IDs mapped to the protein sequence IDs in the two samples, we found Dry2014 (435,754) has more unique protein sequences than Wet2014 (203,633). The rest of proteins (102,191) are found in 46,046 mmseqs2 clusters that have proteins from both samples.

We further clustered all the 1,442,833 proteins from the three protein sets (co-assembly: 701,255, Dry2014: 488,417, Wet2014: 253,161) using mmseqs2. Co-assembly has 148,011 unique proteins that were missed by the individual sample assembly (**Fig. 7**). Co-assembly also recovered most proteins that are shared by Wet2014 and Dry2014, but missed 66,717 and 84,722 proteins that are unique to each sample (likely from low abundant and sample-specific species). The three-way clustering also confirmed that Dry2014 and Wet2014 only share a small percentage of proteins (under the threshold: identity > 95% and coverage > 95%), and Dry2014 has many more proteins than Wet2014 that were recovered by co-assembly. As detailed in **Experimental design** of Introduction, we have discussed about the benefits and limitations of co-assembly. With the contigs and proteins from the co-assembly, we also developed a protocol for Prokka gene prediction, CAZyme occurrence and abundance calculation (**Supplementary Protocols**).

**Fig. 7:**
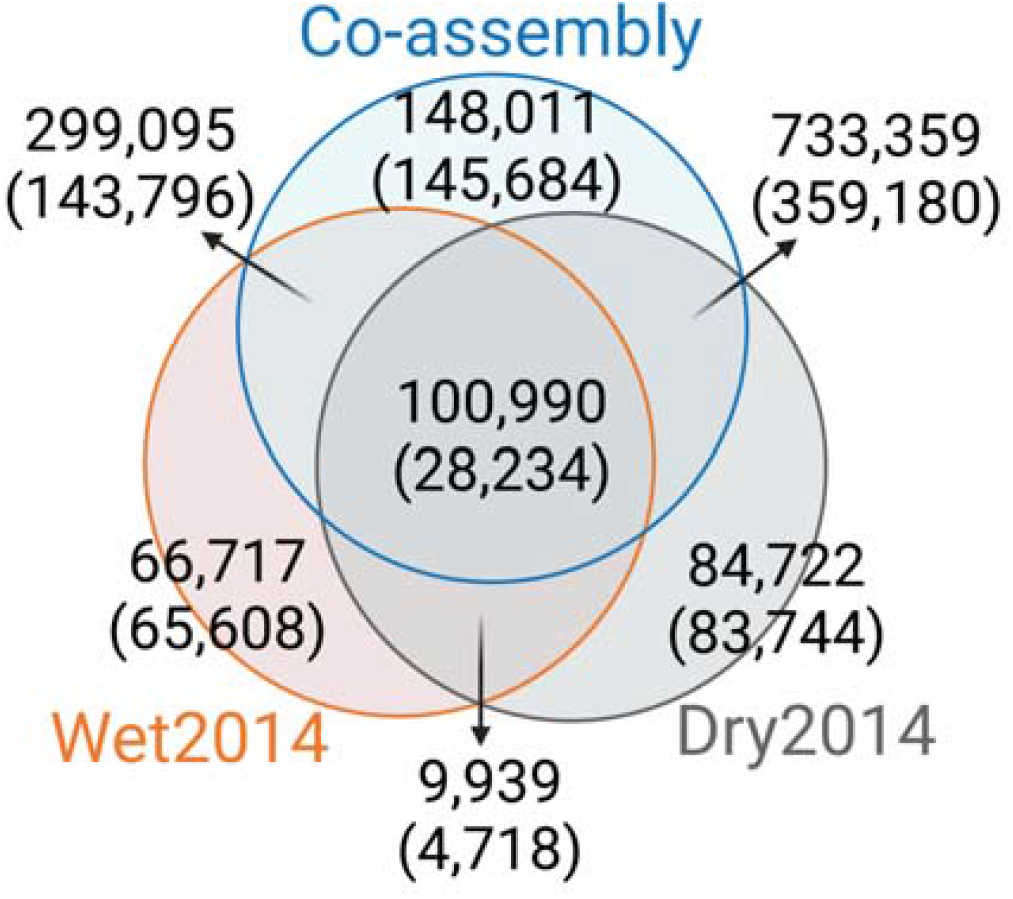
Mmseqs2 clustering of three protein sets. The numbers outside the parentheses are protein counts, while the numbers in parentheses are mmseqs2 cluster counts.

### Predicted CAZymes, CGCs, and glycan substrates

The key results of CAZyme annotation in microbiomes using this protocol are provided in **Table 5**. The two samples from the independent sample assembly route have the same percentage (3.50%) of CAZymes (CAZyme count in Table 5 / total protein count in Table 4). The co-assembly route has a higher percentage (3.56%) of CAZymes. These percentages are also higher than that in metagenome assembled genomes (MAGs) of the Unified Human Gastrointestinal Genome (UHGG) collection (2.55%^56^). This is expected because a majority of MAGs in UHGG are sequenced from the Western population^75^, while the Carter2023 data are from Hadza hunter-gatherers, who consume more dietary fibers^16^.

**Table 5.** CAZymes, CGCs, and substrates.

| **Carter2023** | **Individual sample assembly** |  | **Co-assembly** |
| --- | --- | --- | --- |
| Sample season | Wet2014 | Dry2014 | Wet2014+Dry2014 |
| CAZyme count | 9,017 | 17,217 | 24,974 |
| CAZyme percentage | 3.50% | 3.50% | 3.56% |
| CAZyme count with predicted substrates | 2,093<br>(23.21%) | 3,776<br>(21.93%) | 5,453 (21.83%) |
| CGC count | 1,487 | 2,565 | 3,832 |
| CGC count with predicted substrates from dbCAN-PUL | 696 (46.80%) | 1,006<br>(39.22%) | 1,576 (41.13%) |
| CGC count with predicted substrates from CAZyme majority voting | 71 (4.77%) | 97 (3.78%) | 147 (3.84%) |

For substrate prediction from dbCAN-sub (eCAMI subfamily + EC mapping to curated substrates from literature and CAZy, `fam-substrate-mapping-08012023.tsv`), Wet2014 has 23.21% of CAZymes with predicted substrates, slightly higher than Dry2014 (21.93%) and the co-assembly route (21.83%). For substrate prediction from dbCAN-PUL homology search, Wet2014 has 46.80% of CGCs with predicted substrates, moderately higher than Dry2014 (39.22%) and the co-assembly route (41.13%). These percentages are much higher than what has been reported in the UHGG collection (28.48%^56^), again expected as Hadza hunter-gatherers consume more dietary fibers than Western population. In agreement with our previous paper^12^, the CAZyme majority voting from dbCAN-sub result can only predict a small percentage of CAZymes for substrates.

We have further compared the abundance of each individual CAZyme family, EC number, and substrate across different routes. The complete results can be found in **Supplementary Table 1**, which has 6 sheets providing the TPM read abundances calculated by `dbcan_utils`. This table was filtered to select the most different CAZyme families, substrates, and EC numbers between the two samples to make the heatmaps in **Fig. 8**.

**Fig. 8:**
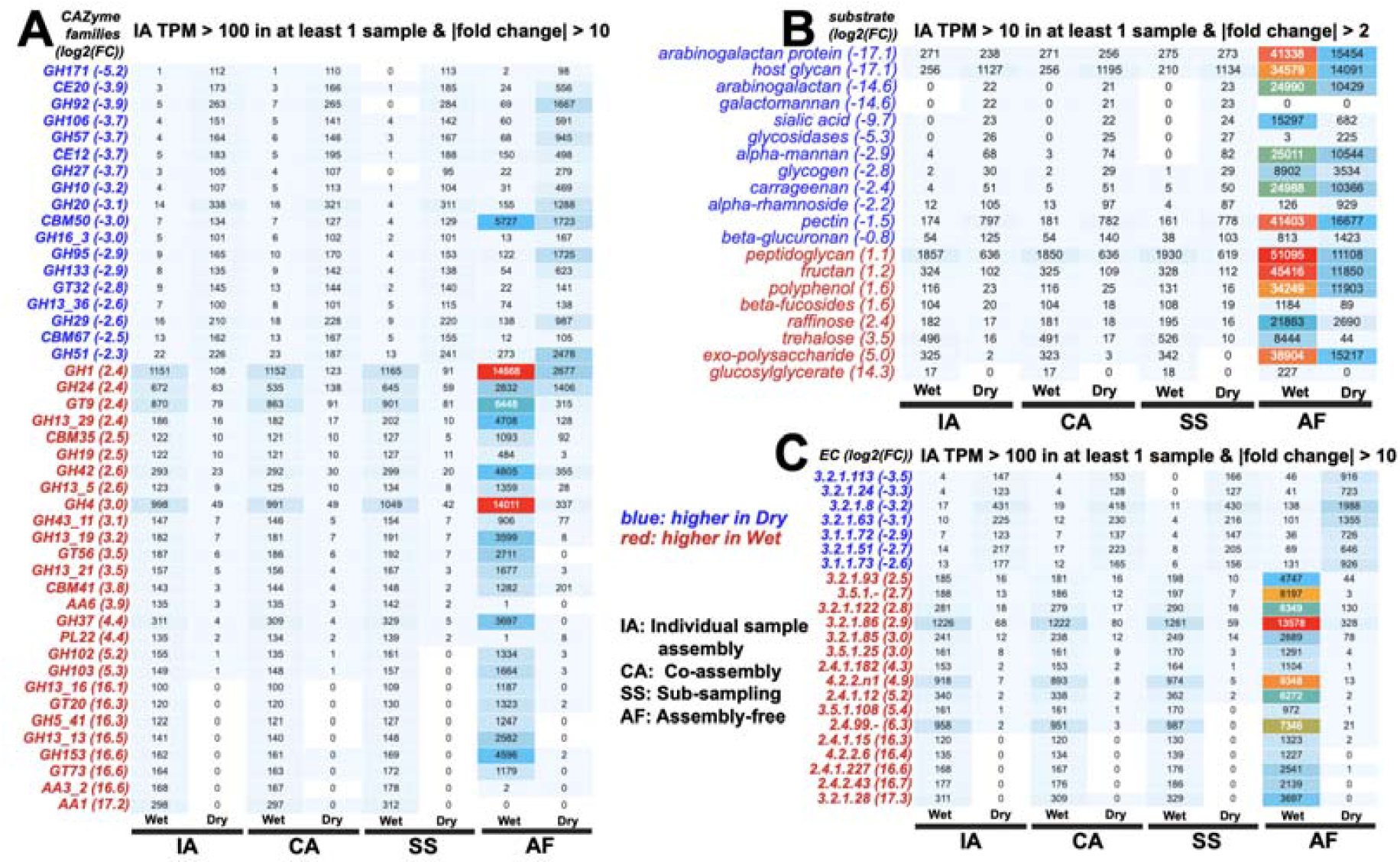
Comparisons of abundances calculated from different routes. (**A**) Abundance of CAZyme families. (**B**) Abundance of CAZyme substrates. (**C**) Abundance of EC numbers. Blue colors are used to indicate families, substrates, and ECs with higher abundances in Dry2014, while red colors are higher in Wet2014 sample, according to the IA (individual sample assembly) route. The values in parentheses are the log2(fold change) calculated for TPM in Wet2014 divided by TPM in Dry2014 from the IA route. Only those that meet the thresholds are shown: the TPM value and FC ratio thresholds are indicated on the top of each plot. The complete data are available in **Supplementary Table 1**.

It is interesting to note that the individual sample assembly route (with or without the sub-sampling) and co-assembly route produced surprisingly consistent results for all the families (**Fig. 8A**), substrates (**Fig. 8B**), and ECs (**Fig. 8C**). In contrast, the assembly-free route produced very different results. The assembly-free route uses DIAMOND blastx to map reads to CAZy annotated proteins (**Supplementary Protocols**), which, unlike assembly contigs, are not derived from the microbiome samples. Therefore, it is highly likely that a high number of false positive matches will be counted into the abundance calculation. Therefore, our result agrees with a recent paper conducted on metatranscriptomic data^58^ that the assembly-free approach is less reliable than assembly-based methods.

Thus, microbiome researchers including dbCAN users should abstain from using assembly-free methods for functional annotation whenever they can, although assembly-free methods are easier to perform and have lower computational demand. Given the fact that co-assembly is often not possible for of reads from multiple samples, we recommend users use the following order to select the suitable route following our protocols for CAZyme annotation of microbiome sequencing data: IA (or SS if sequencing depth is very high) > CA >> AF.

## Supporting information

Supplementary Protocols

## Data availability

https://bcb.unl.edu/dbCAN_tutorial/dataset1-Carter2023/

## Code availability

<u>Github:</u> https://github.com/linnabrown/run_dbcan

<u>Bioconda:</u> https://anaconda.org/bioconda/run-dbcan

<u>Docs:</u> https://dbcan.readthedocs.io

## Acknowledgements

This work was supported by the U.S. National Institutes of Health (NIH) awards [R01GM140370] and [R21AI171952], U.S. National Science Foundation (NSF) CAREER award [DBI-1933521], U.S. Department of Agriculture (USDA) award [58-8042-9-089]. Funding for open access charge: NIH award [R01GM140370]. This work was also partially completed utilizing the Holland Computing Center of the University of Nebraska, which receives support from the Nebraska Research Initiative.

## Author contributions

J.Z. developed the protocols, with help from other co-authors (Y.Yin, L.H., and H.Y.). X.Z., Y.Yan, and J.A. tested the protocol and provided help in troubleshooting. J.Z. is the first author of dbCAN3. L.H. is the main developer of run_dbcan GitHub and Bioconda packages. H.Y. is main developer of the run_dbcan Docker image. J.Z. and Y.Yin wrote the manuscript. Y.Yin supervised the entire project.

## Key references related to this protocol

*dbCAN: Y Yin et al. Nucleic Acids Res (2012). https://doi.org/10.1093/nar/gks479*.

*dbCAN2: H Zhang et al. Nucleic Acids Res (2018). https://doi.org/10.1093/nar/gky418*.

*dbCAN3: J Zheng et al. Nucleic Acids Res (2023). https://doi.org/10.1093/nar/gkad328*.

## Supplementary information

Supplementary Figs. 1–3, Supplementary Table 1, and Supplementary Protocols.

