## Supplementary Protocols for "Carbohydrate-active enzyme annotation in microbiomes using dbCAN"

**There are 5 sections in this document:**

- Procedure of the co-assembly route (**Carter2023**): pages 2-11
- Procedure of the assembly-free route (**Carter2023**): pages 12-15
- Procedure of the sub-sampling route (**Carter2023**): pages 16-22
- Procedure of the individual sample assembly route (**Wastyk2021**): pages 23-34
- Procedure of the individual sample assembly route (**Priest2023**): pages 35-45

References: pages 46-47

**Procedure of the co-assembly route (Carter2023)**

This is the procedure for the co-assembly route of dataset1–Carter2023.

The procedure has 4 modules (**Fig. 3**) and 16 steps (P1-P16). All the steps will be conducted in a separate folder named co-assembly.

*$ trim_galore --paired Dry2014_1.fastq.gz Dry2014_2.fastq.gz --illumina -j 36*

We specified --illumina to indicate that the reads were generated using the Illumina sequencing platform. Nonetheless, trim_galore possesses the ability to automatically detect the adapter, providing flexibility in adapter handling for users who may know the specific sequencing platform. Details of trimming are available in the trimming report file (**Box 2**).

**Box 2| Example output of trim_galore**

In addition to the trimmed read files, Trim_galore also generates a trimming report file. The trimming report contains details on read trimming, such as the number of trimmed reads.

*-rw-rw-r-- 1 jinfang jinfang 4.2K Dec 13 01:48 Dry2014_1.fastq.gz_trimming_report.txt*

*-rw-rw-r-- 1 jinfang jinfang 2.0G Dec 13 01:55 Dry2014_1_val_1.fq.gz*

*-rw-rw-r-- 1 jinfang jinfang 4.4K Dec 13 01:55 Dry2014_2.fastq.gz_trimming_report.txt*

*-rw-rw-r-- 1 jinfang jinfang 2.4G Dec 13 01:55 Dry2014_2_val_2.fq.gz*

*-rw-rw-r-- 1 jinfang jinfang 4.4K Dec 13 01:30 Wet2014_1.fastq.gz_trimming_report.txt*

*-rw-rw-r-- 1 jinfang jinfang 3.4G Dec 13 01:46 Wet2014_1_val_1.fq.gz*

*-rw-rw-r-- 1 jinfang jinfang 4.6K Dec 13 01:46 Wet2014_2.fastq.gz_trimming_report.txt*

*-rw-rw-r-- 1 jinfang jinfang 3.7G Dec 13 01:46 Wet2014_2_val_2.fq.gz*

**! CAUTION**

During the trimming process, certain reads may be entirely removed due to low quality in its entirety. Using the --retain_unpaired parameter in trim_galore allows for the preservation of single-end reads. In this protocol, this option was not select, so that both reads of a forward-revise pair were removed.

**P3| Co-assemble reads into contigs (TIMING ~4h30min)**

*$ megahit --min-contig-len 1000 -1 Dry2014_1_val_1.fq.gz, Wet2014_1_val_1.fq.gz -2 Dry2014_2_val_2.fq.gz, Wet2014_2_val_2.fq.gz -m 0.5 -t 32 -o megahit_Hadza_INDIV_157 --out-prefix Hadza_INDIV_157*

MEGAHIT generates folder megahit_Hadza_INDIV_157 containing five files and one sub-folder (**Box 3**). Hadza_INDIV_157.contigs.fa is the final contig sequence file. We set --min-contig-len 1000, a common practice to retain all contigs longer than 1,000 base pairs. The commands are slightly different from the individual sample assembly route (main text) for this step.

**Box 3| Example output of megahit**

```
-rw-rw-r-- 1 jinfang jinfang  262 Dec 20 06:21 checkpoints.txt
-rw-rw-r-- 1 jinfang jinfang    0 Dec 20 06:21 done
-rw-rw-r-- 1 jinfang jinfang 715M Dec 20 06:21 Hadza_INDIV_157.contigs.fa
-rw-rw-r-- 1 jinfang jinfang 240K Dec 20 06:21 Hadza_INDIV_157.log
drwxrwxr-x 2 jinfang jinfang 4.0K Dec 20 06:21 intermediate_contigs
-rw-rw-r-- 1 jinfang jinfang 1.4K Dec 20 01:52 options.json
```

**P4| Predict genes by Prokka (TIMING 40h42min)**

*$ prokka --outdir prokka_Hadza_INDIV_157 --prefix Hadza_INDIV_157 --addgenes --addmrna --locustag Hadza_INDIV_157 --kingdom Bacteria --cpus 36 megahit_Hadza_INDIV_157/Hadza_INDIV_157.contigs.fa*

The parameter --kingdom Bacteria is required for bacterial gene prediction. To optimize performance, --CPU 36 instructs the utilization of 36 computer processors. The output files comprise of both protein and CDS sequences in Fasta format (e.g., Hadza_INDIV_157.faa and Hadza_INDIV_157.ffn in **Box 4**).

**Box 4| Example output of Prokka**

```
-rw-rw-r-- 1 jinfang jinfang  28M Dec 21 23:02 Hadza_INDIV_157.err
-rw-rw-r-- 1 jinfang jinfang 215M Dec 20 10:50 Hadza_INDIV_157.faa
-rw-rw-r-- 1 jinfang jinfang 580M Dec 20 10:50 Hadza_INDIV_157.ffn
-rw-rw-r-- 1 jinfang jinfang 721M Dec 20 06:21 Hadza_INDIV_157.fna
-rw-rw-r-- 1 jinfang jinfang 731M Dec 20 10:50 Hadza_INDIV_157.fsa
-rw-rw-r-- 1 jinfang jinfang 1.7G Dec 21 23:04 Hadza_INDIV_157.gbk
-rw-rw-r-- 1 jinfang jinfang 1.1G Dec 20 10:50 Hadza_INDIV_157.gff
-rw-rw-r-- 1 jinfang jinfang 5.4M Dec 21 23:04 Hadza_INDIV_157.log
-rw-rw-r-- 1 jinfang jinfang 3.1G Dec 21 23:03 Hadza_INDIV_157.sqn
-rw-rw-r-- 1 jinfang jinfang 201M Dec 20 10:50 Hadza_INDIV_157.tbl
-rw-rw-r-- 1 jinfang jinfang  99M Dec 20 10:50 Hadza_INDIV_157.tsv
-rw-rw-r-- 1 jinfang jinfang  155 Dec 20 10:50 Hadza_INDIV_157.txt
```

**Module 2: run_dbcan annotation (Fig. 3) to obtain CAZymes, CGCs, and substrates**

**P5| CAZyme annotation at family level (TIMING ~7min)**

*$ run_dbcan prokka_Hadza_INDIV_157/Hadza_INDIV_157.faa protein --hmm_cpu 32 --tools hmmer --out_dir Hadza_INDIV_157.CAZyme*

Two arguments are required for run_dbcan: the input sequence file (faa files) and the sequence type (protein). By default, run_dbcan will use three methods (HMMER vs dbCAN HMMdb, DIAMOND vs CAZy, HMMER vs dbCAN-sub HMMdb) for CAZyme annotation (**Table 1, Fig. 2**). This default setting is equivalent to the use *--tools all* parameter (**Box 5**). Here we only invoke the HMMER vs dbCAN HMMdb for CAZyme annotation at the family level.

**Box 5| CAZyme annotation with default setting**

If the *--tools* parameter is not set, it is the default setting, which is the same as *--tools all*. This will take much longer time to finish (~5h) due to the large size of dbCAN-sub HMMdb (used for substrate prediction for CAZymes, see **Table 1**).

*$ run_dbcan prokka_Hadza_INDIV_157/Hadza_INDIV_157.faa protein --hmm_cpu 32 --tools all --out_dir Hadza_INDIV_157.CAZyme*

The sequence type can be protein, prok, meta. If the input sequence file contains metagenomic contig sequences (fna file), the sequence type has to be meta, and Prodigal will be called to predict genes.

*$ run_dbcan prokka_Hadza_INDIV_157/Hadza_INDIV_157.fna meta --out_dir Hadza_INDIV_157.CAZyme --dia_cpu 32 --hmm_cpu 32 --dbcan_thread 32*

**P6| CGC prediction (TIMING ~15 min)**

The following commands will re-run run_dbcan to not only predict CAZymes but also CGCs with protein faa and gene location gff files.

*$ run_dbcan prokka_Hadza_INDIV_157/Hadza_INDIV_157.faa protein --tools hmmer --tf_cpu 32 --stp_cpu 32 -c prokka_Hadza_INDIV_157/Hadza_INDIV_157.gff --out_dir Hadza_INDIV_157.PUL --dia_cpu 32 --hmm_cpu 32*

As mentioned above (**Table 1, Fig. 2**), CGC prediction is a featured function added into dbCAN2 in 2018. To identify CGCs with the protein sequence type, a gene location file (gff) must be provided together. If the input sequence type is prok or meta, meaning users only have contig fna files, the CGC prediction can be activated by setting *-c cluster*.

**! CAUTION**

If the users would like to create their own gff file (instead of using Prokka or Prodigal), it is important to make sure the value of ID attribute in the gff file matches the protein ID in the protein faa file.

**?Troubleshooting**

If no result is found in CGC output file, it is most likely because the sequence IDs in gff file and faa file do not match. Another less likely reason is that the contigs are too short and fragmented and not suitable for CGC prediction.

**P7| Substrate prediction for CAZymes and CGCs (TIMING ~5h)**

The following commands will re-run run_dbcan to predict CAZymes, CGCs, and their substrates with the *--cgc_substrate* parameter.

*$ run_dbcan prokka_Hadza_INDIV_157/Hadza_INDIV_157.faa protein --dbcan_thread 32 --tf_cpu 32 --stp_cpu 32 -c prokka_Hadza_INDIV_157/Hadza_INDIV_157.gff --cgc_substrate --out_dir Hadza_INDIV_157.dbCAN --hmm_cpu 32 --dia_cpu 32*

**! CAUTION**

The above commands do not set the *--tools* parameter, which means all three methods for CAZyme annotation will be activated (**Box 5**). Because dbCAN-sub HMMdb (for CAZyme substrate prediction) is 200 times larger than dbCAN HMMdb, the runtime will be much longer. Users can specify --tools hmmer, so that the HMMER search against dbCAN-sub will be disabled. However, this will turn off the substrate prediction for CAZymes and CGCs based on CAZyme substrate majority voting. Consequently, the substrate prediction will be solely based on homology search against PULs in dbCAN-PUL (**Fig. 1, Table 1**).

*$ run_dbcan prokka_Hadza_INDIV_157/Hadza_INDIV_157.faa protein --dbcan_thread 32 --tf_cpu 32 --stp_cpu 32 -c prokka_Hadza_INDIV_157/Hadza_INDIV_157.gff --cgc_substrate --out_dir Hadza_INDIV_157.dbCAN --hmm_cpu 32 --dia_cpu 32 –-tools hmmer*

**Box 6| Example output folder content of run_dbcan substrate prediction**

In the output directory *Hadza_INDIV_157.dbCAN*, a total of 17 files and 1 folder are generated:

```
-rw-rw-r-- 1 jinfang jinfang 8.9M Dec 22 07:45 CGC.faa
-rw-rw-r-- 1 jinfang jinfang  54M Dec 22 07:45 cgc.gff
-rw-rw-r-- 1 jinfang jinfang 2.5M Dec 22 07:45 cgc.out
-rw-rw-r-- 1 jinfang jinfang 1.1M Dec 22 07:45 cgc_standard.out
-rw-rw-r-- 1 jinfang jinfang 4.7M Dec 22 07:45 cgc_standard.out.json
-rw-rw-r-- 1 jinfang jinfang 2.3M Dec 22 07:35 dbcan-sub.hmm.out
-rw-rw-r-- 1 jinfang jinfang 1.5M Dec 22 07:35 diamond.out
-rw-rw-r-- 1 jinfang jinfang 1.9M Dec 22 07:35 dtemp.out
-rw-rw-r-- 1 jinfang jinfang 1.3M Dec 22 07:35 hmmer.out
-rw-rw-r-- 1 jinfang jinfang 1.3M Dec 22 07:45 overview.txt
-rw-rw-r-- 1 jinfang jinfang  85M Dec 22 07:48 PUL_blast.out
-rw-rw-r-- 1 jinfang jinfang 7.6M Dec 22 07:44 stp.out
-rw-rw-r-- 1 jinfang jinfang 151K Dec 22 07:48 substrate.out
drwxrwxr-x 2 jinfang jinfang  92K Dec 22 07:54 syntenic.pdf
-rw-rw-r-- 1 jinfang jinfang 2.2M Dec 22 07:38 tf-1.out
-rw-rw-r-- 1 jinfang jinfang 1.8M Dec 22 07:41 tf-2.out
-rw-rw-r-- 1 jinfang jinfang 6.0M Dec 22 07:44 tp.out
-rw-rw-r-- 1 jinfang jinfang 215M Dec 21 23:41 uniInput
```

**PUL_blast.out**: BLAST results between CGCs and PULs.

**CGC.faa**: Protein Fasta sequences encoded in all CGCs.

**cgc.gff**: reformatted from the user input gff file by marking CAZymes, TFs, TCs, and STPs.

**cgc.out**: raw output of CGC predictions.

**cgc_standard.out**: simplified version of cgc.out for easy parsing in TSV format. An example row has the following columns:

1. CGC_id: *CGC1*
2. type: *CAZyme*
3. contig_id: *k141_372285*
4. gene_id: *Hadza_INDIV_157_00546*
5. start: *3067*
6. end: *4389*
7. strand: *+*
8. annotation: *GH32*

*Explanation: the gene Hadza_INDIV_157_00546 encodes a GH32 CAZyme in the CGC1 of the contig k141_372285. CGC1 also has other genes, which are provided in other rows. Hadza_INDIV_157_00546 is on the positive strand of k141_372285 from 3067 to 4389. The type can be one of the four signature gene types (CAZymes, TCs, TFs, STPs) or the null type (not annotated as one of the four signature genes).*

**cgc_standard.out.json**: JSON format of cgc_standard.out.

**dbcan-sub.hmm.out**: HMMER search result against dbCAN-sub HMMdb, including a column with CAZyme substrates extracted from *fam-substrate-mapping-08012023.tsv*.

**diamond.out**: DIAMOND search result against the CAZy annotated protein sequences (*CAZyDB.07262023.fa*).

**dtemp.out**: temporary file.

**hmmer.out**: HMMER search result against dbCAN HMMdb.

**overview.txt**: summary of CAZyme annotation from three methods in TSV format. An example row has the following columns:

1. Gene_ID: *Hadza_INDIV_157_00661*
2. EC#: *2.4.1.25:3*
3. dbCAN: *GH77(11-489)*
4. dbCAN_sub: *GH77_e1*
5. DIAMOND: *GH77*
6. #ofTools: *3*

*Explanation: the protein Hadza_INDIV_157_00661 is annotated by 3 tools to be a CAZyme: (1) GH77 (CAZy defined family GH77) by HMMER vs dbCAN HMMdb with a domain range from aa position 11 to 489, (2) GH77_e1 (eCAMI defined subfamily e1; e indicates it is from eCAMI not CAZy) by HMMER vs dbCAN-sub HMMdb (derived from eCAMI subfamilies), and (3) GH77 by DIAMOND vs CAZy annotated protein sequences. The second column 2.4.1.25:3 is extracted from eCAMI, meaning that the eCAMI subfamily GH77_e1 contains three member proteins which have an EC 2.4.1.25 according to CAZy. In most cases, the 3 tools will have the same CAZyme family assignment. When they give different assignment. We recommend a preference order: dbCAN > eCAMI/dbCAN-sub > DIAMOND. See our dbCAN2 paper^1^, dbCAN3 paper^2^, and eCAMI^3^ for more details.*

*Note: If users invoked the --use_signalP parameter when running run_dbcan, there will be an additional column called signalP in the overview.txt.*

**stp.out**: HMMER search result against the MiST^4^ compiled signal transduction protein HMMs from Pfam.

**tf-1.out**: HMMER search result against the DBD^5^ compiled transcription factor HMMs from Pfam^6^.

**tf-2.out**: HMMER search result against the DBD compiled transcription factor HMMs from Superfamily^7^.

**tp.out**: DIAMOND search result against the TCDB^8^ annotated protein sequences.

**substrate.out**: summary of substrate prediction results for CGCs in TSV format from two approaches^2^ (dbCAN-PUL blast search and dbCAN-sub majority voting). An example row has the following columns:

1. CGC_ID: *k141_597014|CGC1*
2. Best hit PUL_ID in dbCAN-PUL: *PUL0484*
3. Substrate of the hit PUL: *pectin*
4. Sum of bitscores for homologous gene pairs between CGC and PUL: *1363.0*
5. Types of homologous gene pairs: *TC-TC;TC-TC;CAZyme-CAZyme;CAZyme-CAZyme;CAZyme-CAZyme*
6. Substrate predicted by majority voting of CAZymes in CGC: *xylan*
7. Voting score: *2.0*

*The CGC1 of contig k141_597014 has its best hit PUL0484 (from PUL_blast.out) with pectin as substrate (from dbCAN-PUL_12-12-2023.xlsx). Five signature genes are matched between k141_597014|CGC1 and PUL0484 (from PUL_blast.out): three are CAZymes and the others are TCs. The sum of blast bitscores of the five homologous pairs (TC-TC;TC-TC;CAZyme-CAZyme;CAZyme-CAZyme;CAZyme-CAZyme) is 1363.0. Hence, the substrate of k141_597014|CGC1 is predicted to be pectin according to dbCAN-PUL blast search. The last two columns are based on the dbCAN-sub result (dbcan-sub.hmm.out), according to which two CAZymes in k141_597014|CGC1 are predicted to have xylan substrate. The voting score is thus 2.0, so that according to the majority voting rule, k141_597014|CGC1 is predicted to have a xylan substrate.*

*Note: for many CGCs, only one of the two approaches produces substrate prediction. In this case, the two approaches produce different substrate assignments. We recommend a preference order: dbCAN-PUL blast search > dbCAN-sub majority voting. See our dbCAN3 paper*^2^ *for more details.*

**synteny.pdf**: a folder with syntenic block alignment plots between all CGCs and PULs.

**uniInput**: renamed Fasta file from input protein sequence file.

**Module 3: Read mapping (Fig. 3) to calculate abundance for CAZyme families, subfamilies, CGCs, and substrates**

**P8| Map reads of each sample to all CDS of co-assembled contigs (TIMING ~36 min)**

*$ bwa index prokka_Hadza_INDIV_157/Hadza_INDIV_157.ffn*

*$ mkdir samfiles*

*$ bwa mem -t 32 -o samfiles/Dry2014.CDS.sam prokka_Hadza_INDIV_157/Hadza_INDIV_157.ffn Dry2014_1_val_1.fq.gz Dry2014_2_val_2.fq.gz*

*$ bwa mem -t 32 -o samfiles/Wet2014.CDS.sam prokka_Hadza_INDIV_157/Hadza_INDIV_157.ffn Wet2014_1_val_1.fq.gz Wet2014_2_val_2.fq.gz*

Reads are mapped to the ffn files from Prokka.

**P9| Map reads of each sample to co-assembled contigs (TIMING ~36min)**

*$ bwa index megahit_Hadza_INDIV_157/Hadza_INDIV_157.contigs.fa*

*$ bwa mem -t 32 -o samfiles/Dry2014.sam megahit_Hadza_INDIV_157/Hadza_INDIV_157.contigs.fa Dry2014_1_val_1.fq.gz Dry2014_2_val_2.fq.gz*

*$ bwa mem -t 32 -o samfiles/Wet2014.sam megahit_Hadza_INDIV_157/Hadza_INDIV_157.contigs.fa Wet2014_1_val_1.fq.gz Wet2014_2_val_2.fq.gz*

Reads are mapped to the contig files from MEGAHIT.

**P10| Sort SAM files by coordinates (TIMING ~26min)**

*$ cd samfiles*

*$ samtools sort -@ 32 -o Wet2014.CDS.bam Wet2014.CDS.sam*

*$ samtools sort -@ 32 -o Dry2014.CDS.bam Dry2014.CDS.sam*

*$ samtools sort -@ 32 -o Wet2014.bam Wet2014.sam*

*$ samtools sort -@ 32 -o Dry2014.bam Dry2014.sam*

*$ rm -rf *sam*

*$ cd ..*

**P11| Calculate read count for all proteins in each sample using Bedtools (TIMING ~12min)**

*$ mkdir Wet2014_abund && cd Wet2014_abund*

*$ seqkit fx2tab -l -n -i ../prokka_Hadza_INDIV_157/Hadza_INDIV_157.ffn | awk '{print $1"\t"$2}' > Hadza_INDIV_157.length*

*$ seqkit fx2tab -l -n -i ../prokka_Hadza_INDIV_157/Hadza_INDIV_157.ffn | awk '{print $1"\t"0"\t"$2}' > Hadza_INDIV_157.bed*

*$ bedtools coverage -g Hadza_INDIV_157.length -sorted -a Hadza_INDIV_157.bed -counts -b Wet2014.CDS.bam > Wet2014.depth.txt*

*$ cd .. && mkdir Dry2014_abund && cd Dry2014_abund*

*$ seqkit fx2tab -l -n -i ../prokka_Hadza_INDIV_157/Hadza_INDIV_157.ffn | awk '{print $1"\t"$2}' > Hadza_INDIV_157.length*

*$ seqkit fx2tab -l -n -i ../prokka_Hadza_INDIV_157/Hadza_INDIV_157.ffn | awk '{print $1"\t"0"\t"$2}' > Hadza_INDIV_157.bed*

*$ bedtools coverage -g Hadza_INDIV_157.length -sorted -a Hadza_INDIV_157.bed -counts -b Dry2014.CDS.bam > Dry2014.depth.txt*

*$ cd ..*

Read counts are saved in *depth.txt* files of each sample.

**P12| Read count calculation for a given region of contigs using Samtools (TIMING ~2min)**

*$ cd Wet2014_abund*

*$ samtools index ../samfiles/Wet2014.bam*

*$ samtools depth -r k141_597014:160-8694 ../samfiles/Wet2014.bam > Wet2014.cgc.depth.txt*

**P13| dbcan_utils to calculate the abundance of CAZyme families, subfamilies, CGCs, and substrates (TIMING ~1min)**

*$ cd Wet2014_abund*

*$ dbcan_utils fam_abund -bt Wet2014.depth.txt -i ../Wet2014.dbCAN -a TPM*

*$ dbcan_utils fam_substrate_abund -bt Wet2014.depth.txt -i ../Wet2014.dbCAN -a TPM*

*$ dbcan_utils CGC_abund -bt Wet2014.depth.txt -i ../Wet2014.dbCAN -a TPM*

*$ dbcan_utils CGC_substrate_abund -bt Wet2014.depth.txt -i ../Wet2014.dbCAN -a TPM*

*$ cd .. && cd Dry2014_abund*

*$ dbcan_utils fam_abund -bt Dry2014.depth.txt -i ../Dry2014.dbCAN -a TPM*

*$ dbcan_utils fam_substrate_abund -bt Dry2014.depth.txt -i ../Dry2014.dbCAN -a TPM*

*$ dbcan_utils CGC_abund -bt Dry2014.depth.txt -i ../Dry2014.dbCAN -a TPM*

*$ dbcan_utils CGC_substrate_abund -bt Dry2014.depth.txt -i ../Dry2014.dbCAN -a TPM*

*cd ..*

We developed a set of Python scripts as dbcan_utils to take the raw read counts for all CDS as input and output the normalized abundances (**Box 7**) of CAZyme families, subfamilies, CGCs, and substrates (**Fig. 4**). The parameter *-a TPM* can also be two other metrics: RPM, or FPKM.

Four data folders will be needed as the input for dbcan_plot: (i) two abundance folders *Wet2014_abund* and *Dry2014_abund*, (ii) the CAZyme annotation folder *Hadza_INDIV_157.dbCAN,* and (iii) the *dbCAN-PUL* folder (under the *db* folder, released from *dbCAN-PUL.tar.gz*).

**P14| Heatmap for CAZyme substrate abundance across samples (Fig. S1A, TIMING 1min)**

*$ dbcan_plot heatmap_plot --samples Wet2014,Dry2014 -i Wet2014_abund/fam_substrate_abund.out,Dry2014_abund/fam_substrate_abund.out --show_abund --top 20*

Here we plot the top 20 substrates in the two samples. The input files are the two CAZyme substrate abundance files calculated based on dbCAN-sub result. The default heatmap is ranked by substrate abundances. To rank the heatmap according to abundance profile using the function clustermap of seaborn package, users can invoke the --cluster_map parameter.

**P15| Barplot for CAZyme family/subfamily/EC abundance across samples (Fig. S1B-D, TIMING 1min)**

*$ dbcan_plot bar_plot --samples Wet2014,Dry2014 --vertical_bar --top 20 -i Wet2014_abund/fam_abund.out,Dry2014_abund/fam_abund.out*

*dbcan_plot bar_plot --samples Wet2014,Dry2014 --vertical_bar --top 20 -i Wet2014_abund/subfam_abund.out,Dry2014_abund/subfam_abund.out*

Users can choose to generate a barplot instead of heatmap using the bar_plot method.

**P16| Synteny plot between a CGC and its best PUL hit with read mapping coverage to CGC (Fig. S1E, TIMING 1min)**

*$ dbcan_plot CGC_synteny_coverage_plot -i Hadza_INDIV_157.dbCAN --cgcid 'k141_597014|CGC1' --readscount Wet2014_abund/Wet2014.cgc.depth.txt*

*$ dbcan_plot CGC_synteny_coverage_plot -i Hadza_INDIV_157.dbCAN --cgcid 'k141_597014|CGC1' --readscount Dry2014_abund/Dry2014.cgc.depth.txt*

The Hadza_INDIV_157.dbCAN folder contains the PUL_blast.out file. Using this file, the cgc_standard.out file, and the best PUL’s gff file in dbCAN-PUL.tar.gz, the CGC_synteny_plot method will create the CGC-PUL synteny plot. The –cgcid parameter is required to specify which CGC to plot (‘k141_597014|CGC1' in this example). The *Wet2014.cgc.depth.txt* file is used to plot the read mapping coverage. Different from the individual sample assembly route, the CGCs are predicted for the co-assembled contigs, so occurrence and abundance of CGCs can be directly compared between the two samples. For example, the above commands will generate the CGC-PUL synteny plots in both Dry2014 and Wet2014 samples for the same ‘k141_597014|CGC1'.

If users only want to plot the CGC structure:

*$ dbcan_plot CGC_plot -i Hadza_INDIV_157.dbCAN --cgcid 'k141_597014|CGC1’*

If users only want to plot the CGC structure plus the read mapping coverage:

*$ dbcan_plot CGC_coverage_plot -i Hadza_INDIV_157.dbCAN --cgcid 'k141_597014|CGC1' --readscount Wet2014_abund/Wet2014.cgc.depth.txt*

If users only want to plot the synteny between the CGC and PUL:

*$ dbcan_plot CGC_synteny_plot -i Hadza_INDIV_157.dbCAN --cgcid 'k141_597014|CGC1'*

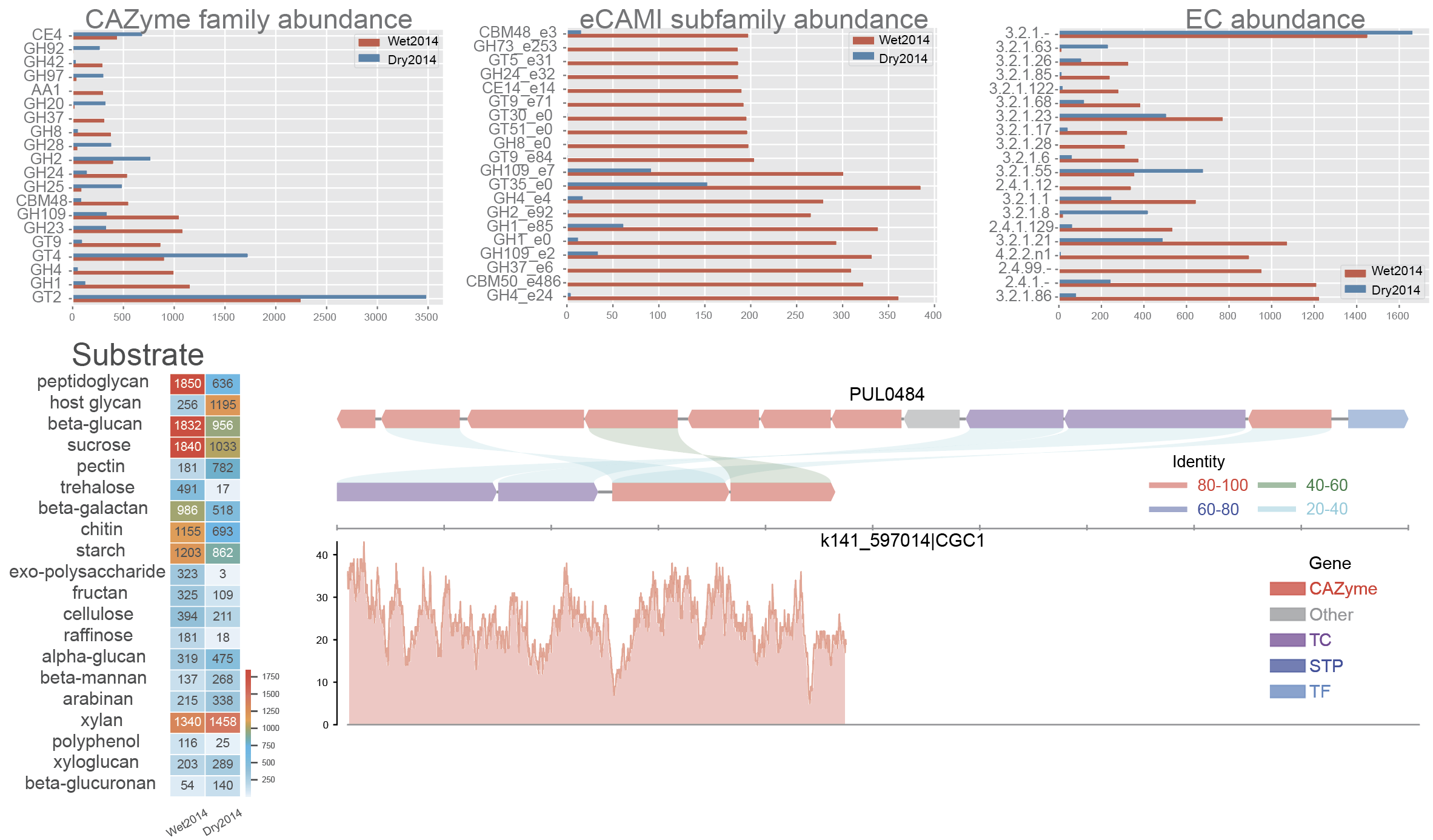

***Fig. S1.*** ***Data visualization of abundances of CAZymes, CGCs, and substrates****. (****A****) Heatmap of CAZyme substrate abundance in the two samples (Carter2023 dataset: see Table 2) calculated based on dbCAN-sub search results and substrate mapping. Abundance values are TPM. (****B****) Barplot of 20 CAZyme family abundance in the two samples calculated based on dbCAN search results. (****C****) Barplot of 20 CAZyme eCAMI subfamily abundance in the two samples calculated based on dbCAN-sub search results. (****D****) Barplot of 20 CAZyme EC abundance in the two samples calculated based on dbCAN-sub search results. (****E****) Synteny plot between an example CGC (CGC1 of contig k141_63712 from the Wet2014 sample) and its best PUL hit (PUL0373 from dbCAN-PUL, with experimentally verified substrate sucrose and raffinose) with read mapping coverage plot (y-axis is the read depth) shown in the bottom.*

**Procedure of the assembly-free route (Carter2023)**

This is the procedure for the assembly-free route of dataset1–Carter2023.

The procedure has 2 modules (**Fig. 3**) and 5 steps (P1-P5). Unlike the individual sample assembly and co-assembly routes, the assembly-free route does not assemble the reads to contigs. Since no assembled contigs are involved, CGCs and CGC-based substrate prediction steps are not applicable for this route.

Compared to the two assembly-based routes, the assembly-free route has advantages in two aspects: (i) higher speed, and (ii) ability to detect CAZymes with low abundance. However, a recent metatranscriptomic workflow development work showed that the assembly-free approach had much higher false positive rate and lower precision than the assembly-based approach^9^. Therefore, it is expected that the assemble-free route produces less accurate result than the other two routes and may over-estimate the CAZyme abundance and occurrence (**Table S1**).

All the steps will be conducted in a separate folder named assembly-free.

<https://bcb.unl.edu/dbCAN_tutorial/dataset1-Carter2023/assembly_free/>

**Module 1: Reads processing (Fig. 3) to prepare Fasta files (fna) of reads**

**P1| Check contamination (TIMING ~10min)**

*$ kraken2 --threads 32 --quick --paired --db K2 --report Wet2014.kreport --output Wet2014. kraken.output Wet2014_1.fastq.gz Wet2014_2.fastq.gz*

*$ kraken2 --threads 32 --quick --paired --db K2 --report Dry2014.kreport --output Dry2014. kraken.output Dry2014_1.fastq.gz Dry2014_2.fastq.gz*

Kraken2 found very little contamination in the Carter2023 data. Consequently, there was no need for the contamination removal step.

If contamination is identified, users can align the reads to the reference genomes of potential contamination source organisms to remove the aligned reads (**Box 1**). The most common source in human microbiome studies is from human hosts.

**Box 1| Example to remove contamination reads from human**

Kraken2 will produce the following output files.

*-rw-rw-r-- 1 jinfang jinfang 2.0G Dec 12 10:24 Dry2014.kraken.output*

*-rw-rw-r-- 1 jinfang jinfang 1.2M Dec 12 10:25 Dry2014.kreport*

*-rw-rw-r-- 1 jinfang jinfang 5.1G Dec 12 09:47 Wet2014.kraken.output*

*-rw-rw-r-- 1 jinfang jinfang 1.1M Dec 12 09:48 Wet2014.kreport*

Suppose from these files, we have identified humans as the contamination source, we can use the following commands to remove the contamination reads by aligning reads to the human reference genome.

*$ wget https://ftp.ensembl.org/pub/release-110/fasta/homo_sapiens/dna/Homo_sapiens.GRCh38.dna.primary_assembly.fa.gz*

*$ bwa index -p hg38 Homo_sapiens.GRCh38.dna.primary_assembly.fa.gz*

*$ bwa mem hg38 Wet2014_1.fastq.gz Wet2014_2.fastq.gz -t 32 -o Wet2014.hg38.sam*

*$ bwa mem hg38 Dry2014_1.fastq.gz Dry2014_2.fastq.gz -t 32 -o Dry2014.hg38.sam*

*$ samtools view -f 12 Wet2014.hg38.sam > Wet2014.hg38.unmap.bam*

*$ samtools view -f 12 Dry2014.hg38.sam > Dry2014.hg38.unmap.bam*

*$ samtools fastq -1 Wet2014_1.clean.fq.gz -2 Wet2014_2.clean.fq.gz Wet2014.hg38.unmap.bam*

*$ samtools fastq -1 Dry2014_1.clean.fq.gz -2 Dry2014_2.clean.fq.gz Dry2014.hg38.unmap.bam*

**P2| Trim adapter and low-quality reads (TIMING ~20min)**

*$ trim_galore --paired Wet2014_1.fastq.gz Wet2014_2.fastq.gz --illumina -j 36*

*$ trim_galore --paired Dry2014_1.fastq.gz Dry2014_2.fastq.gz --illumina -j 36*

We specified --illumina to indicate that the reads were generated using the Illumina sequencing platform. Nonetheless, trim_galore possesses the ability to automatically detect the adapter, providing flexibility in adapter handling for users who may know the specific sequencing platform. Details of trimming are available in the trimming report file (**Box 2**).

**Box 2| Example output of trim_galore**

In addition to the trimmed read files, Trim_galore also generates a trimming report file. The trimming report contains details on read trimming, such as the number of trimmed reads.

*-rw-rw-r-- 1 jinfang jinfang 4.2K Dec 13 01:48 Dry2014_1.fastq.gz_trimming_report.txt*

*-rw-rw-r-- 1 jinfang jinfang 2.0G Dec 13 01:55 Dry2014_1_val_1.fq.gz*

*-rw-rw-r-- 1 jinfang jinfang 4.4K Dec 13 01:55 Dry2014_2.fastq.gz_trimming_report.txt*

*-rw-rw-r-- 1 jinfang jinfang 2.4G Dec 13 01:55 Dry2014_2_val_2.fq.gz*

*-rw-rw-r-- 1 jinfang jinfang 4.4K Dec 13 01:30 Wet2014_1.fastq.gz_trimming_report.txt*

*-rw-rw-r-- 1 jinfang jinfang 3.4G Dec 13 01:46 Wet2014_1_val_1.fq.gz*

*-rw-rw-r-- 1 jinfang jinfang 4.6K Dec 13 01:46 Wet2014_2.fastq.gz_trimming_report.txt*

*-rw-rw-r-- 1 jinfang jinfang 3.7G Dec 13 01:46 Wet2014_2_val_2.fq.gz*

**! CAUTION**

During the trimming process, certain reads may be entirely removed due to low quality in its entirety. Using the --retain_unpaired parameter in trim_galore allows for the preservation of single-end reads. In this protocol, this option was not select, so that both reads of a forward-revise pair were removed.

**P3| Covert the fastq file to fasta (~9min)**

*$ seqtk seq -a Dry2014_1_val_1.fq.gz > Dry2014_1.fna*

*$ seqtk seq -a Dry2014_2_val_2.fq.gz > Dry2014_2.fna*

*$ seqtk seq -a Wet2014_1_val_1.fq.gz > Wet2014_1.fna*

*$ seqtk seq -a Wet2014_2_val_2.fq.gz > Wet2014_2.fna*

**Module 2: Read mapping (Fig. 3) to calculate abundance for CAZyme families, subfamilies, and substrates**

**P4|** **Map reads to proteins in CAZyDB by using DIAMOND blastx (~5h10min)**

*$ ln -s db/CAZy.dmnd .*

*$ diamond blastx --db CAZy --out Dry2014_1.blastx --threads 32 --outfmt 6 qseqid sseqid pident length mismatch gapopen qstart qend sstart send evalue bitscore qcovhsp qlen slen --id 80 --query-cover 90 --query Dry2014_1.fna --max-target-seqs 1 --quiet*

*$ diamond blastx --db CAZy --out Dry2014_2.blastx --threads 32 --outfmt 6 qseqid sseqid pident length mismatch gapopen qstart qend sstart send evalue bitscore qcovhsp qlen slen --id 80 --query-cover 90 --query Dry2014_2.fna --max-target-seqs 1 --quiet*

*$ diamond blastx --db CAZy --out Wet2014_1.blastx --threads 32 --outfmt 6 qseqid sseqid pident length mismatch gapopen qstart qend sstart send evalue bitscore qcovhsp qlen slen --id 80 --query-cover 90 --query Wet2014_1.fna --max-target-seqs 1 --quiet*

*$ diamond blastx --db CAZy --out Wet2014_2.blastx --threads 32 --outfmt 6 qseqid sseqid pident length mismatch gapopen qstart qend sstart send evalue bitscore qcovhsp qlen slen --id 80 --query-cover 90 --query Wet2014_2.fna --max-target-seqs 1 --quiet*

The parameters --query-cover 90 and --id 80 are required to align the reads to CAZy database. The thresholds of these parameters will significantly impact the results. The --max-target-seqs 1 parameter specify to only keep the best CAZy hit, which means each read will be mapped to only one CAZyme protein in the CAZyDB.

**P5| dbcan_asmfree to** **calculate the abundance of CAZyme families, subfamily, EC and substrate (~11min)**

We developed a Python script as dbcan_asmfree (included in the run_dbcan package) to take the DIAMOND blastx results from pair-end reads and the raw reads as inputs. With the *diamond_fam_abund* method, it will output the CAZyme family abundance, with the *diamond_subfam_abund* method, it will output the eCAMI subfamily abundance, with the *diamond_EC_abund* method, it will output the EC abundance and with the *diamond_substrate_abund* method, it will output the substrate abundance.

*$ dbcan_asmfree diamond_fam_abund -paf1 Dry2014_1.blastx -paf2 Dry2014_2.blastx --raw_reads Dry2014_1_val_1.fq.gz -n FPKM -o Dry2014_fam_abund*

*$ dbcan_asmfree diamond_fam_abund -paf1 Wet2014_1.blastx -paf2 Wet2014_2.blastx --raw_reads Wet2014_1_val_1.fq.gz -n FPKM -o Wet2014_fam_abund*

*$ dbcan_asmfree* *diamond_subfam_abund -paf1 Dry2014_1.blastx -paf2 Dry2014_2.blastx --raw_reads Dry2014_1_val_1.fq.gz -o Dry2014_subfam_abund -n FPKM*

*$ dbcan_asmfree diamond_subfam_abund -paf1 Wet2014_1.blastx -paf2 Wet2014_2.blastx --raw_reads Wet2014_1_val_1.fq.gz -o Wet2014_subfam_abund -n FPKM*

*$ dbcan_asmfree diamond_EC_abund -i Dry2014_subfam_abund -o Dry2014_EC_abund*

*$ dbcan_asmfree diamond_EC_abund -i Wet2014_subfam_abund -o Wet2014_EC_abund*

*$ dbcan_asmfree diamond_substrate_abund -i Dry2014_subfam_abund -o Dry2014_substrate_abund*

*$ dbcan_asmfree* *diamond_substrate_abund -i Wet2014_subfam_abund -o Wet2014_substrate_abund*

The substrate prediction in the assemble-free route is based on the fam-substrate-mapping-08012023.tsv mapping table. Specifically, from the blastx result files, reads are first grouped according to their best CAZyDB proteins. Then, reads are further grouped according to the CAZyme families and eCAMI subfamilies where the best CAZyDB proteins belong to. Lastly, the fam-substrate-mapping-08012023.tsv mapping table is used to map CAZyme families and ECs (from eCAMI subfamilies) to substrates. Therefore, the substrate abundance is calculated in a similar way as implemented in dbCAN-sub.

**Module 4: dbcan_plot for data visualization (Fig. 3) of abundances of CAZymes and substrates (TIMING variable)**

**P6| Barplot for CAZyme family, subfamily, and EC abundance across samples (Fig. S2A-C) (TIMING 1min)**

*$ dbcan_plot bar_plot --samples Wet2014,Dry2014 --vertical_bar --top 20 -i Dry2014.blastx.CAZy.FPKM.tsv,Wet2014.blastx.CAZy.FPKM.tsv*

**P7| Heatmap for CAZyme substrate abundance across samples (Fig. S2D) (TIMING 1min)**

*$ dbcan_plot heatmap_plot --samples Wet2014,Dry2014 --show_abund --top 20 -i Dry2014.blastx.substrate.tsv,Wet2014.blastx.substrate.tsv*

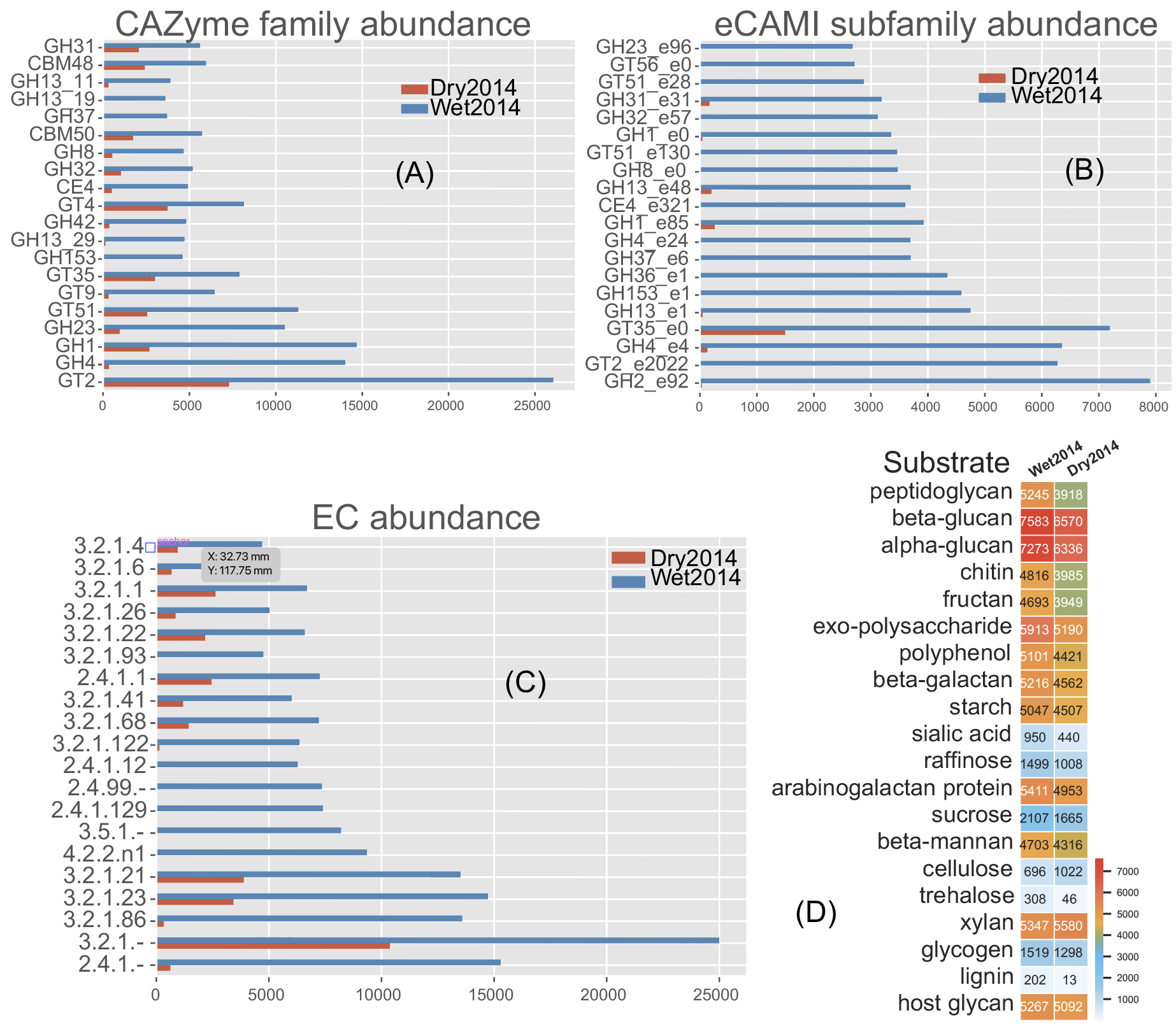

***Fig. S2.*** ***Data visualization of abundances of CAZymes, substrate, EC, and substrates from the assembly-free route****. (****A****) Barplot of 20 CAZyme family abundance in the two (Carter2023 dataset: see Table 2) samples calculated based on Diamond blastx read mapping to CAZyDB proteins. (****C****) Barplot of 20 CAZyme EC abundance in the two samples calculated based on Diamond blastx read mapping to CAZyDB proteins and then to eCAMI subfamilies. (****D****) Heatmap of CAZyme substrate abundance in the two samples calculated based on Diamond blastx read mapping to CAZyDB proteins and then to eCAMI subfamilies and to substrates. Abundance values are FPKM.*

**Procedure of the sub-sampling route (Carter2023)**

This is the procedure for the sub-sampling route of Carter2023.

As mentioned in the main text, in order to verify the findings made in the independent sample assembly route, we developed a sub-sampling procedure to rule out the unequal sequencing depth in different samples. The sub-sampling can also reduce the CPU time and RAM use in running all the analyses, especially when combining multiple samples for a co-assembly of metagenomic reads, e.g., in a recent co-assembly study of 124 marine metagenomic samples^10^.

Specifically, after an equal number of reads (20,000,000) were randomly extracted from Wet2014 and Dry2014 (**Table 2**), we repeated the METAHIT assembly and Prokka gene prediction and all the analyses using the same procedure in the independent sample assembly route.

The procedure has 4 modules (**Fig. 3**) and 16 steps (P1-P16). All the steps will be conducted in a separate folder named sub_samples.

<https://bcb.unl.edu/dbCAN_tutorial/dataset1-Carter2023/sub_samples/>

**Module 1: Reads processing (Fig. 3) to obtain contigs**

**P1| Check contamination (TIMING ~10min)**

*$ kraken2 --threads 32 --quick --paired --db K2 --report Wet2014.kreport --output Wet2014. kraken.output Wet2014_1.fastq.gz Wet2014_2.fastq.gz*

*$ kraken2 --threads 32 --quick --paired --db K2 --report Dry2014.kreport --output Dry2014. kraken.output Dry2014_1.fastq.gz Dry2014_2.fastq.gz*

Kraken2 found very little contamination in the Carter2023 data. Consequently, there was no need for the contamination removal step.

If contamination is identified, users can align the reads to the reference genomes of potential contamination source organisms to remove the aligned reads (**Box 1**). The most common source in human microbiome studies is from human hosts.

**P2| Sub-sample raw reads, trim adapter and low-quality reads (TIMING ~36min)**

*$ reformat.sh in=Dry2014_1_val_1.fq.gz in2=Dry2014_2_val_2.fq.gz out=subsample_Dry2014_1_val_1.fq.gz out2=subsample_Dry2014_2_val_2.fq.gz sample=20000000*

*$ reformat.sh in=Wet2014_1_val_1.fq.gz in2=Wet2014_2_val_2.fq.gz out=subsample_Wet2014_1_val_1.fq.gz out2=subsample_Wet2014_2_val_2.fq.gz sample=20000000*

*$ mv subsample_Dry2014_1_val_1.fq.gz Dry2014_1_val_1.fq.gz*

*$ mv subsample_Dry2014_2_val_2.fq.gz Dry2014_2_val_2.fq.gz*

*$ mv subsample_Wet2014_2_val_2.fq.gz Wet2014_2_val_2.fq.gz*

*$ mv subsample_Wet2014_1_val_1.fq.gz Wet2014_1_val_1.fq.gz*

*$ trim_galore --paired Wet2014_1.fastq.gz Wet2014_2.fastq.gz --illumina -j 36*

*$ trim_galore --paired Dry2014_1.fastq.gz Dry2014_2.fastq.gz --illumina -j 36*

The *reformat.sh* script in the BBTools package is used for sub-sampling 20 million reads, followed by trim_galore quality trimming (**Box 2**).

**! CAUTION**

During the trimming process, certain reads may be entirely removed due to low quality in its entirety. Using the --retain_unpaired parameter in trim_galore allows for the preservation of single-end reads. In this protocol, this option was not select, so that both reads of a forward-revise pair were removed.

The rest of the procedure (P3-P16) is the same as what is presented in the main text for the individual sample assembly route.

**P3| Assemble reads into contigs (TIMING ~50min)**

*$ megahit -m 0.5 -t 32 -o megahit_Wet2014 -1 Wet2014_1_val_1.fq.gz -2 Wet2014_2_val_2.fq.gz --out-prefix Wet2014 --min-contig-len 1000*

*$ megahit -m 0.5 -t 32 -o megahit_Dry2014 -1 Dry2014_1_val_1.fq.gz -2 Dry2014_2_val_2.fq.gz --out-prefix Dry2014 --min-contig-len 1000*

MEGAHIT generates two output folders *megahit_Wet2014 and megahit_Dry2014*. Each contains five files and one sub-folder (**Box 3**). Wet2014.contigs.fa is the final contig sequence file. We set --min-contig-len 1000, a common practice to retain all contigs longer than 1,000 base pairs.

**P4| Predict genes by Prokka (TIMING ~40min)**

*$ prokka --kingdom Bacteria --cpus 36 --outdir prokka_Wet2014 --prefix Wet2014 --addgenes --addmrna --locustag Wet2014 megahit_Wet2014/Wet2014.contigs.fa*

*$ prokka --kingdom Bacteria --cpus 36 --outdir prokka_Dry2014 --prefix Dry2014 --addgenes --addmrna --locustag Dry2014 megahit_Dry2014/Dry2014.contigs.fa*

The parameter --kingdom Bacteria is required for bacterial gene prediction. To optimize performance, --CPU 36 instructs the utilization of 36 computer processors. The output files comprise of both protein and CDS sequences in Fasta format (e.g., *Wet2014.faa* and *Wet2014.ffn* in **Box 4**).

**Module 2: run_dbcan annotation (Fig. 3) to obtain CAZymes, CGCs, and substrates**

**P5| CAZyme annotation at family level (TIMING ~10min)**

*$ run_dbcan prokka_Wet2014/Wet2014.faa protein --hmm_cpu 32 --out_dir Wet2014.CAZyme --tools hmmer --db_dir db*

*$ run_dbcan prokka_Dry2014/Dry2014.faa protein --dbcan_thread 32 --stp_cpu 32 -c prokka_Dry2014/Dry2014.gff --cgc_substrate --out_dir Dry2014.dbCAN --dia_cpu 32 --hmm_cpu 32 --tf_cpu 32*

Files in in an example output folder is explained in **Box 6**.

*$ cd ..*

Read counts are saved in *depth.txt* files of each sample.

**P12| Read count calculation for a given region of contigs using Samtools (TIMING ~1min)**

*$ cd Wet2014_abund*

*$ samtools index ../samfiles/Wet2014.bam*

*$ samtools depth -r k141_63712:36760-50331 ../samfiles/Wet2014.bam > Wet2014.cgc.depth.txt*

*$ cd ..*

The parameter *-r* *k141_63712:36760-50331* specifies a region in a contig. For any CGC, its positional range can be found in the file cgc_standard.out produced by run_dbcan (**Box 6**). The *depth.txt* files contain the raw read counts for the specified region.

**P16| Synteny plot between a CGC and its best PUL hit with read mapping coverage to CGC (Fig. S3E) (TIMING 1min)**

*$ dbcan_plot CGC_synteny_coverage_plot -i Wet2014.dbCAN --cgcid ‘**k141_63712|CGC1' --readscount Wet2014_abund/Wet2014.cgc.depth.txt*

The Wet2014.dbCAN folder contains the PUL_blast.out file. Using this file, the cgc_standard.out file, and the best PUL’s gff file in dbCAN-PUL.tar.gz, the CGC_synteny_plot method will create the CGC-PUL synteny plot. The –cgcid parameter is required to specify which CGC to be plotted (‘k141_63712|CGC1' in this example). The *Wet2014.cgc.depth.txt* file is used to plot the read mapping coverage.

If users only want to plot the CGC structure:

*$ dbcan_plot CGC_plot -i Wet2014.dbCAN --cgcid ‘k141_63712|CGC1'*

If users only want to plot the CGC structure plus the read mapping coverage:

*$ dbcan_plot CGC_coverage_plot -i Wet2014.dbCAN --cgcid ‘k141_63712|CGC1’--readscount Wet2014_abund/Wet2014.cgc.depth.txt*

If users only want to plot the synteny between the CGC and PUL:

*$ dbcan_plot CGC_synteny_plot -i Wet2014.dbCAN --cgcid* *‘k141_63712|CGC1’*

**! CAUTION**

The CGC IDs in different samples do not match each other. For example, specifying -i Wet2014.dbCAN is to plot the 'k141_63712|CGC1’ in the Wet2014 sample. The ‘k141_63712|CGC1’ in the Dry2014 sample will be different.

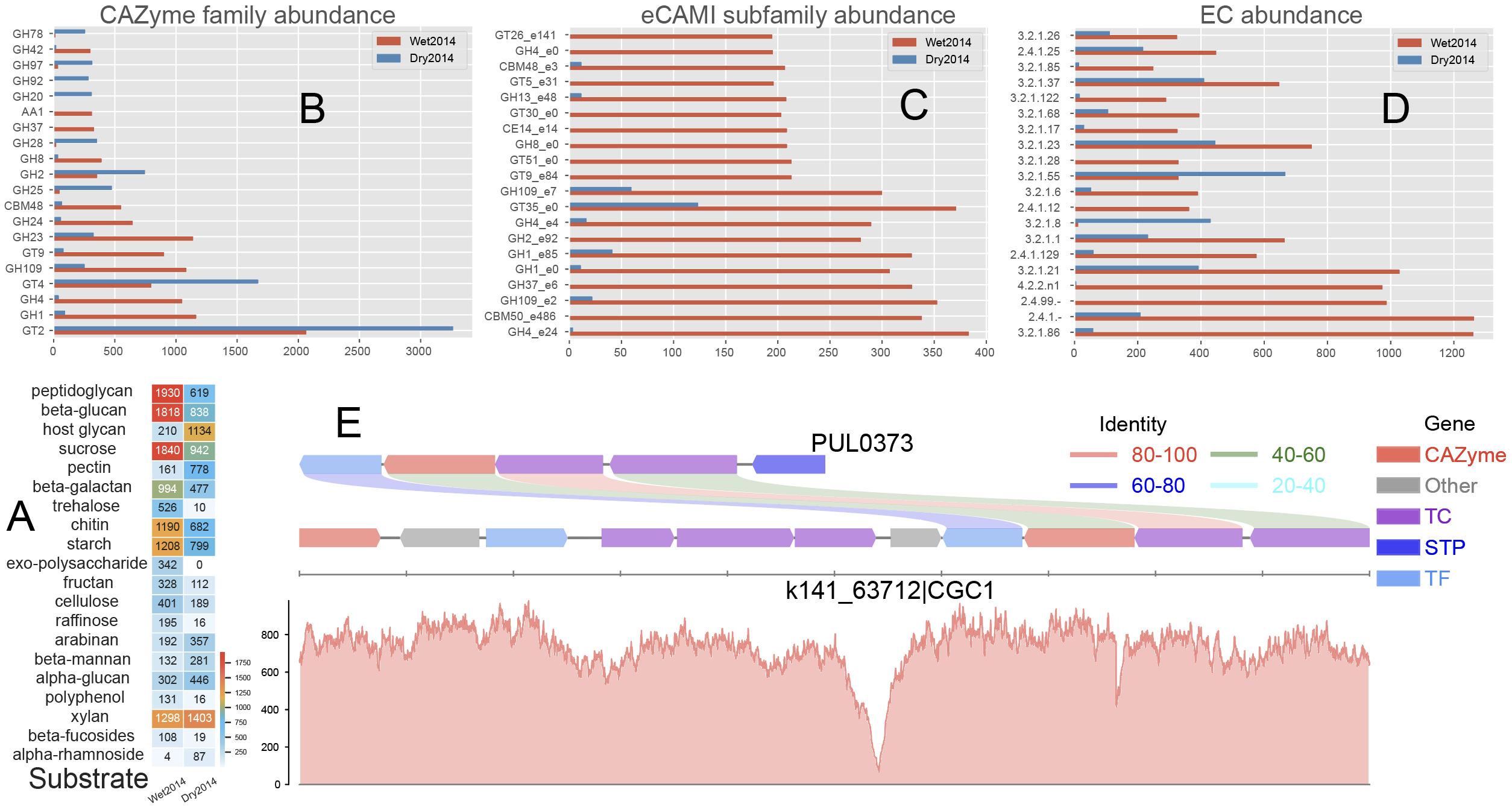

***Fig. S3.*** ***Data visualization of abundances of CAZymes, CGCs, and substrates****. (****A****) Heatmap of CAZyme substrate abundance in the two samples (Carter2023 dataset: see Table 2) calculated based on dbCAN-sub search results and substrate mapping. Abundance values are TPM. (****B****) Barplot of 20 CAZyme family abundance in the two samples calculated based on dbCAN search results. (****C****) Barplot of 20 CAZyme eCAMI subfamily abundance in the two samples calculated based on dbCAN-sub search results. (****D****) Barplot of 20 CAZyme EC abundance in the two samples calculated based on dbCAN-sub search results. (****E****) Synteny plot between an example CGC (CGC1 of contig k141_63712 from the Wet2014 sample) and its best PUL hit (PUL0373 from dbCAN-PUL, with experimentally verified substrate sucrose and raffinose) with read mapping coverage plot (y-axis is the read depth) shown in the bottom.*

**Procedure of individual sample assembly for Wastyk2021**

The Wastyk2021 dataset was published in 2021^12^ from a human dietary intervention study. In the published paper, researchers studied how high-fermented and high-fiber diets influence the human microbiome metabolism and modulate the human immune status. Among various data analyses conducted in the paper^12^, CAZymes were mined from shotgun metagenomic reads of 18 healthy human participants, and each participant had four time points of stool samples for metagenome sequencing. CAZyme abundance profiles were compared before and after the high-fiber intervention (baseline vs high-fiber). One of the main findings from their CAZyme analysis was that high-fiber consumption increased the CAZyme abundance. For this protocol, we will select two samples (paired-end 2x146bp reads) of two time points (day 2 before high-fiber diet as baseline, and 10 weeks after high-fiber diet as intervention) from one participant (**Table S2**). The protocol is for the individual sample route (**Fig. 3**).

**Table S2. Summary of the example microbiome dataset**

| **Dataset** (Wastyk HC et al., Cell 2021 ^12^) | | |
| --- | --- | --- |
| Sample name | fefifo_8022_1 | fefifo_8022_7 |
| SRA ID | SRR15032216 | SRR15032213 |
| Treatment | baseline | high fiber diet |
| Clean read count | 9,480,450 | 17,336,994 |

The raw read data, intermediate data from each analysis step, and final result data and visualization files are organized in nested folders available on our website <https://bcb.unl.edu/dbCAN_tutorial/dataset2-Wastyk2021/>, **Fig. 5**) and <https://dbcan.readthedocs.io>.

The procedure has 4 modules (**Fig. 3**) and 16 steps (P1-P16).

**Module 1: Reads processing (Fig. 3) to obtain contigs**

**P1| Check contamination (TIMING ~10min)**

*$ wget https://bcb.unl.edu/dbCAN_tutorial/dataset2-Wastyk2021/fefifo_8022_1__shotgun_1.fastq.gz*

*$ wget https://bcb.unl.edu/dbCAN_tutorial/dataset2-Wastyk2021/fefifo_8022_1__shotgun_2.fastq.gz*

*$ wget https://bcb.unl.edu/dbCAN_tutorial/dataset2-Wastyk2021/fefifo_8022_7__shotgun_1.fastq.gz*

*$ wget https://bcb.unl.edu/dbCAN_tutorial/dataset2-Wastyk2021/fefifo_8022_7__shotgun_2.fastq.gz*

*$ kraken2 --threads 32 --quick --paired --db K2 --report fefifo_8022_1.kreport --output fefifo_8022_1.kraken.output fefifo_8022_1__shotgun_1.fastq.gz fefifo_8022_1__shotgun_2.fastq.gz*

*$ kraken2 --threads 32 --quick --paired --db K2 --report fefifo_8022_7.kreport --output fefifo_8022_7.kraken.output fefifo_8022_7__shotgun_1.fastq.gz fefifo_8022_7__shotgun_2.fastq.gz*

Kraken2 found very little contamination in the data. Consequently, there was no need for the contamination removal step.

If contamination is identified, users can align the reads to the reference genomes of potential contamination source organisms to remove the aligned reads (**Box 1**). The most common source in human microbiome studies is from human hosts.

**Box 1| Example to remove contamination reads from human**

Kraken2 will produce the following output files.

*-rw-rw-r-- 1 jinfang jinfang 1.1G Sep 21 23:17 fefifo_8022_1.kraken.output*

*-rw-rw-r-- 1 jinfang jinfang 991K Sep 21 23:19 fefifo_8022_1.kreport*

*-rw-rw-r-- 1 jinfang jinfang 574M Sep 21 23:21 fefifo_8022_7.kraken.output*

*-rw-rw-r-- 1 jinfang jinfang 949K Sep 21 23:22 fefifo_8022_7.kreport*

Suppose from these files, we have identified humans as the contamination source, we can use the following commands to remove the contamination reads by aligning reads to the human reference genome.

*$ wget https://ftp.ensembl.org/pub/release-110/fasta/homo_sapiens/dna/Homo_sapiens.GRCh38.dna.primary_assembly.fa.gz*

*$ bwa index -p hg38 Homo_sapiens.GRCh38.dna.primary_assembly.fa.gz*

*$ bwa mem hg38 fefifo_8022_1__shotgun_1.fastq.gz fefifo_8022_1__shotgun_2.fastq.gz -t 32 -o fefifo_8022_1.hg38.sam*

*$ bwa mem hg38 fefifo_8022_7__shotgun_1.fastq.gz fefifo_8022_7__shotgun_2.fastq.gz -t 32 -o fefifo_8022_7.hg38.sam*

*$ samtools view -f 12 fefifo_8022_1.hg38.sam > fefifo_8022_1.hg38.unmap.bam*

*$ samtools view -f 12 fefifo_8022_7.hg38.sam > fefifo_8022_7.hg38.unmap.bam*

*$ samtools fastq -1 fefifo_8022_1_1.clean.fq.gz -2 fefifo_8022_1_2.clean.fq.gz fefifo_8022_1.hg38.unmap.bam*

*$ samtools fastq -1 fefifo_8022_7_1.clean.fq.gz -2 fefifo_8022_7_2.clean.fq.gz fefifo_8022_7.hg38.unmap.bam*

**P2| Trim adapter and low-quality reads (TIMING ~20min)**

*$ trim_galore --paired fefifo_8022_1__shotgun_1.fastq.gz fefifo_8022_1__shotgun_2.fastq.gz --illumina -j 36*

*$ trim_galore --paired fefifo_8022_7__shotgun_1.fastq.gz fefifo_8022_7__shotgun_2.fastq.gz --illumina -j 36*

We specified --illumina to indicate that the reads were generated using the Illumina sequencing platform. Nonetheless, trim_galore can automatically detect adapters, providing flexibility for users who may know the specific sequencing platform. Details of trimming are available in the trimming report file (**Box 2**).

**Box 2| Example output of trim_galore**

In addition to the trimmed read files, Trim_galore also generates a trimming report file. The trimming report contains details on read trimming, such as the number of trimmed reads.

```
-rw-rw-r-- 1 jinfang jinfang  429M Oct 30 22:44 fefifo_8022_1__shotgun_1.fastq.gz
-rw-rw-r-- 1 jinfang jinfang  4.1K Oct 31 05:15 fefifo_8022_1__shotgun_1.fastq.gz_trimming_report.txt
-rw-rw-r-- 1 jinfang jinfang  390M Oct 31 05:16 fefifo_8022_1__shotgun_1_val_1.fq.gz
-rw-rw-r-- 1 jinfang jinfang  540M Oct 30 22:44 fefifo_8022_1__shotgun_2.fastq.gz
-rw-rw-r-- 1 jinfang jinfang  4.2K Oct 31 05:16 fefifo_8022_1__shotgun_2.fastq.gz_trimming_report.txt
-rw-rw-r-- 1 jinfang jinfang  499M Oct 31 05:16 fefifo_8022_1__shotgun_2_val_2.fq.gz
-rw-rw-r-- 1 jinfang jinfang  931M Oct 30 22:34 fefifo_8022_7__shotgun_1.fastq.gz
-rw-rw-r-- 1 jinfang jinfang  4.2K Oct 31 05:17 fefifo_8022_7__shotgun_1.fastq.gz_trimming_report.txt
-rw-rw-r-- 1 jinfang jinfang  861M Oct 31 05:20 fefifo_8022_7__shotgun_1_val_1.fq.gz
-rw-rw-r-- 1 jinfang jinfang  1.1G Oct 30 22:34 fefifo_8022_7__shotgun_2.fastq.gz
-rw-rw-r-- 1 jinfang jinfang  4.4K Oct 31 05:20 fefifo_8022_7__shotgun_2.fastq.gz_trimming_report.txt
-rw-rw-r-- 1 jinfang jinfang 1003M Oct 31 05:20 fefifo_8022_7__shotgun_2_val_2.fq.gz
```

**! CAUTION**

During the trimming process, certain reads may be entirely removed due to low quality in its entirety. Using the --retain_unpaired parameter in trim_galore allows for the preservation of single-end reads. In this protocol, this option was not selected, so that both reads of a forward-revise pair were removed.

**P3| Assemble reads into contigs (TIMING ~84min)**

*$ megahit -m 0.5 -t 32 -o megahit_fefifo_8022_1 -1 fefifo_8022_1__shotgun_1_val_1.fq.gz -2 fefifo_8022_1__shotgun_2_val_2.fq.gz --out-prefix fefifo_8022_1 --min-contig-len 1000*

*$ megahit -m 0.5 -t 32 -o megahit_fefifo_8022_7 -1 fefifo_8022_7__shotgun_1_val_1.fq.gz -2 fefifo_8022_7__shotgun_2_val_2.fq.gz --out-prefix fefifo_8022_7 --min-contig-len 1000*

MEGAHIT generates two output folders megahit_fefifo_8022_1 and megahit_fefifo_8022_7. Each contains five files and one sub-folder (**Box 3**). fefifo_8022_1.contigs.fa is the final contig sequence file. We set --min-contig-len 1000, a common practice to retain all contigs longer than 1,000 base pairs.

**Box 3| Example output of MEGAHIT**

```
-rw-rw-r--  1 jinfang jinfang  262 Oct 31 05:49 checkpoints.txt
-rw-rw-r--  1 jinfang jinfang    0 Oct 31 05:49 done
-rw-rw-r--  1 jinfang jinfang  97M Oct 31 05:49 fefifo_8022_1.contigs.fa
-rw-rw-r--  1 jinfang jinfang 149K Oct 31 05:49 fefifo_8022_1.log
drwxrwxr-x  2 jinfang jinfang 4.0K Oct 31 05:49 intermediate_contigs
-rw-rw-r--  1 jinfang jinfang 1.1K Oct 31 05:27 options.json
```

**P4| Predict genes by Prokka (TIMING ~40min)**

*$ prokka --kingdom Bacteria --cpus 36 --outdir prokka_fefifo_8022_1 --prefix fefifo_8022_1 --addgenes --addmrna --locustag fefifo_8022_1 megahit_fefifo_8022_1/fefifo_8022_1.contigs.fa*

*$ prokka --kingdom Bacteria --cpus 36 --outdir prokka_fefifo_8022_7 --prefix fefifo_8022_7 --addgenes --addmrna --locustag fefifo_8022_7 megahit_fefifo_8022_7/fefifo_8022_7.contigs.fa*

The parameter --kingdom Bacteria is required for bacterial gene prediction. To optimize performance, --CPU 36 instructs the utilization of 36 computer processors. The output files comprise of both protein and CDS sequences in Fasta format (e.g., fefifo_8022_1.faa and fefifo_8022_1.ffn in **Box 4**).

**Box 4| Example output of Prokka**

```
-rw-rw-r--  1 jinfang jinfang  181 Oct 31 22:28 errorsummary.val
-rw-rw-r--  1 jinfang jinfang 2.6M Oct 31 22:28 fefifo_8022_1.err
-rw-rw-r--  1 jinfang jinfang  31M Oct 31 22:09 fefifo_8022_1.faa
-rw-rw-r--  1 jinfang jinfang  83M Oct 31 22:09 fefifo_8022_1.ffn
-rw-rw-r--  1 jinfang jinfang  22K Oct 31 22:21 fefifo_8022_1.fixedproducts
-rw-rw-r--  1 jinfang jinfang  98M Oct 31 21:50 fefifo_8022_1.fna
-rw-rw-r--  1 jinfang jinfang  99M Oct 31 22:09 fefifo_8022_1.fsa
-rw-rw-r--  1 jinfang jinfang 222M Oct 31 22:24 fefifo_8022_1.gbf
-rw-rw-r--  1 jinfang jinfang 142M Oct 31 22:09 fefifo_8022_1.gff
-rw-rw-r--  1 jinfang jinfang 692K Oct 31 22:29 fefifo_8022_1.log
-rw-rw-r--  1 jinfang jinfang 406M Oct 31 22:22 fefifo_8022_1.sqn
-rw-rw-r--  1 jinfang jinfang  26M Oct 31 22:09 fefifo_8022_1.tbl
-rw-rw-r--  1 jinfang jinfang  12M Oct 31 22:09 fefifo_8022_1.tsv
-rw-rw-r--  1 jinfang jinfang  131 Oct 31 22:09 fefifo_8022_1.txt
-rw-rw-r--  1 jinfang jinfang 145K Oct 31 22:24 fefifo_8022_1.val
```

**Module 2: run_dbcan annotation (Fig. 3) to obtain CAZymes, CGCs, and substrates**

**CRITICAL STEP**

Users can skip P5 and P6, and directly run P7 (much slower though), if they want to predict not only CAZymes and CGCs, but also substrates.

**P5| CAZyme annotation at family level (TIMING ~10min)**

*$ run_dbcan prokka_fefifo_8022_1/fefifo_8022_1.faa protein --hmm_cpu 32 --out_dir fefifo_8022_1.CAZyme --tools hmmer --db_dir db*

*$ run_dbcan prokka_fefifo_8022_7/fefifo_8022_7.faa protein --hmm_cpu 32 --out_dir fefifo_8022_7.CAZyme --tools hmmer --db_dir db*

Two arguments are required for run_dbcan: the input sequence file (faa files) and the sequence type (protein). By default, run_dbcan will use three methods (HMMER vs dbCAN HMMdb, DIAMOND vs CAZy, HMMER vs dbCAN-sub HMMdb) for CAZyme annotation (**Table 1, Fig. 2**). This default setting is equivalent to the use *--tools all* parameter (**Box 5**). Here we only invoke the HMMER vs dbCAN HMMdb for CAZyme annotation at the family level.

**Box 5| CAZyme annotation with default setting**

If the *--tools* parameter is not set, it is the default setting, which is the same as *--tools all*. This will take much longer time to finish (~5h) due to the large size of dbCAN-sub HMMdb (used for substrate prediction for CAZymes, see **Table 1**).

*$ run_dbcan prokka_fefifo_8022_1/fefifo_8022_1.faa protein --out_dir fefifo_8022_1.CAZyme --dia_cpu 32 --hmm_cpu 32 --dbcan_thread 32 [--tools all]*

*$ run_dbcan prokka_fefifo_8022_7/fefifo_8022_7.faa protein --out_dir fefifo_8022_7.CAZyme --dia_cpu 32 --hmm_cpu 32 --dbcan_thread 32 [--tools all]*

The sequence type can be protein, prok, meta. If the input sequence file contains metagenomic contig sequences (fna file), the sequence type has to be meta, and prodigal will be called to predict genes.

*$ run_dbcan prokka_fefifo_8022_1/fefifo_8022_1.fna meta --out_dir fefifo_8022_1.CAZyme --dia_cpu 32 --hmm_cpu 32 --dbcan_thread 32*

*$ run_dbcan prokka_fefifo_8022_7/fefifo_8022_7.fna meta --out_dir fefifo_8022_7.CAZyme --dia_cpu 32 --hmm_cpu 32 --dbcan_thread 32*

**P6| CGC prediction (TIMING ~15 min)**

The following commands will re-run run_dbcan to not only predict CAZymes but also CGCs with protein faa and gene location gff files.

*$ run_dbcan prokka_fefifo_8022_1/fefifo_8022_1.faa protein --tools hmmer --tf_cpu 32 --stp_cpu 32 -c prokka_fefifo_8022_1/fefifo_8022_1.gff --out_dir fefifo_8022_1.PUL --dia_cpu 32 --hmm_cpu 32*

*$ run_dbcan prokka_fefifo_8022_7/fefifo_8022_7.faa protein --tools hmmer --tf_cpu 32 --stp_cpu 32 -c prokka_fefifo_8022_7/fefifo_8022_7.gff --out_dir fefifo_8022_7.PUL --dia_cpu 32 --hmm_cpu 32*

As mentioned above (**Table 1, Fig. 2**), CGC prediction is a featured function added into dbCAN2 in 2018. To identify CGCs with the protein sequence type, a gene location file (gff) must be provided together. If the input sequence type is prok or meta, meaning users only have contig fna files, the CGC prediction can be activated by setting *-c cluster*.

**! CAUTION**

If the users would like to create their own gff file (instead of using Prokka or Prodigal), it is important to make sure the value of ID attribute in the gff file matches the protein ID in the protein faa file.

**?Troubleshooting**

If no result is found in CGC output file, it is most likely because the sequence IDs in gff file and faa file do not match. Another less likely reason is that the contigs are too short and fragmented and not suitable for CGC prediction.

**P7| Substrate prediction for CAZymes and CGCs (TIMING ~5h)**

The following commands will re-run run_dbcan to predict CAZymes, CGCs, and their substrates with the *--cgc_substrate* parameter.

*$ run_dbcan prokka_fefifo_8022_1/fefifo_8022_1.faa protein --dbcan_thread 32 --tf_cpu 32 --stp_cpu 32 -c prokka_fefifo_8022_1/fefifo_8022_1.gff --cgc_substrate --hmm_cpu 32 --out_dir fefifo_8022_1.dbCAN --dia_cpu 32*

*$ run_dbcan prokka_fefifo_8022_7/fefifo_8022_7.faa protein --dbcan_thread 32 --tf_cpu 32 --stp_cpu 32 -c prokka_fefifo_8022_7/fefifo_8022_7.gff --cgc_substrate --hmm_cpu 32 --out_dir fefifo_8022_7.dbCAN --dia_cpu 32*

**! CAUTION**

The above commands do not set the *--tools* parameter, which means all three methods for CAZyme annotation will be activated (**Box 5**). Because dbCAN-sub HMMdb (for CAZyme substrate prediction) is 200 times larger than dbCAN HMMdb, the runtime will be much longer. Users can specify --tools hmmer, so that the HMMER search against dbCAN-sub will be disabled. However, this will turn off the substrate prediction for CAZymes and CGCs based on CAZyme substrate majority voting. Consequently, the substrate prediction will be solely based on homology search against PULs in dbCAN-PUL (**Fig. 1, Table 1**).

*$ run_dbcan prokka_fefifo_8022_1/fefifo_8022_1.faa protein --tools hmmer --stp_cpu 32 -c prokka_fefifo_8022_1/fefifo_8022_1.gff --cgc_substrate --out_dir fefifo_8022_1.PUL.Sub --dia_cpu 32 --hmm_cpu 32 --tf_cpu 32*

*$ run_dbcan prokka_fefifo_8022_7/fefifo_8022_7.faa protein --tools hmmer --stp_cpu 32 -c prokka_fefifo_8022_7/fefifo_8022_7.gff --cgc_substrate --out_dir fefifo_8022_7.PUL.Sub --dia_cpu 32 --hmm_cpu 32 --tf_cpu 32*

**Box 6| Example output folder content of run_dbcan substrate prediction**

In the fefifo_8022_1.dbCAN directory (<https://bcb.unl.edu/dbCAN_tutorial/dataset2-Wastyk2021/fefifo_8022_1.dbCAN/>), a total of 17 files and 1 folder are generated:

```
-rw-rw-r--  1 jinfang jinfang  39M Nov  1 22:18 PUL_blast.out
-rw-rw-r--  1 jinfang jinfang 3.1M Nov  1 22:15 CGC.faa
-rw-rw-r--  1 jinfang jinfang 6.9M Nov  1 22:15 cgc.gff
-rw-rw-r--  1 jinfang jinfang 702K Nov  1 22:15 cgc.out
-rw-rw-r--  1 jinfang jinfang 321K Nov  1 22:15 cgc_standard.out
-rw-rw-r--  1 jinfang jinfang 1.5M Nov  1 22:15 cgc_standard.out.json
-rw-rw-r--  1 jinfang jinfang 556K Nov  1 22:14 dbcan-sub.hmm.out
-rw-rw-r--  1 jinfang jinfang 345K Nov  1 22:14 diamond.out
-rw-rw-r--  1 jinfang jinfang 455K Nov  1 22:14 dtemp.out
-rw-rw-r--  1 jinfang jinfang 298K Nov  1 22:14 hmmer.out
-rw-rw-r--  1 jinfang jinfang 270K Nov  1 22:15 overview.txt
-rw-rw-r--  1 jinfang jinfang 1.1M Nov  1 22:15 stp.out
-rw-rw-r--  1 jinfang jinfang  54K Nov  1 22:18 substrate.out
drwxrwxr-x  2 jinfang jinfang  32K Nov  2 09:48 synteny.pdf
-rw-rw-r--  1 jinfang jinfang 288K Nov  1 22:14 tf-1.out
-rw-rw-r--  1 jinfang jinfang 237K Nov  1 22:14 tf-2.out
-rw-rw-r--  1 jinfang jinfang 804K Nov  1 22:15 tp.out
-rw-rw-r--  1 jinfang jinfang  31M Nov  1 21:07 uniInput
```

**PUL_blast.out**: BLAST results between CGCs and PULs.

**CGC.faa**: CGC Fasta sequences.

**cgc.gff**: reformatted from the user input gff file by marking CAZymes, TFs, TCs, and STPs.

**cgc.out**: raw output of CGC predictions.

**cgc_standard.out**: simplified version of cgc.out for easy parsing in TSV format. An example row has the following columns:

9. CGC_id: *CGC1*
10. type: *CAZyme*
11. contig_id: *k141_32617*
12. gene_id: *fefifo_8022_1_00137*
13. start: *1755*
14. end: *3332*
15. strand: *-*
16. annotation: *GH13*

*Explanation: the gene fefifo_8022_1_00137 encodes a GH13 CAZyme in the CGC1 of the contig k141_32617. CGC1 also has other genes, which are provided in other rows. fefifo_8022_1_00137 is on the negative strand of k141_32617 from 1755 to 3332. The type can be one of the four signature gene types (CAZymes, TCs, TFs, STPs) or the null type (not annotated as one of the four signature genes).*

**cgc_standard.out.json**: JSON format of cgc_standard.out.

**dbcan-sub.hmm.out**: HMMER search result against dbCAN-sub HMMdb, including a column with CAZyme substrates extracted from *fam-substrate-mapping-08012023.tsv*.

**diamond.out**: DIAMOND search result against the CAZy annotated protein sequences (*CAZyDB.07262023.fa*).

**dtemp.out**: temporary file.

**hmmer.out**: HMMER search result against dbCAN HMMdb.

**overview.txt**: summary of CAZyme annotation from three methods in TSV format. An example row has the following columns:

7. Gene_ID: *fefifo_8022_1_00719*
8. EC#: *PL8_e13:2*
9. dbCAN: *PL8_2(368-612)*
10. dbCAN_sub: *PL8_e13*
11. DIAMOND: *PL8_2*
12. #ofTools: *3*

*Explanation: the protein fefifo_8022_1_00719 is annotated by 3 tools to be a CAZyme: (1) PL8_2 (CAZy defined subfamily 2 of PL8) by HMMER vs dbCAN HMMdb with a domain range from aa position 368 to 612, (2) PL8_e13 (eCAMI defined subfamily e13; e indicates it is from eCAMI not CAZy) by HMMER vs dbCAN-sub HMMdb (derived from eCAMI subfamilies), and (3) PL8_2 by DIAMOND vs CAZy annotated protein sequences. The second column 4.2.2.20:2 is extracted from eCAMI, meaning that the eCAMI subfamily PL8_e13 contains two member proteins which have an EC 4.2.2.20 according to CAZy. In most cases, the 3 tools will have the same CAZyme family assignment. When they give different assignment. We recommend a preference order: dbCAN > eCAMI/dbCAN-sub > DIAMOND. See our dbCAN2 paper^1^, dbCAN3 paper^2^, and eCAMI^3^ for more details.*

*Note: If users invoked the --use_signalP parameter when running run_dbcan, there will be an additional column called signal in the overview.txt.*

**stp.out**: HMMER search result against the MiST^4^ compiled signal transduction protein HMMs from Pfam.

**tf-1.out**: HMMER search result against the DBD^5^ compiled transcription factor HMMs from Pfam ^6^.

**tf-2.out**: HMMER search result against the DBD compiled transcription factor HMMs from Superfamily ^7^.

**tp.out**: DIAMOND search result against the TCDB ^8^ annotated protein sequences.

**substrate.out**: summary of substrate prediction results for CGCs in TSV format from two approaches^2^ (dbCAN-PUL blast search and dbCAN-sub majority voting). An example row has the following columns:

8. CGC_ID: *k141_31366|CGC2*
9. Best hit PUL_ID in dbCAN-PUL: *PUL0008*
10. Substrate of the hit PUL: *fructan*
11. Sum of bitscores for homologous gene pairs between CGC and PUL: *6132.0*
12. Types of homologous gene pairs: *CAZyme-CAZyme;CAZyme-CAZyme;TC-TC;CAZyme-CAZyme;CAZyme-CAZyme;TC-TC*
13. Substrate predicted by majority voting of CAZymes in CGC: *fructan*
14. Voting score: *2.0*

*The CGC1 of contig k141_31366 has its best hit PUL0008 (from PUL_blast.out) with fructan as substrate (from dbCAN-PUL_12-12-2023.xlsx). Six signature genes are matched between k141_31366|CGC2 and PUL0008 (from PUL_blast.out): four are CAZymes and the other two are TCs. The sum of blast bitscores of the six homologous pairs (CAZyme-CAZyme, CAZyme-CAZyme, TC-TC, CAZyme-CAZyme, CAZyme-CAZyme and TC-TC) is 6132.0. Hence, the substrate of k141_31366|CGC2 is predicted to be fructan according to dbCAN-PUL blast search. The last two columns are based on the dbCAN-sub result (dbcan-sub.hmm.out), according to which two CAZymes in k141_31366|CGC2 are predicted to have fructan substrate. The voting score is thus 2.0, so that according to the majority voting rule, k141_31366|CGC2 is predicted to have a fructan substrate.*

*Note: for many CGCs, only one of the two approaches produces substrate prediction. In some cases, the two approaches produce different substrate assignments. We recommend a preference order: dbCAN-PUL blast search > dbCAN-sub majority voting. See our dbCAN3 paper*^2^ *for more details.*

**synteny.pdf**: a folder with syntenic block alignment plots between all CGCs and PULs.

**uniInput**: renamed Fasta file from input protein sequence file.

**Module 3: Read mapping (Fig. 3) to calculate abundance for CAZyme families, subfamilies, CGCs, and substrates**

**P8| Read mapping to all CDS of each sample (TIMING ~10 min)**

*$ bwa index prokka_fefifo_8022_1/fefifo_8022_1.ffn*

*$ bwa index prokka_fefifo_8022_7/fefifo_8022_7.ffn*

*$ mkdir samfiles*

*$ bwa mem -t 32 -o samfiles/fefifo_8022_1.CDS.sam prokka_fefifo_8022_1/fefifo_8022_1.ffn fefifo_8022_1__shotgun_1_val_1.fq.gz fefifo_8022_1__shotgun_2_val_2.fq.gz*

*$ bwa mem -t 32 -o samfiles/fefifo_8022_7.CDS.sam prokka_fefifo_8022_7/fefifo_8022_7.ffn fefifo_8022_7__shotgun_1_val_1.fq.gz fefifo_8022_7__shotgun_2_val_2.fq.gz*

Reads are mapped to the ffn files from Prokka.

**P9| Read mapping to all contigs of each sample (TIMING ~10min)**

*$ bwa index megahit_fefifo_8022_1/fefifo_8022_1.contigs.fa*

*$ bwa index megahit_fefifo_8022_7/fefifo_8022_7.contigs.fa*

*$ bwa mem -t 32 -o samfiles/fefifo_8022_1.sam megahit_fefifo_8022_1/fefifo_8022_1.contigs.fa fefifo_8022_1__shotgun_1_val_1.fq.gz fefifo_8022_1__shotgun_2_val_2.fq.gz*

*$ bwa mem -t 32 -o samfiles/fefifo_8022_7.sam megahit_fefifo_8022_7/fefifo_8022_7.contigs.fa fefifo_8022_7__shotgun_1_val_1.fq.gz fefifo_8022_7__shotgun_2_val_2.fq.gz*

Reads are mapped to the contig files from MEGAHIT.

**P10| Sort SAM files by coordinates (TIMING ~6min)**

*$ cd samfiles*

*$ samtools sort -@ 32 -o fefifo_8022_1.CDS.bam fefifo_8022_1.CDS.sam*

*$ samtools sort -@ 32 -o fefifo_8022_7.CDS.bam fefifo_8022_7.CDS.sam*

*$ samtools sort -@ 32 -o fefifo_8022_1.bam fefifo_8022_1.sam*

*$ samtools sort -@ 32 -o fefifo_8022_7.bam fefifo_8022_7.sam*

*$ rm -rf *sam*

*$ cd ..*

**P11| Read count calculation for all proteins of each sample using Bedtools (TIMING ~1min)**

*$ mkdir fefifo_8022_1_abund && cd fefifo_8022_1_abund*

*$ seqkit fx2tab -l -n -i ../prokka_fefifo_8022_1/fefifo_8022_1.ffn | awk '{print $1"\t"$2}' > fefifo_8022_1.length*

*$ seqkit fx2tab -l -n -i ../prokka_fefifo_8022_1/fefifo_8022_1.ffn | awk '{print $1"\t"0"\t"$2}' > fefifo_8022_1.bed*

*$ bedtools coverage -g fefifo_8022_1.length -sorted -a fefifo_8022_1.bed -counts -b ../samfiles/fefifo_8022_1.CDS.bam > fefifo_8022_1.depth.txt*

*$ cd .. && mkdir fefifo_8022_7_abund && cd fefifo_8022_7_abund*

*$ seqkit fx2tab -l -n -i ../prokka_fefifo_8022_7/fefifo_8022_7.ffn | awk '{print $1"\t"$2}' > fefifo_8022_7.length*

*$ seqkit fx2tab -l -n -i ../prokka_fefifo_8022_7/fefifo_8022_7.ffn | awk '{print $1"\t"0"\t"$2}' > fefifo_8022_7.bed*

*$ bedtools coverage -g fefifo_8022_7.length -sorted -a fefifo_8022_7.bed -counts -b ../samfiles/fefifo_8022_7.CDS.bam > fefifo_8022_7.depth.txt*

*$ cd ..*

Read counts are saved in *depth.txt* files of each sample.

**P12| Read count calculation for a given region of contigs using Samtools (TIMING ~1min)**

*$ cd fefifo_8022_1_abund*

*$ samtools index ../samfiles/fefifo_8022_1.bam*

*$ samtools depth -r k141_2168:4235-19858 ../samfiles/fefifo_8022_1.bam > fefifo_8022_1.cgc.depth.txt*

*$ cd ..*

The parameter *-r k141_2168:4235-19858* specifies a region in a contig. For any CGC, its positional range can be found in the file cgc_standard.out produced by run_dbcan (**Box 6**). The *depth.txt* files contain the raw read counts for the specified region.

**! CAUTION**

The contig IDs are automatically generated by MEGAHIT. There is a small chance that a same contig ID appears in both samples. However, the two contigs in the two samples do not match each other even the ID is the same. For example, the contig ID k141_2168 is most likely only found in the fefifo_8022_1 sample. Even if there is a k141_2168 in fefifo_8022_7, the actual contigs in two samples are different.

**P13| dbcan_utils to calculate the abundance of CAZyme families, subfamilies, CGCs, and substrates (TIMING ~1min)**

*$ dbcan_utils fam_abund -bt* *fefifo_8022_1.depth.txt -i ../fefifo_8022_1.dbCAN -a TPM*

*$ dbcan_utils fam_substrate_abund -bt fefifo_8022_1.depth.txt -i ../fefifo_8022_1.dbCAN -a TPM*

*$ dbcan_utils CGC_abund -bt fefifo_8022_1.depth.txt -i ../fefifo_8022_1.dbCAN -a TPM*

*$ dbcan_utils CGC_substrate_abund -bt fefifo_8022_1.depth.txt -i ../fefifo_8022_1.dbCAN -a TPM*

*$ cd .. && cd fefifo_8022_7_abund*

*$ dbcan_utils fam_abundfam_substrate_abund -bt fefifo_8022_7.depth.txt -i ../fefifo_8022_7.dbCAN -a TPM*

*$ dbcan_utils fam_substrate_abund -bt fefifo_8022_7.depth.txt -i ../fefifo_8022_7.dbCAN -a TPM*

*$ dbcan_utils CGC_abund -bt fefifo_8022_7.depth.txt -i ../fefifo_8022_7.dbCAN -a TPM*

*$ dbcan_utils CGC_substrate_abund -bt fefifo_8022_7.depth.txt -i ../fefifo_8022_7.dbCAN -a TPM*

*cd ..*

We developed a set of Python scripts as dbcan_utils (included in the run_dbcan package) to take the raw read counts for all genes as input and output the normalized abundances (**Box 7**) of CAZyme families, subfamilies, CGCs, and substrates (**Fig. 4**). The parameter *-a TPM* can also be two other metrics: RPM, or RPKM^11^.

RPKM = # of mapped reads to a gene G / [(total # of mapped reads to all genes /106) x (gene G length/1000)]

RPM = # of mapped reads to a gene G / (total # of mapped reads to all genes/106).

TPM = [# of mapped reads to a gene G / (gene G length/1000)] / sum [# of mapped reads to each gene / (the gene length/1000)].

**Box 7| Example output of dbcan_utils**

As an example, fefifo_8022_1_abund (<https://bcb.unl.edu/dbCAN_tutorial/dataset2-Wastyk2021/fefifo_8022_1_abund/>) has 7 TSV files:

```
-rw-rw-r--  1 jinfang jinfang 178K Jan  2 04:08 CGC_abund.out
-rw-rw-r--  1 jinfang jinfang 3.3K Jan  2 04:08 CGC_substrate_majority_voting.out
-rw-rw-r--  1 jinfang jinfang  12K Jan  2 04:08 CGC_substrate_PUL_homology.out
-rw-rw-r--  1 jinfang jinfang 2.5K Jan  2 04:08 EC_abund.out
-rw-rw-r--  1 jinfang jinfang 4.1K Jan  2 04:08 fam_abund.out
-rw-rw-r--  1 jinfang jinfang  42K Jan  2 04:08 fam_substrate_abund.out
-rw-rw-r--  1 jinfang jinfang  26K Jan  2 04:08 subfam_abund.out
```

Explanation of columns in these TSV files is as follows:

**fam_abund.out**: CAZy family (from HMMER vs dbCAN HMMdb), sum of TPM, # of CAZymes in the family

**subfam_abund.out:** eCAMI subfamily (from HMMER vs dbCAN-sub HMMdb), sum of TPM, # of CAZymes in the subfamily

**EC_abund.out:** EC number (extracted from dbCAN-sub subfamily), sum of TPM, # of CAZymes with the EC

**fam_substrate_abund.out:** Substrate (from HMMER vs dbCAN-sub HMMdb), sum of TPM (all CAZymes in this substrate group), GeneID (all CAZyme IDs in this substrate group)

**CGC_abund.out:** CGC_ID (e.g., k141_338400|CGC1), mean of TPM (all genes in the CGC), Seq_IDs (IDs of all genes in the CGC), TPM (of all genes in the CGC), Families (CAZyme family or other signature gene type of all genes in the CGC)

**CGC_substrate_PUL_homology.out:** Substrate (from dbCAN-PUL blast search), sum of TPM, CGC_IDs (all CGCs predicted to have the substrate from dbCAN-PUL blast search), TPM (of CGCs in this substrate group)

**CGC_substrate_majority_voting.out:** Substrate (from dbCAN-sub majority voting), sum of TPM, CGC_IDs (all CGCs predicted to have the substrate from dbCAN-sub majority voting), TPM (of CGCs in this substrate group)

**Module 4: dbcan_plot for data visualization (Fig. 3) of abundances of CAZymes, CGCs, and substrates (TIMING variable)**

**CRITICAL STEP**

To visualize the CAZyme annotation result, we provide a set of Python scripts as dbcan_plot to make publication quality plots with the dbcan_utils results as the input. The dbcan_plot scripts are included in the run_dbcan package. Once the plots are made in PDF format, they can be transferred to users’ Windows or Mac computers for visualization.

Five data folders will be needed as the input for dbcan_plot: (i) two abundance folders fefifo_8022_1_abund and fefifo_8022_7_abund, (ii) two CAZyme annotation folders fefifo_8022_1.dbCAN and fefifo_8022_7.dbCAN, and (iii) the dbCAN-PUL folder (under the db folder, released from dbCAN-PUL.tar.gz).

**P14| Heatmap for CAZyme substrate abundance across samples (Fig. S4B) (TIMING 1min)**

*$ dbcan_plot heatmap_plot --samples fefifo_8022_1,fefifo_8022_7 -i fefifo_8022_1_abund/* *fam_substrate_abund.out,* *fefifo_8022_7_abund/fam_substrate_abund.out --show_abund --top 20*

Here we plot the top 20 substrates in the two samples. The input files are the two CAZyme substrate abundance files calculated based on dbCAN-sub result. The default heatmap is ranked by substrate abundances. To rank the heatmap according to abundance profile using the function clustermap of seaborn package, users can invoke the --cluster_map parameter.

**P15| Barplot for CAZyme family abundance across samples (Fig. S4C) (TIMING 1min)**

*$ dbcan_plot bar_plot --samples fefifo_8022_1,fefifo_8022_7 --vertical_bar --top 20 -i fefifo_8022_1_abund/fam_abund.out,fefifo_8022_7_abund/fam_abund.out*

Users can choose to generate a barplot instead of heatmap using the bar_plot method.

**P16| Synteny plot between a CGC and its best PUL hit with read mapping coverage to CGC (Fig. S4A) (TIMING 1min)**

*$ dbcan_plot CGC_synteny_coverage_plot -i fefifo_8022_1.dbCAN --cgcid ‘k141_2168|CGC1' --readscount fefifo_8022_1_abund/fefifo_8022_1.cgc.depth.txt*

The fefifo_8022_1.dbCAN folder contains the PUL_blast.out file. Using this file, the cgc_standard.out file, and the best PUL’s gff file in dbCAN-PUL.tar.gz, the CGC_synteny_plot method will create the CGC-PUL synteny plot. The –cgcid parameter is required to specify which CGC to be plotted (‘k141_2168|CGC1' in this example). The *fefifo_8022_1.cgc.depth.txt* file is used to plot the read mapping coverage.

If users only want to plot the CGC structure:

*$ dbcan_plot CGC_plot -i fefifo_8022_1.dbCAN --cgcid ‘k141_2168|CGC1'*

If users only want to plot the CGC structure plus the read mapping coverage:

*$ dbcan_plot CGC_coverage_plot -i fefifo_8022_1.dbCAN --cgcid ‘k141_2168|CGC1’--readscount fefifo_8022_1_abund/fefifo_8022_1.cgc.depth.txt*

If users only want to plot the synteny between the CGC and PUL:

*$ dbcan_plot CGC_synteny_plot -i fefifo_8022_1.dbCAN --cgcid ‘k141_2168|CGC1’*

**! CAUTION**

The CGC IDs in different samples do not match each other. For example, specifying -i fefifo_8022_1.dbCAN is to plot the 'k141_2168|CGC1’ in the fefifo_8022_1 sample. The ‘k141_2168|CGC1’ in the fefifo_8022_7 sample most likely does not exist, and even it does, the CGC has a different sequence even if the ID is the same.

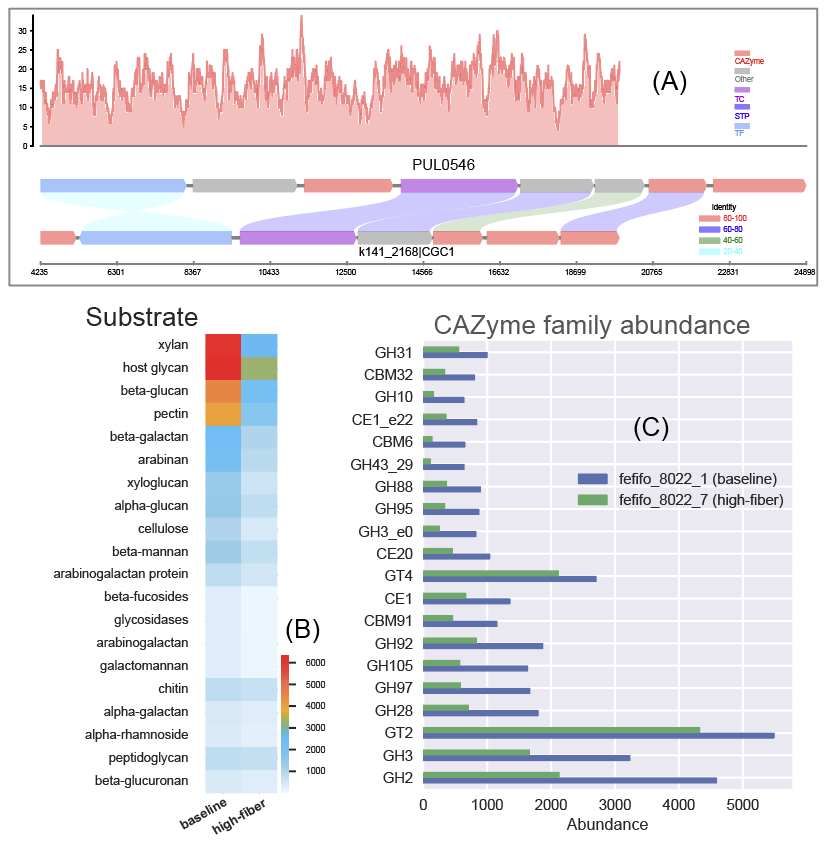

***Fig. S4.*** ***Data visualization of abundances of CAZymes, CGCs, and substrates****. (****A****) Synteny plot between an example CGC (CGC1 of contig k141_2168 from the fefifo_8022_1 sample) and its best PUL hit (PUL0546 from dbCAN-PUL, with experimentally verified substrate arabinogalactan) with read mapping coverage plot (y-axis is the read depth) shown in the bottom. (****B****) Heatmap of CAZyme substrate abundance in the two samples (Table S2) calculated based on dbCAN-sub search results and substrate mapping. Abundance values are TPM. (****C****) Barplot of 20 CAZyme family abundance in the two samples calculated based on dbCAN search results.*

**Procedure of individual sample assembly for Priest2023**

The Priest2023 dataset was published in 2023 from a study of glycan utilization in the particulate and dissolved organic matter pools (POM and DOM) in the ocean waters of the Arctic during late summer^13^. This study applied PacBio HiFi long read sequencing for metagenomes (MGs) of 8 samples (4 sites of two depth) and Illumina short read sequencing for metatranscriptomes (MTs) of 4 samples (4 sites). It investigated CAZyme abundance and transcription profiles as well as inferred glycan degradation activities by comparing MGs and MTs from samples of different sites and depth. The sites are categorized into above-slope, above-shelf and open-ocean groups, and the water depth includes surface water (SRF) and bottom of the surface mixed layer (BML). CAZyme abundance and expression showed different profiles for the degradation of complex algal glycans such as sulfated fucan, laminarin and xylan in different site groups and water depths.

For this protocol, we will select two MG samples (HiFi DNA long reads, FRAM_WSC20_S25_SRF_MG and FRAM_WSC20_S25_BML_MG) and two MT samples (FRAM_WSC20_S25_SRF_MT and FRAM_WSC20_S25_BML_MT, paired-end 2x151bp reads) for CAZyme annotation (**Table S3**). We will assemble HiFi long MG reads into contigs, which will be used to predict CAZymes and glycan substrates. We will map the MT reads to the contigs to calculate expression levels.

**Table S3. Summary of the example microbiome dataset**

| **Dataset** (Priest T et al., ISME Communications 2023 ^13^) | | |
| --- | --- | --- |
| Sample name | S25_SRF | S25_BML |
| Water depth | surface water | bottom of mixed layer |
| MG SRA ID | ERR10947144 | ERR10946524 |
| MG HiFi read count | 1,446,226 | 1,643,900 |
| MT SRA ID | ERR10949699 | ERR10949700 |
| MT Illumina read count | 73,377,699 | 87,034,266 |

The procedure has 4 modules (**Fig. 3**) and 17 steps (P1-P17).

**Module 1: HiFi long reads processing (Fig. 3) to obtain contigs**

**P1| Check contamination (TIMING ~20min)**

*$ wget https://bcb.unl.edu/dbCAN_tutorial/dataset3-Priest2023/BML_MG.fastq.gz*

*$ wget https://bcb.unl.edu/dbCAN_tutorial/dataset3-Priest2023/SRF_MG.fastq.gz*

*$ wget https://bcb.unl.edu/dbCAN_tutorial/dataset3-Priest2023/BML_MT_1.fastq.gz*

*$ wget https://bcb.unl.edu/dbCAN_tutorial/dataset3-Priest2023/BML_MT_2.fastq.gz*

*$ wget https://bcb.unl.edu/dbCAN_tutorial/dataset3-Priest2023/SRF_MT_1.fastq.gz*

*$ wget https://bcb.unl.edu/dbCAN_tutorial/dataset3-Priest2023/SRF_MT_2.fastq.gz*

*$ kraken2 --threads 32 --quick --paired --db K2 --report SRF_MT.kreport --output SRF_MT.kraken.output SRF_MT_1.fastq.gz SRF_MT_2.fastq.gz*

*$ kraken2 --threads 32 --quick --paired --db K2 --report BML_MT.kreport --output BML_MT.kraken.output BML_MT_1.fastq.gz BML _MT_2.fastq.gz*

*$ kraken2 --threads 32 --quick --paired --db K2 --report SRF_MG.kreport --output SRF_MG.kraken.output SRF_MG_1.fastq.gz SRF_MG_2.fastq.gz*

*$ kraken2 --threads 32 --quick --paired --db K2 --report BML_MG.kreport --output BML_MG.kraken.output BML_MG_1.fastq.gz BML_MG_2.fastq.gz*

Kraken2 found much contamination (**Box 1**) from human in the Priest2023 data. Consequently, human reads need to be removed before assembly.

Reads can be aligned to the reference genomes of potential contamination source organisms to remove the aligned reads. The most common source in microbiome studies is from human.

**Box 1| Example of Kraken2 output files**

The *kreport* files can be examined to identify potential contamination source organisms.

```
-rw-rw-r-- 1 jinfang jinfang   54M Dec 27 07:42 BML_MG.kraken.output
-rw-rw-r-- 1 jinfang jinfang  1.2M Dec 27 07:42 BML_MG.kreport
-rw-rw-r-- 1 jinfang jinfang  3.4G Dec 27 08:01 BML_MT.kraken.output
-rw-rw-r-- 1 jinfang jinfang 1023K Dec 27 08:02 BML_MT.kreport
-rw-rw-r-- 1 jinfang jinfang   61M Dec 27 07:39 SRF_MG.kraken.output
-rw-rw-r-- 1 jinfang jinfang  1.2M Dec 27 07:39 SRF_MG.kreport
-rw-rw-r-- 1 jinfang jinfang  2.6G Dec 27 07:50 SRF_MT.kraken.output
-rw-rw-r-- 1 jinfang jinfang  1.1M Dec 27 07:51 SRF_MT.kreport
```

**P2| Remove contamination reads from human (TIMING ~40min)**

From the Kraken2 output files, we identified humans as the contamination source, we can use the following commands to remove the contamination reads by aligning reads to the human reference genome.

*$ mkdir hg38 && cd hg38 && wget https://ftp.ensembl.org/pub/release-110/fasta/homo_sapiens/dna/Homo_sapiens.GRCh38.dna.primary_assembly.fa.gz*

*$ cd .. && mkdir contamination && cd contamination*

*$ minimap2 -a -x map-hifi -MD -t 32 -o SRF_MG.hg38.sam ../hg38/Homo_sapiens.GRCh38.dna.primary_assembly.fa.gz ../SRF_MG.fastq.gz*

*$ minimap2 -a -x map-hifi -MD -t 32 -o BML_MG.hg38.sam ../hg38/Homo_sapiens.GRCh38.dna.primary_assembly.fa.gz ../BML_MG.fastq.gz*

*$ samtools fastq -f 4 -@ 32 -0 ../SRF_MG.clean.fq.gz SRF_MG.hg38.sam*

*$ samtools fastq -f 4 -@ 32 -0 ../BML_MG.clean.fq.gz BML_MG.hg38.sam*

*$ bwa mem ../hg38/hg38 ../SRF_MT_1.fastq.gz ../SRF_MT_2.fastq.gz -t 32 -o SRF_MT.hg38.sam*

*$ bwa mem ../hg38/hg38 ../BML_MT_1.fastq.gz ../BML_MT_2.fastq.gz -t 32 -o BML_MT.hg38.sam*

*$ samtools fastq -f 12 -@ 32 -1 ../SRF_MT_1.clean.fq.gz -2 ../SRF_MT_2.clean.fq.gz SRF_MT.hg38.sam*

*$ samtools fastq -f 12 -@ 32 -1 ../BML_MT_1.clean.fq.gz -2 ../BML_MT_2.clean.fq.gz BML_MT.hg38.sam*

*$ cd ..*

**P3| Trim adapter and low-quality reads (TIMING ~20min)**

*$ trim_galore --illumina -j 8 --paired BML_MT_1.clean.fastq.gz BML_MT_2.clean.fastq.gz*

*$ trim_galore --illumina -j 8 --paired SRF_MT_1.clean.fastq.gz SRF_MT_2.clean.fastq.gz*

The HiFi long reads do not need to be trimmed. Hence, this step only applies to MT illumina short read data. We specified --illumina to indicate that the reads were generated using the Illumina sequencing platform. Nonetheless, trim_galore possesses the ability to automatically detect the adapter, providing flexibility in adapter handling for users who may know the specific sequencing platform. Details of trimming are available in the trimming report file (**Box 2**).

**Box 2| Example output of trim_galore**

In addition to the trimmed read files, Trim_galore also generates a trimming report file. The trimming report contains details on read trimming, such as the number of trimmed reads.

*-rw-rw-r-- 1 jinfang jinfang 4.2K Dec 28 21:56 BML_MT_1.clean.fq.gz_trimming_report.txt*

*-rw-rw-r-- 1 jinfang jinfang 2.3G Dec 28 22:05 BML_MT_1.clean_val_1.fq.gz*

*-rw-rw-r-- 1 jinfang jinfang 4.7K Dec 28 22:05 BML_MT_2.clean.fq.gz_trimming_report.txt*

*-rw-rw-r-- 1 jinfang jinfang 3.0G Dec 28 22:05 BML_MT_2.clean_val_2.fq.gz*

*-rw-rw-r-- 1 jinfang jinfang 4.9K Dec 28 10:07 SRF_MT_1.clean.fq.gz_trimming_report.txt*

*-rw-rw-r-- 1 jinfang jinfang 2.7G Dec 28 10:19 SRF_MT_1.clean_val_1.fq.gz*

*-rw-rw-r-- 1 jinfang jinfang 5.1K Dec 28 10:19 SRF_MT_2.clean.fq.gz_trimming_report.txt*

*-rw-rw-r-- 1 jinfang jinfang 3.3G Dec 28 10:19 SRF_MT_2.clean_val_2.fq.gz*

**! CAUTION**

During the trimming process, certain reads may be entirely removed due to low quality in its entirety. Using the --retain_unpaired parameter in trim_galore allows for the preservation of single-end reads. In this protocol, this option was not selected, so that both reads of a forward-revise pair were removed.

**P4| Assemble HiFi reads into metagenome (TIMING ~4h20min)**

Flye was used to assemble the HiFi long reads into contigs.

*$ flye --threads 32 --meta --pacbio-hifi BML_MG.clean.fq.gz --hifi-error 0.01 --keep-haplotypes --out-dir flye_BML_MG*

*$ flye --threads 32 --meta --pacbio-hifi SRF_MG.clean.fq.gz --hifi-error 0.01 --keep-haplotypes --out-dir flye_SRF_MG*

Flye generates two folders flye_BML_MG and flye_SRF_MG. Each folder contains 6 files and 5 sub-folders (**Box 3**), among them assembly.fasta is the final contig sequence file. We set --hifi-error 0.01, a generally accepted error rate of HiFi sequencing. Parameter --meta is set to assemble reads into metagenomes.

**Box 3| Example output of Flye**

```
drwxrwxr-x  2 jinfang jinfang 4.0K Dec 27 20:15 00-assembly
drwxrwxr-x  2 jinfang jinfang 4.0K Dec 27 20:43 10-consensus
drwxrwxr-x  2 jinfang jinfang 4.0K Dec 27 21:14 20-repeat
drwxrwxr-x  2 jinfang jinfang 4.0K Dec 27 21:16 30-contigger
drwxrwxr-x  2 jinfang jinfang 4.0K Dec 27 22:06 40-polishing
-rw-rw-r--  1 jinfang jinfang 314M Dec 27 22:06 assembly.fasta
-rw-rw-r--  1 jinfang jinfang 311M Dec 27 22:06 assembly_graph.gfa
-rw-rw-r--  1 jinfang jinfang 6.6M Dec 27 22:06 assembly_graph.gv
-rw-rw-r--  1 jinfang jinfang 867K Dec 27 22:06 assembly_info.txt
-rw-rw-r--  1 jinfang jinfang  61M Dec 27 22:06 flye.log
-rw-rw-r--  1 jinfang jinfang   92 Dec 27 22:06 params.json
```

**P5| Predict genes by Prokka (~21h)**

*$ prokka --outdir prokka_BML_MG --prefix BML_MG --addgenes --addmrna --locustag BML_MG --kingdom Bacteria --cpus 36 flye_BML_MG/assembly.fasta*

*$ prokka --outdir prokka_SRF_MG --prefix SRF_MG --addgenes --addmrna --locustag* *SRF_MG --kingdom Bacteria --cpus 36 flye_SRF_MG/assembly.fasta*

The parameter --kingdom Bacteria is required for bacterial gene prediction. To optimize performance, --CPU 36 instructs the utilization of 36 computer processors. The output files comprise of both protein and CDS sequences in Fasta format (e.g., *BML_MG.faa* and *SRF_MG.ffn* in **Box 4**).

**Box 4| Example output of Prokka**

```
-rw-rw-r--  1 jinfang jinfang 2.2M Dec 28 05:38 BML_MG.err
-rw-rw-r--  1 jinfang jinfang 105M Dec 27 23:26 BML_MG.faa
-rw-rw-r--  1 jinfang jinfang 288M Dec 27 23:26 BML_MG.ffn
-rw-rw-r--  1 jinfang jinfang 314M Dec 27 22:06 BML_MG.fna
-rw-rw-r--  1 jinfang jinfang 315M Dec 27 23:26 BML_MG.fsa
-rw-rw-r--  1 jinfang jinfang 724M Dec 28 05:39 BML_MG.gbk
-rw-rw-r--  1 jinfang jinfang 467M Dec 27 23:26 BML_MG.gff
-rw-rw-r--  1 jinfang jinfang 1.9M Dec 28 05:39 BML_MG.log
-rw-rw-r--  1 jinfang jinfang 1.5G Dec 28 05:39 BML_MG.sqn
-rw-rw-r--  1 jinfang jinfang  89M Dec 27 23:26 BML_MG.tbl
-rw-rw-r--  1 jinfang jinfang  40M Dec 27 23:26 BML_MG.tsv
-rw-rw-r--  1 jinfang jinfang  152 Dec 27 23:26 BML_MG.txt
```

**Module 2: run_dbcan annotation (Fig. 3) to obtain CAZymes, CGCs, and substrates**

**CRITICAL STEP**

Users can skip P6 and P7, and directly run P8 (much slower though), if they want to predict not only CAZymes and CGCs, but also substrates.

**P6| CAZyme annotation at family level (TIMING ~10min)**

*$ run_dbcan prokka_BML_MG/BML_MG.faa protein --hmm_cpu 32 --out_dir BML_MG.CAZyme --tools hmmer --db_dir db*

*$ run_dbcan prokka_SRF_MG/SRF_MG.faa protein --hmm_cpu 32 --out_dir SRF_MG.CAZyme --tools hmmer --db_dir db*

Two arguments are required for run_dbcan: the input sequence file (faa) and the sequence type (protein). By default, run_dbcan will use three methods (HMMER vs dbCAN HMMdb, DIAMOND vs CAZy, HMMER vs dbCAN-sub HMMdb) for CAZyme annotation (**Table 1, Fig. 2**). This default setting is equivalent to the use *--tools all* parameter (**Box 5**). Here we only invoke the HMMER vs dbCAN HMMdb for CAZyme annotation at the family level.

**Box 5| CAZyme annotation with default setting**

If the *--tools* parameter is not set, it is the default setting, which is the same as *--tools all*. This will take much longer time to finish (~5h) due to the large size of dbCAN-sub HMMdb (used for substrate prediction for CAZymes, see **Table 1**).

*$ run_dbcan prokka_BML_MG/BML_MG.faa protein --out_dir BML_MG.CAZyme --dia_cpu 32 --hmm_cpu 32 --dbcan_thread 32 --tools all*

*$ run_dbcan prokka_SRF_MG/SRF_MG.faa protein --out_dir SRF_MG.CAZyme --dia_cpu 32 --hmm_cpu 32 --dbcan_thread 32 --tools all*

The sequence type can be protein, prok, meta. If the input sequence file contains metagenomic contig sequences (fna file), the sequence type has to be meta, and prodigal will be called to predict genes.

*$ run_dbcan prokka_BML_MG/BML_MG.fna meta --out_dir BML_MG.CAZyme --dia_cpu 32 --hmm_cpu 32 --dbcan_thread 32*

*$ run_dbcan prokka_SRF_MG/SRF_MG.fna meta --out_dir SRF_MG.CAZyme --dia_cpu 32 --hmm_cpu 32 --dbcan_thread 32*

**P7| CGC prediction (TIMING ~15 min)**

The following commands will re-run run_dbcan to not only predict CAZymes but also CGCs with protein faa and gene location gff files.

*$ run_dbcan prokka_BML_MG/BML_MG.faa protein --tools hmmer --tf_cpu 32 --stp_cpu 32 -c prokka_BML_MG/BML_MG.gff --out_dir BML_MG.PUL --dia_cpu 32 --hmm_cpu 32*

*$ run_dbcan prokka_SRF_MG/SRF_MG.faa protein --tools hmmer --tf_cpu 32 --stp_cpu 32 -c prokka_SRF_MG/SRF_MG.gff --out_dir SRF_MG.PUL --dia_cpu 32 --hmm_cpu 32*

As mentioned above (**Table 1, Fig. 2**), CGC prediction is a featured function added into dbCAN2 in 2018. To identify CGCs with the protein sequence type, a gene location file (gff) must be provided together. If the input sequence type is prok or meta, meaning users only have contig fna files, the CGC prediction can be activated by setting *-c cluster*.

**! CAUTION**

If the users would like to create their own gff file (instead of using Prokka or Prodigal), it is important to make sure the value of ID attribute in the gff file matches the protein ID in the protein faa file.

**?Troubleshooting**

If no result is found in CGC output file, it is most likely because the sequence IDs in gff file and faa file do not match. Another less likely reason is that the contigs are too short and fragmented and not suitable for CGC prediction.

**P8| Substrate prediction for CAZymes and CGCs (TIMING ~5h)**

The following commands will re-run run_dbcan to predict CAZymes, CGCs, and their substrates with the *--cgc_substrate* parameter.

*$ run_dbcan prokka_BML_MG/BML_MG.faa protein --dbcan_thread 32 --tf_cpu 32 --stp_cpu 32 -c prokka_BML_MG/BML_MG.gff --cgc_substrate --hmm_cpu 32 --out_dir BML_MG.dbCAN --dia_cpu 32*

*$ run_dbcan prokka_SRF_MG/SRF_MG.faa protein --dbcan_thread 32 --stp_cpu 32 -c prokka_SRF_MG/SRF_MG.gff --cgc_substrate --out_dir SRF_MG.dbCAN --dia_cpu 32 --hmm_cpu 32 --tf_cpu 32*

**! CAUTION**

The above commands do not set the *--tools* parameter, which means all three methods for CAZyme annotation will be activated (**Box 5**). Because dbCAN-sub HMMdb (for CAZyme substrate prediction) is 200 times larger than dbCAN HMMdb, the runtime will be much longer. Users can specify --tools hmmer, so that the HMMER search against dbCAN-sub will be disabled. However, this will turn off the substrate prediction for CAZymes and CGCs based on CAZyme substrate majority voting. Consequently, the substrate prediction will be solely based on homology search against PULs in dbCAN-PUL (**Fig. 1, Table 1**).

*$ run_dbcan prokka_BML_MG/BML_MG.faa protein --tools hmmer --stp_cpu 32 -c prokka_BML_MG/BML_MG.gff --cgc_substrate --out_dir BML_MG.PUL.Sub --dia_cpu 32 --hmm_cpu 32 --tf_cpu 32*

*$ run_dbcan prokka_SRF_MG/SRF_MG.faa protein --tools hmmer --stp_cpu 32 -c prokka_SRF_MG/SRF_MG.gff --cgc_substrate --out_dir SRF_MG.PUL.Sub --dia_cpu 32 --hmm_cpu 32 --tf_cpu 32*

**Box 6| Example output folder content of run_dbcan substrate prediction**

In the output directory (<https://bcb.unl.edu/dbCAN_tutorial/dataset3-Priest2023/BML_MG.dbCAN/>), a total of 17 files and 1 folder are generated:

```
-rw-rw-r--  1 jinfang jinfang  9.6M Dec 28 10:18 PUL_blast.out
-rw-rw-r--  1 jinfang jinfang  1.8M Dec 28 10:18 CGC.faa
-rw-rw-r--  1 jinfang jinfang   26M Dec 28 10:18 cgc.gff
-rw-rw-r--  1 jinfang jinfang  450K Dec 28 10:18 cgc.out
-rw-rw-r--  1 jinfang jinfang  212K Dec 28 10:18 cgc_standard.out
-rw-rw-r--  1 jinfang jinfang 1005K Dec 28 10:18 cgc_standard.out.json
-rw-rw-r--  1 jinfang jinfang  406K Dec 28 10:11 dbcan-sub.hmm.out
-rw-rw-r--  1 jinfang jinfang  325K Dec 28 10:11 diamond.out
-rw-rw-r--  1 jinfang jinfang  332K Dec 28 10:11 dtemp.out
-rw-rw-r--  1 jinfang jinfang  220K Dec 28 10:11 hmmer.out
-rw-rw-r--  1 jinfang jinfang  240K Dec 28 10:18 overview.txt
-rw-rw-r--  1 jinfang jinfang  1.7M Dec 28 10:17 stp.out
-rw-rw-r--  1 jinfang jinfang   17K Dec 28 10:18 substrate.out
drwxrwxr-x  2 jinfang jinfang   12K Dec 28 10:19 synteny.pdf
-rw-rw-r--  1 jinfang jinfang  293K Dec 28 10:13 tf-1.out
-rw-rw-r--  1 jinfang jinfang  222K Dec 28 10:15 tf-2.out
-rw-rw-r--  1 jinfang jinfang  1.7M Dec 28 10:17 tp.out
-rw-rw-r--  1 jinfang jinfang  105M Dec 28 05:57 uniInput
```

**PUL_blast.out**: BLAST results between CGCs and PULs.

**CGC.faa**: CGC Fasta sequences.

**cgc.gff**: reformatted from the user input gff file by marking CAZymes, TFs, TCs, and STPs.

**cgc.out**: raw output of CGC predictions.

**cgc_standard.out**: simplified version of cgc.out for easy parsing in TSV format. An example row has the following columns:

17. CGC_id: *CGC1*
18. type: *CAZyme*
19. contig_id: *contig_10157*
20. gene_id: *BML_MG_01992*
21. start: *33003*
22. end: *36077*
23. strand: *+*
24. annotation: *GH2*

*Explanation: the gene BML_MG_01992 encodes a GH2 CAZyme in the CGC1 of the contig contig_10157. CGC1 also has other genes, which are provided in other rows. BML_MG_01992 is on the positive strand of contig_10157 from 33003 to 36077. The type can be one of the four signature gene types (CAZymes, TCs, TFs, STPs) or the null type (not annotated as one of the four signature genes).*

**cgc_standard.out.json**: JSON format of cgc_standard.out.

**dbcan-sub.hmm.out:** HMMER search result against dbCAN-sub HMMdb, including a column with CAZyme substrates extracted from *fam-substrate-mapping-08012023.tsv*.

**diamond.out**: DIAMOND search result against the CAZy annotated protein sequences (*CAZyDB.07262023.fa*).

**dtemp.out**: temporary file.

**hmmer.out**: HMMER search result against dbCAN HMMdb.

**overview.txt**: summary of CAZyme annotation from three methods in TSV format. An example row has the following columns:

13. Gene_ID: *BML_MG_01761*
14. EC#: *2.4.99.-:5*
15. dbCAN: *GT112(19-370)*
16. dbCAN_sub: *GT112_e0*
17. DIAMOND: *GT112*
18. #ofTools: *3*

*Explanation: the protein BML_MG_01761 is annotated by 3 tools to be a CAZyme: (1) GT112 (CAZy defined family GT112) by HMMER vs dbCAN HMMdb with a domain range from aa position 19 to 370, (2) GT112_e0 (eCAMI defined subfamily e0; e indicates it is from eCAMI not CAZy) by HMMER vs dbCAN-sub HMMdb (derived from eCAMI subfamilies), and (3) GT112 by DIAMOND vs CAZy annotated protein sequences. The second column 2.4.99.-:5 is extracted from eCAMI, meaning that the eCAMI subfamily GT112_e0 contains 5 member proteins which have an EC 2.4.99.- according to CAZy. In most cases, the 3 tools will have the same CAZyme family assignment. When they give different assignment. We recommend a preference order: dbCAN > eCAMI/dbCAN-sub > DIAMOND. See our dbCAN2 paper^1^, dbCAN3 paper^2^, and eCAMI^3^ for more details.*

*Note: If users invoked the --use_signalP parameter when running run_dbcan, there will be an additional column called signal in the overview.txt.*

**stp.out**: HMMER search result against the MiST^4^ compiled signal transduction protein HMMs from Pfam.

**tf-1.out**: HMMER search result against the DBD^5^ compiled transcription factor HMMs from Pfam ^6^.

**tf-2.out**: HMMER search result against the DBD compiled transcription factor HMMs from Superfamily ^7^.

**tp.out**: DIAMOND search result against the TCDB ^8^ annotated protein sequences.

**substrate.out**: summary of substrate prediction results for CGCs in TSV format from two approaches^2^ (dbCAN-PUL blast search and dbCAN-sub majority voting). An example row has the following columns:

15. CGC_ID: *contig_10778|CGC2*
16. Best hit PUL_ID in dbCAN-PUL: *PUL0400*
17. Substrate of the hit PUL: *alginate*
18. Sum of bitscores for homologous gene pairs between CGC and PUL: *851.0*
19. Types of homologous gene pairs: *CAZyme-CAZyme;CAZyme-CAZyme;CAZyme-CAZyme;CAZyme-CAZyme*
20. Substrate predicted by majority voting of CAZymes in CGC: *alginate*
21. Voting score: *2.0*

*Explanation: The CGC2 of contig_10778 has its best hit PUL0400 (from PUL_blast.out) with alginate as substrate (from dbCAN-PUL_12-12-2023.xlsx). Four signature genes are matched between contig_10778|CGC2 and PUL0400 (from PUL_blast.out): all the four are CAZymes. The sum of blast bitscores of the 4 homologous pairs (CAZyme-CAZyme;CAZyme-CAZyme;CAZyme-CAZyme;CAZyme-CAZyme) is 851.0. Hence, the substrate of contig_10778|CGC2 is predicted to be alginate according to dbCAN-PUL blast search. The last two columns are based on the dbCAN-sub result (dbcan-sub.hmm.out), according to which two CAZymes in contig_10778|CGC2 are predicted to have alginate substrate. The voting score is thus 2.0, so that according to the majority voting rule, contig_10778|CGC2 is predicted to have an alginate substrate.*

*Note: for many CGCs, only one of the two approaches produces substrate prediction. In some cases, the two approaches produce different substrate assignments. We recommend a preference order: dbCAN-PUL blast search > dbCAN-sub majority voting. See our dbCAN3 paper*^2^ *for more details.*

**synteny.pdf**: a folder with syntenic block alignment plots between all CGCs and PULs.

**uniInput**: renamed Fasta file from input protein sequence file.

**Module 3: Read mapping (Fig. 3) to calculate abundance for CAZyme families, subfamilies, CGCs, and substrates**

In the Priest2023 dataset, we have both HiFi long reads for MG sequencing and Illumina short reads for MT sequencing. We have mapped MG reads to contigs, and MT reads to CDS of all Prokka predicted proteins.

**P9| Illumina MT short read mapping to all CDS of each sample (TIMING ~36 min)**

*$ bwa index prokka_SRF_MG/SRF_MG.ffn*

*$ bwa index prokka_BML_MG/BML_MG.ffn*

*$ bwa mem -t 32 -o samfiles/SRF_MT.CDS.sam prokka_SRF_MG/SRF_MG.ffn SRF_MT_1.clean_val_1.fq.gz SRF_MT_2.clean_val_2.fq.gz*

*$ bwa mem -t 32 -o samfiles/BML_MT.CDS.sam prokka_BML_MG/BML_MG.ffn BML_MT_1.clean_val_1.fq.gz BML_MT_2.clean_val_2.fq.gz*

Illunima reads are mapped to the ffn files from Prokka.

**P10| HiFi MG long read mapping to all contigs of each sample (TIMING ~20min)**

*$ minimap2 -a -x map-hifi -MD -t 32 -o samfiles/BML_MG.sam flye_BML_MG/assembly.fasta BML_MG.clean.fq.gz*

*$ minimap2 -a -x map-hifi -MD -t 32 -o samfiles/SRF_MG.sam flye_SRF_MG/assembly.fasta SRF_MG.clean.fq.gz*

**P11| Sort SAM files by coordinates (TIMING ~8min)**

*$ cd samfiles*

*$ samtools sort -@ 32 -o SRF_MT.CDS.bam SRF_MT.CDS.sam*

*$ samtools sort -@ 32 -o BML_MT.CDS.bam BML_MT.CDS.sam*

*$ samtools sort -@ 32 -o SRF_MG.bam SRF_MG.sam*

*$ samtools sort -@ 32 -o BML_MG.bam BML_MG.sam*

*$ rm -rf *sam*

*$ cd ..*

**P12| Read count calculation for all proteins of each sample using Bedtools (TIMING ~6min)**

Calculate HiFi MG read abundance for proteins in the two samples:

*$ mkdir SRF_MG_abund && cd SRF_MG_abund*

*$ seqkit fx2tab -l -n -i ../flye_SRF_MG/assembly.fasta | awk '{print $1"\t"$2}' > SRF_MG.length*

*$ grep -v "^#" ../prokka_SRF_MG/SRF_MG.gff | awk '{if($3=="CDS") print $0}' > SRF_MG.gff*

*$ bedtools coverage -g SRF_MG.length -sorted -a SRF_MG.gff -counts -b ../samfiles/SRF_MG.bam | awk -F"\t" '{s=index($(NF-1),"=");e=index($(NF-1),";");geneID=substr($(NF-1),s+1,e-s-1); print geneID"\t"0"\t"$5-$4+1"\t"$NF}' > SRF_MG.depth.txt*

*$ cd ..*

*$ mkdir BML_MG_abund && cd BML_MG_abund*

*$ seqkit fx2tab -l -n -i ../flye_BML_MG/assembly.fasta | awk '{print $1"\t"$2}' > BML_MG.length*

*$ grep -v "^#" ../prokka_BML_MG/BML_MG.gff | awk '{if($3=="CDS") print $0}' > BML_MG.gff*

*$ bedtools coverage -g BML_MG.length -sorted -a BML_MG.gff -counts -b ../samfiles/BML_MG.bam | awk -F"\t" '{s=index($(NF-1),"=");e=index($(NF-1),";");geneID=substr($(NF-1),s+1,e-s-1); print geneID"\t"0"\t"$5-$4+1"\t"$NF}' > BML_MG.depth.txt*

*$ cd ..*

Calculate Illumina MT read abundance for proteins in the two samples:

*$ mkdir BML_MT_abund && cd BML_MT_abund*

*$ seqkit fx2tab -l -n -i ../prokka_BML_MG/BML_MG.ffn | awk '{print $1"\t"$2}' > BML_MT.length*

*$ seqkit fx2tab -l -n -i ../prokka_BML_MG/BML_MG.ffn | awk '{print $1"\t"0"\t"$2}' > BML_MT.bed*

*$ bedtools coverage -g BML_MT.length -sorted -a BML_MT.bed -counts -b ../samfiles/BML_MT.CDS.bam > BML_MT.depth.txt*

*$ cd ..*

*$ mkdir SRF_MT_abund && cd SRF_MT_abund*

*$ seqkit fx2tab -l -n -i ../prokka_SRF_MG/SRF_MG.ffn | awk '{print $1"\t"$2}' > SRF_MT.length*

*$ seqkit fx2tab -l -n -i ../prokka_SRF_MG/SRF_MG.ffn | awk '{print $1"\t"0"\t"$2}' > SRF_MT.bed*

*$ bedtools coverage -g SRF_MT.length -sorted -a SRF_MT.bed -counts -b ../samfiles/SRF_MT.CDS.bam > SRF_MT.depth.txt*

*$ cd ..*

Read counts are saved in *depth.txt* files of each sample.

**P13| Read count calculation for a given region of contigs using Samtools (TIMING ~2min)**

*$ cd SRF_MG_abund*

*$ samtools index ../samfiles/SRF_MG.bam*

*$ samtools depth -r contig_34043:9828-35768 ../samfiles/SRF_MG.bam > SRF_MG.cgc.depth.txt*

*$ cd ..*

The parameter *-r contig_34043:9828-35768* specifies a region in a contig. For any CGC, its positional range can be found in the file cgc_standard.out produced by run_dbcan (**Box 6**). The *depth.txt* files contain the raw read counts for the specified region.

**P14| dbcan_utils to calculate the abundance of CAZyme families, subfamilies, CGCs, and substrates (TIMING ~1min)**

As the Priest2023 dataset has both MG and MT data, for each sample we will have two abundance folders, e.g., *BML_MG_abund, BML_MT_abund* for BML sample, and *SRF_MG_abund,* *SRF_MT_abund* for SRF sample.

*$ cd BML_MT_abund*

*$ dbcan_utils fam_abund -bt BML_MT.depth.txt -i ../BML_MG.dbCAN -a TPM*

*$ dbcan_utils fam_substrate_abund -bt BML_MT.depth.txt -i ../BML_MG.dbCAN -a TPM*

*$ dbcan_utils CGC_abund -bt BML_MT.depth.txt -i ../BML_MG.dbCAN -a TPM*

*$ dbcan_utils CGC_substrate_abund -bt BML_MT.depth.txt -i ../BML_MG.dbCAN -a TPM*

*$ cd .. && cd SRF_MG_abund*

*$ dbcan_utils fam_abund -bt SRF_MG.depth.txt -i ../SRF_MG.dbCAN -a TPM*

*$ dbcan_utils fam_substrate_abund -bt SRF_MG.depth.txt -i ../SRF_MG.dbCAN -a TPM*

*$ dbcan_utils CGC_abund -bt SRF_MG.depth.txt -i ../SRF_MG.dbCAN -a TPM*

*$ dbcan_utils CGC_substrate_abund -bt SRF_MG.depth.txt -i ../SRF_MG.dbCAN -a TPM*

*$ cd .. && cd BML_MG_abund*

*$ dbcan_utils fam_abund -bt BML_MG.depth.txt -i ../BML_MG.dbCAN -a TPM*

*$ dbcan_utils fam_substrate_abund -bt BML_MG.depth.txt -i ../BML_MG.dbCAN -a TPM*

*$ dbcan_utils CGC_abund -bt BML_MG.depth.txt -i ../BML_MG.dbCAN -a TPM*

*$ dbcan_utils CGC_substrate_abund -bt BML_MG.depth.txt -i ../BML_MG.dbCAN -a TPM*

*$ cd .. && cd SRF_MT_abund*

*$ dbcan_utils fam_abund -bt SRF_MT.depth.txt -i ../SRF_MG.dbCAN -a TPM*

*$ dbcan_utils fam_substrate_abund -bt SRF_MT.depth.txt -i ../SRF_MG.dbCAN -a TPM*

*$ dbcan_utils CGC_abund -bt SRF_MT.depth.txt -i ../SRF_MG.dbCAN -a TPM*

*$ dbcan_utils CGC_substrate_abund -bt SRF_MT.depth.txt -i ../SRF_MG.dbCAN -a TPM*

*cd ..*

We developed a set of Python scripts as dbcan_utils (included in the run_dbcan package) to take the raw read counts for all genes as input and output the normalized abundances (**Box 7**) of CAZyme families, subfamilies, CGCs, and substrates (**Fig. 4**). The parameter *-a TPM* can also be two other metrics: RPM, or RPKM ^11^.

RPKM = # of mapped reads to a gene G / [(total # of mapped reads to all genes /10^6^) x (gene G length/1000)].

RPM = # of mapped reads to a gene G / (total # of mapped reads to all genes/10^6^).

TPM = [# of mapped reads to a gene G / (gene G length/1000)] / sum [# of mapped reads to each gene / (the gene length/1000)].

**Box 7| Example output of dbcan_utils**

As an example, the *SRF_MG_abund* folder (<https://bcb.unl.edu/dbCAN_tutorial/dataset3-Priest2023/SRF_MG_abund/>) has 7 TSV files:

```
-rw-rw-r--  1 jinfang jinfang 103K Jan  2 07:26 CGC_abund.out
-rw-rw-r--  1 jinfang jinfang 1.3K Jan  2 07:26 CGC_substrate_majority_voting.out
-rw-rw-r--  1 jinfang jinfang 4.4K Jan  2 07:26 CGC_substrate_PUL_homology.out
-rw-rw-r--  1 jinfang jinfang 2.2K Jan  2 07:26 EC_abund.out
-rw-rw-r--  1 jinfang jinfang 3.0K Jan  2 07:26 fam_abund.out
-rw-rw-r--  1 jinfang jinfang  13K Jan  2 07:26 fam_substrate_abund.out
-rw-rw-r--  1 jinfang jinfang  16K Jan  2 07:26 subfam_abund.out
```

Explanation of columns in these TSV files is as follows:

**fam_abund.out**: CAZy family (from HMMER vs dbCAN HMMdb), sum of TPM, # of CAZymes in the family

**subfam_abund.out:** eCAMI subfamily (from HMMER vs dbCAN-sub HMMdb), sum of TPM, # of CAZymes in the subfamily

**EC_abund.out:** EC number (extracted from dbCAN-sub subfamily), sum of TPM, # of CAZymes with the EC

**fam_substrate_abund.out:** Substrate (from HMMER vs dbCAN-sub HMMdb), sum of TPM (all CAZymes in this substrate group), GeneID (all CAZyme IDs in this substrate group)

**CGC_abund.out:** CGC_ID (e.g., contig_34043|CGC1), mean of TPM (all genes in the CGC), Seq_IDs (IDs of all genes in the CGC), TPM (of all genes in the CGC), Families (CAZyme family or other signature gene type of all genes in the CGC)

**CGC_substrate_PUL_homology.out:** Substrate (from dbCAN-PUL blast search), sum of TPM, CGC_IDs (all CGCs predicted to have the substrate from dbCAN-PUL blast search), TPM (of CGCs in this substrate group)

**CGC_substrate_majority_voting.out:** Substrate (from dbCAN-sub majority voting), sum of TPM, CGC_IDs (all CGCs predicted to have the substrate from dbCAN-sub majority voting), TPM (of CGCs in this substrate group)

**Module 4: dbcan_plot for data visualization (Fig. 3) of abundances of CAZymes, CGCs, and substrates (TIMING variable)**

**CRITICAL STEP**

To visualize the CAZyme annotation result, we provide a set of Python scripts as dbcan_plot to make publication quality plots with the dbcan_utils results as the input. The dbcan_plot scripts are included in the run_dbcan package. Once the plots are made in PDF format, they can be transferred to users’ Windows or Mac computers for visualization.

Seven data folders will be needed as the input for dbcan_plot: (i) four abundance folders BML_MG_abund, BML_MT_abund, SRF_MG_abund and SRF_MT_abund, (ii) two CAZyme annotation folders BML_MG.dbCAN and SRF_MG.dbCAN, and (iii) the dbCAN-PUL folder (under the db folder, released from dbCAN-PUL.tar.gz).

**P15| Heatmap for CAZyme substrate abundance across samples (Fig. S5D) (TIMING 1min)**

*$ dbcan_plot heatmap_plot --show_abund --top 20 --samples BML_MG,BML_MT,SRF_MG,SRF_MT -i BML_MG_abund/fam_substrate_abund.out,BML_MT_abund/fam_substrate_abund.out,SRF_MG_abund/fam_substrate_abund.out,SRF_MT_abund/fam_substrate_abund.out*

Here we plot the top 20 substrates in the two samples. The input files are the four CAZyme substrate abundance files in both MG and MT data of each sample calculated based on dbCAN-sub result. The default heatmap is ranked by substrate abundances. To rank the heatmap according to abundance profile using the using the function clustermap of seaborn package, users can invoke the --cluster_map parameter.

**P16| Barplot for CAZyme family/subfamily/EC abundance across samples (Fig. S5A-C) (TIMING 1min)**

*$ dbcan_plot bar_plot --samples BML_MG,BML_MT,SRF_MG,SRF_MT --vertical_bar --top 20 -i BML_MG_abund/fam_abund.out,BML_MT_abund/fam_abund.out,SRF_MG_abund/fam_abund.out,SRF_MT_abund/fam_abund.out*

*$ dbcan_plot bar_plot --samples BML_MG,BML_MT,SRF_MG,SRF_MT --vertical_bar --top 20 -i BML_MG_abund/subfam_abund.out,BML_MT_abund/subfam_abund.out,SRF_MG_abund/subfam_abund.out,SRF_MT_abund/subfam_abund.out*

*$ dbcan_plot bar_plot --samples BML_MG,BML_MT,SRF_MG,SRF_MT --vertical_bar --top 20 -i BML_MG_abund/EC_abund.out,BML_MT_abund/EC_abund.out,SRF_MG_abund/EC_abund.out,SRF_MT_abund/EC_abund.out*

**P17| Synteny plot between a CGC and its best PUL hit with read mapping coverage to CGC (Fig. S5E) (TIMING 1min)**

*$ dbcan_plot CGC_synteny_coverage_plot -i SRF_MG.dbCAN --cgcid 'contig_34043|CGC1' --readscount SRF_MG_abund/SRF_MG.cgc.depth.txt*

The SRF_MG.dbCAN folder contains the PUL_blast.out file. Using this file, the cgc_standard.out file, and the best PUL’s gff file in dbCAN-PUL.tar.gz, the CGC_synteny_plot method will create the CGC-PUL synteny plot. The –cgcid parameter is required to specify which CGC to be plotted (contig_34043|CGC1 in this example). The *SRF_MG.cgc.depth.txt* file is used to plot the read mapping coverage.

If users only want to plot the CGC structure:

*$ dbcan_plot CGC_plot -i SRF_MG.dbCAN --cgcid 'contig_34043|CGC1'*

If users only want to plot the CGC structure plus the read mapping coverage:

*$ dbcan_plot CGC_coverage_plot -i SRF_MG.dbCAN --cgcid 'contig_34043|CGC1' --readscount SRF_MG_abund/SRF_MG.cgc.depth.txt*

If users only want to plot the synteny between the CGC and PUL:

*$ dbcan_plot CGC_synteny_plot -i SRF_MG.dbCAN --cgcid 'contig_34043|CGC1'*

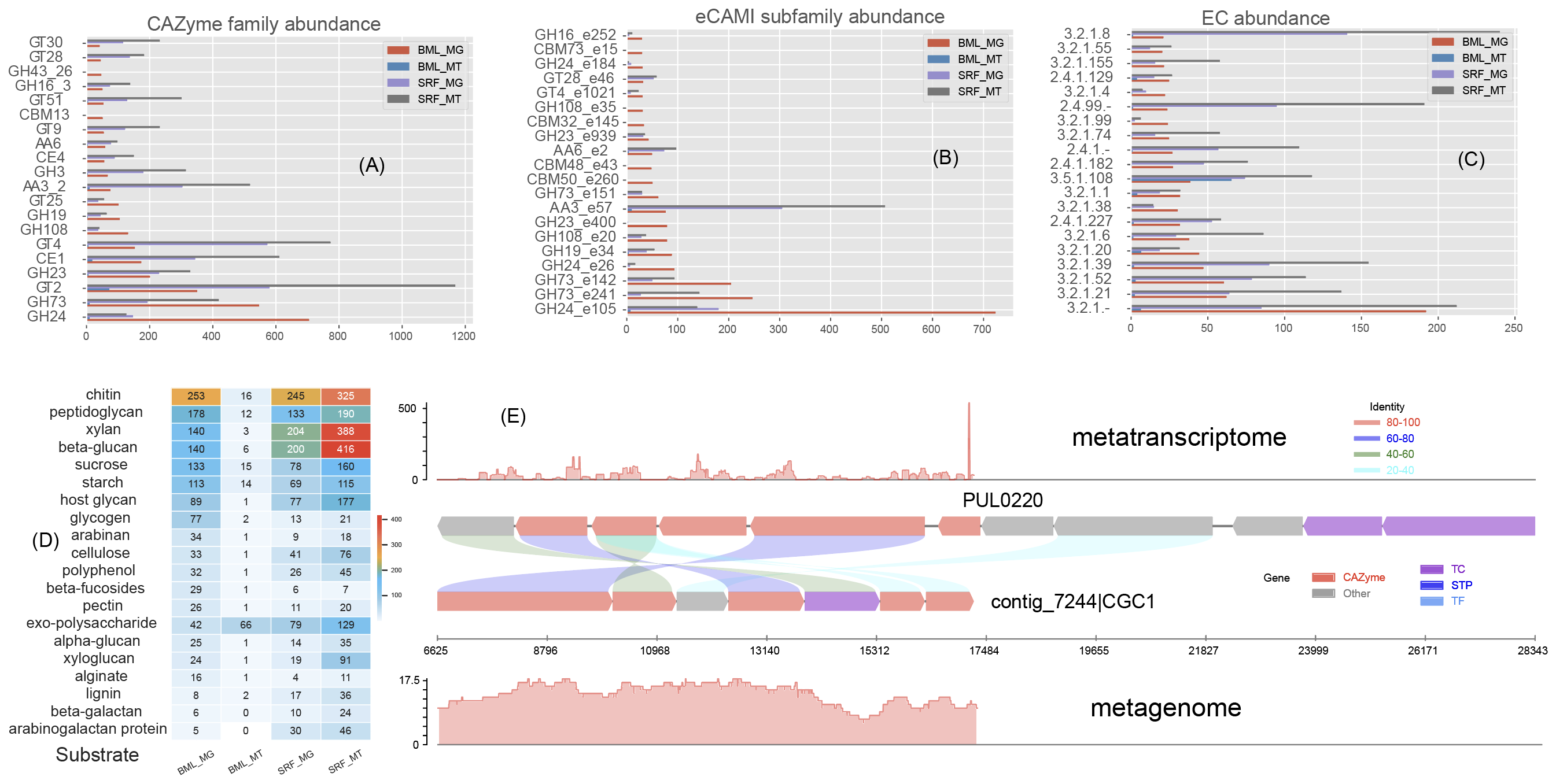

***Fig. S5.*** ***Data visualization of abundances of CAZymes, CGCs, and substrates****. (****A****) Barplot of 20 CAZyme family abundance in both MG and MT data of the two samples (see Table S3) calculated based on dbCAN search results. (****B****) Barplot of 20 eCAMI subfamily abundance calculated based on dbCAN-sub search results. (****C****) Barplot of 20 CAZyme EC abundance calculated based on dbCAN-sub search results. (****D****) Heatmap of CAZyme substrate abundance calculated based on dbCAN-sub search results and substrate mapping. Abundance values are TPM. (****E****) Synteny plot between an example CGC (CGC1 of contig_7244 from the S25_SRF sample) and its best PUL hit (PUL0220 from dbCAN-PUL, with experimentally verified substrate beta-glucan:* [*https://bcb.unl.edu/dbCAN_tutorial/dataset3-Priest2023/SRF_MG.dbCAN/substrate.out*](https://bcb.unl.edu/dbCAN_tutorial/dataset3-Priest2023/SRF_MG.dbCAN/substrate.out)*) with MT and MG read mapping coverage plots (y-axis is the read depth) shown on the top and in the bottom.*
